# Personalized Therapy Design for Liquid Tumors via Optimal Control Theory

**DOI:** 10.1101/662858

**Authors:** Fabrizio Angaroni, Alex Graudenzi, Marco Rossignolo, Davide Maspero, Tommaso Calarco, Rocco Piazza, Simone Montangero, Marco Antoniotti

## Abstract

One of the key challenges in current cancer research is the development of reliable methods for the definition of personalized therapeutic strategies, based on increasingly available experimental data on single patients. To this end, methods from control theory can be effectively employed on patient-specific pharmacokinetic and pharmacodynamic models to generate robust data-driven experimental hypotheses.

Here we introduce the Control Theory for Therapy Design (CT4TD) theoretical framework for the generation of optimized personalized therapeutic strategies in cancer patients, based on optimal control theory and population dynamics modeling. The CT4TD framework can help clinicians in designing patient-specific therapeutic regimens, with the specific goal of optimizing the efficacy of the cure while reducing the costs, especially in terms of toxicity and adverse effects. CT4TD can be used at the time of the diagnosis in order to set optimized personalized therapies to reach selected target drug concentrations. Furthermore, if longitudinal data on patients under treatment are available, our approach introduces the possibility of adjusting the therapy with the explicit goal of minimizing the tumor burden measured in each case.

As a case study, we present the application of CT4TD to Imatinib administration in Chronic Myeloid Leukemia, in which we show that the optimized therapeutic strategies are extremely diversified among patients, and display improvements with respect to the currently employed regimes. Interestingly, we prove that much of the variance in therapeutic response observed among patients is due to the individual differences in pharmacokinetics, rather than in pharmacodynamics.

## Introduction

The increasing availability of reliable experimental data on cancer patients and the concurrent decreasing costs of computational power are fueling the development of algorithmic strategies for the automated generation of experimental hypotheses in cancer research. This is particularly relevant in the sphere of precision and personalized medicine, as efficient methods are urgently needed to make sense of available data and support clinicians in delivering patient-specific therapeutic strategies [1]. To this end, methods borrowed from optimal control theory [2, 3, 4, 5] can be employed in combination with efficient techniques for data analysis [6, 7, 8, 9], to produce accurate predictive model of the clinical outcome of a given therapy in single cancer patients.

Here, we introduce a theoretical framework named CT4TD (Control Theory for Therapy Design), which employs the RedCRAB optimal control algorithm [10], on patient-specific pharmacokinetics and pharmacodynamics (PK/PD) models [11], with the goal of delivering an optimized drug administration schedule (see Figure 1 for a schematic representation of the framework).

**Figure 1:**
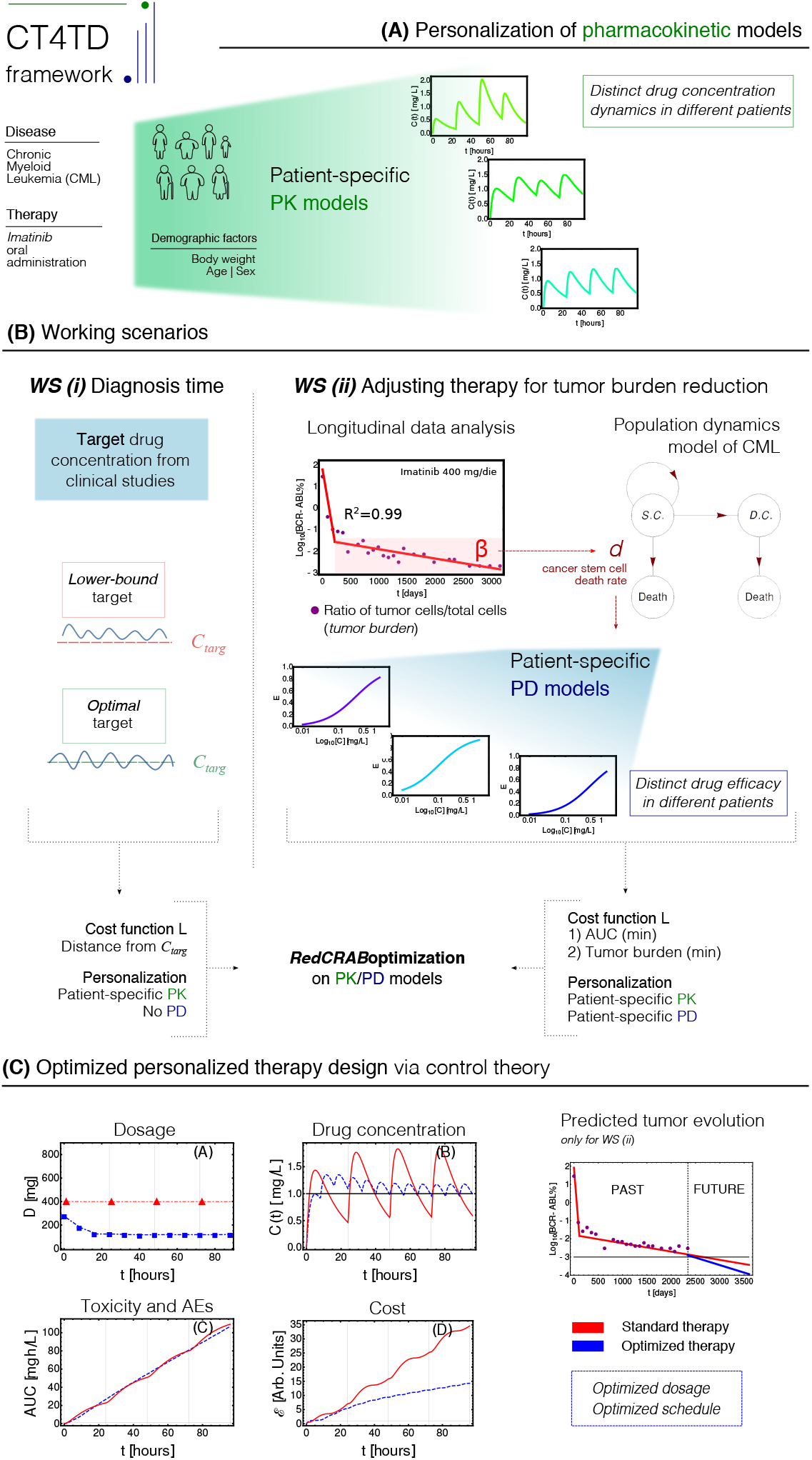
CT4TD pipeline. **(A)** The CT4TD framework employs demographic factors such as body weight, age and sex to define patient-specific parameters of the pharmacokinetic models. We here focus on the case of Imatinib administration in Chronic Myeloid Leukemia (CML). **(B)** CT4TD manages two working scenarios: (*i*) at time of diagnosis, CT4TD can be used to reach given optimal/lower-bound drug concentration targets, e.g., from clinical studies; (*ii*) when longitudinal data on tumor burden variation under standard therapy are available, CT4TD fits the data points with a hierarchical population dynamics model of CML, and this allows to estimate patient-specific pharmacodynamics (PD) parameters, based on the observed cancer cell death rate. In both scenarios, optimization on pharmacokinetics/pharmacodynamics (PK/PD) models is performed via RedCRAB, on distinct cost functions, aimed at: either being close to given target concentrations (and strictly larger in the lower-bound case) – WS (*i*); minimizing the Area Under the Curve (AUC) *and* the tumor burden -Working Scenario (WS) (*ii*). **(C)** Optimized personalized dosage and schedule are returned, allowing to measure in silico the differences with respect to standard administration, in terms of dosage, drug concentration, cost and AUC. WS (*ii*) allows to predict the tumor burden evolution in case of an optimized therapy.

In brief, PK models describe the temporal dynamics of the concentration of a given drug in a certain tissue or organ, whereas PD models depict the efficacy of the drug with respect to distinct concentration values. CT4TD framework first defines patient-specific PK models based on demographic factors, such as, e.g., age, sex and body weight. Such models are employed to automatically identify optimized therapy dosages and/or schedules to reach given target drug concentrations, as those commonly used in clinical practice, also by respecting any desired constraint such as, e.g., the maximum allowed number of doses per day or the maximum dosage. In this way, our framework can guide clinicians in the setting of optimized regimes at diagnosis, allowing for an either more or less aggressive tuning; this approach mimics “steady state” optimization commonly used in pharmacological studies.

Furthermore, when longitudinal experimental data on tumor burden – e.g., the fraction of tumor cells on the total, in liquid tumors – are available for patients under standard treatment, CT4TD allows to determine optimized personalized strategies to be used in order to minimize or even eradicating the cancer cell subpopulation. In fact, with the CT4TD framework it is possible to fit experimental data with a hierarchical population dynamics model, which describes the temporal evolution of cancer subpopulations in a given tumor [6, 12, 13, 14]. Such model allows to measure the impact of a given therapy over the tumor’s ability to expand and develop and, accordingly, to estimate patient-specific PD models from clinical data, which are then employed to design optimized therapeutic regimes aimed at reducing the tumor burden.

Therefore, CT4TD can support clinicians in designing personalized therapies both at diagnosis and when longitudinal data on disease progression have become available. In all scenarios, with our approach it is possible to compare the actual therapeutic regime with the optimized one, showing the qualitative and quantitative improvements in terms of efficacy, toxicity and overall costs.

The CT4TD theoretical framework is general and applicable to any kind of disease, as long as PK/PD models can be retrieved or estimated. Yet, liquid tumours allow to safely adopt several simplifications and define simple and reliable models of population dynamics, avoiding possible complications due to the spatial and morphological properties of solid tumors [15].

For this reason, in this work we apply the CT4TD to the specific case of Imatinib administration in patients with Chronic Myeloid Leukemia (CML), and we show the advantages of employing our automated and data-driven framework in terms of increased efficacy of the therapy and reduction of the overall costs and toxicity. In particular, we here present the results of the application of the CT4TD framework in two ideally subsequent scenarios.

In the first case, CT4TD is used to identify the best therapeutic regime to reach selected target drug concentrations, as those commonly used in the clinic (e.g., [16, 17, 18]). This scenario provides indications which can be employed by clinicians at the time of the diagnosis. Importantly, the inclusion of demographic factors within the PK models [19] allows to define personalized drug schedules that are remarkably different from standard practice. A robustness analysis on intra- and inter-patient variability shown in the Supplementary Information proves the safety of the hypotheses generated with our approach, especially with respect to possible technical or measurement errors.

In the second case, we employ longitudinal data on tumor burden of a selected cohort of CML patients under standard treatment, in order to retrieve personalized PD models and, accordingly, to identify an “adjusted” therapy that is most effective in minimizing cancer subpopulation, once the major molecular response has been observed. In both cases the results allow to explicitly evaluate the advantages in costs and improved efficacy with respect to standard therapies.

## Background

### Pharmacokinetic and pharmacodynamic models

Pharmacokinetic (PK) models [11] are mathematical models that describe the temporal evolution of the concentration of a substance in a certain tissue of the body. Commonly used techniques to study such processes are the so-called *compartmental* models i.e., dynamical models based on the law of conservation of mass, and which assume that the body is composed by a certain number of macroscopic coupled subsystem, namely compartments. Such models assume an instantaneous mixing of the drug in a compartment and a perfect transport among them, and are usually defined via systems of differential equations. The solution of such systems provide predictions about the variation of drug concentration in time, in a certain tissue. A limitation of PK models is the employment of coarse-grained oversimplifications, which require ad-hoc assumptions and are valid only for sufficiently long timescales.

Pharmacodynamic (PD) models [11, 20] study the relationship between the concentration of a drug and the resulting effect, in terms of efficacy and possible adverse effects (AEs). The effects of a certain substance are estimated by modeling relevant biochemical reactions, usually by exploiting the law of mass action (see, e.g., [21]). One of the major limitations of PD models is that it is usually impossible to have all the measurements necessary to determine the kinetic constants of the involved chemical reactions. For this reason, the efficacy of a drug is usually estimated with statistical methods and target concentrations are defined with respect to some arbitrary criteria [17, 22, 23, 24, 18].

PK/PD models are increasingly used to define new drug dosage guidelines and protocols [17]. Nonetheless, standard approaches to this end are affected by several major issues. Usually the optimal dose is identified in phase I dose escalating clinical trials. Often these trials suffer from possible idiosyncrasies of the study, from the presence of unknown confounding factors and from the often limited sample size. Another problem is that the recommended dosage is often defined as optimal for an ideal – and non existing – *average* patient, because the efficacy is only defined statistically. As a consequence, the same drug dosage/schedule might be either insufficient or exceeding for different patients. In the former case, this might lead to a non-optimal clinical outcome, in terms of lower efficacy of the treatment, whereas in the latter case an excess of drug may raise the probability of AEs, as well as the economic cost of the therapy, an aspect that is particular relevant for oncological therapies [25, 26, 27, 28, 29].

Therefore, effective strategies for the identification of optimized personalized therapy schedules are urgently needed, in order to possibly reduce the amount of drug – and minimize the probability of related adverse effects –, while providing the same or an even better efficacy – i.e., clinical outcome –, with respect to the standard administration schedule. As a side effect, an optimized personalized schedule would also deliver a minimal economic cost, i.e., more patients will be able to afford its costs.

In this respect, CT4TD allows to: (*i*) define patient-specific PK models that depend on a number of demographic factors and biological covariates, such as age, sex, ethnicity, and body weight, as proposed in [19]; (*ii*) estimate personalized PD models from longitudinal experimental data on tumor burden (if available). This allows to identify personalized therapeutic strategies, which explicitly account for the expected differences in PK and PD, due to the physiological heterogeneity of the individuals.

### Applications of optimal control theory in medicine

Control theory is an interdisciplinary field bridging engineering and mathematics, whose main objective is to define an opportune control function that modifies the state of a given dynamical system in order to perform a specific task, while minimizing the cost and maximizing the performance [2, 4, 5] (see Materials and Methods for a technical description).

Two main classes of controls exist: (*i*) open-loop control, and (*ii*) closed-loop (*feedback*) control. In the former case, the set and sequence of control actions is chosen a priori, by exploiting theoretical study on the models. In this case, the input is independent with respect to the output (e.g., possible measurements on the system) [4]. Closed-loop control, instead, introduces in the procedure one or more feedback loops, which are able to quantify the real response of the system to variations of the control functions, and adjust them according to the differences recorded between the theoretical and real behaviors of the system [5].

There are several examples of successful applications of control theory in pharmacology [3, 30]. In this respect, the final goal is to determine the optimal set of therapeutic choices – e.g., dosages and schedules – to obtained a desired efficacy, while minimizing the overall costs. Closed-loop controls are extremely effective in achieving this goal and have been often implemented in real-world health-care [31, 32, 30, 33, 34, 35]. Nonetheless, technological problems, such the absence of real-time measurements, as well as possible problems in titrating drugs to the right concentration, are still limiting real-life applications [3, 36]. Despite the lower complexity, open-loop controls are still a viable option, mostly because of the applicability in a wide range of real-world scenarios for which, for instance, real-time measurements and/or therapy adjustments are unfeasible. Moreover, open-loop controls have proven to identify more effective drug concentration in therapeutic ranges than standard clinical practice [37, 38, 39, 40, 41].

However, many approaches in both categories are based on limiting assumptions. Certain techniques, for instance, assume continuous – yet unrealistic – drug infusion procedures (e.g., [42]). Some methods rely on often speculative mathematical models, which cannot be evaluated due to the lack of opportune experimental data [41]. Some further methods are based on Pontryagin Maximum principle [4] and/or they assume a quadratic cost in order to ensure the optimality of the protocol, whereas such assumptions are not realistic in real-world scenarios.

The CT4TD framework aims at improving the current state-of-the-art, by solving an open-loop control problem on PK/PD models via RedCRAB (**Re**mote **d**ressed **C**hopped **Ra**ndom **B**asis) [43, 44, 10], a gradient-free framework for optimization and control. Extensions to the closed-loop case are underway.

### Mathematical modeling of cell population dynamics

Many healthy and aberrant biological tissues are characterized by a hierarchical organization, constituted by an ordered sequence of discrete maturation states, as driven by differentiation processes. In this respect, a number of mathematical models have been proposed to study the cell population dynamics, both in healthy systems [45] and in cancer [6, 7, 12, 8, 46, 13, 14].

In such models, cells are divided in *n* non-intersecting compartments, with every ensemble representing a certain stage of cell differentiation. The time ordering of the differentiation stage defines an explicit hierarchy among such ensembles. Accordingly, a *lineage* is defined as a collection of compartments that fully describe all the stages of differentiation of cells within a certain tissue (see Supplementary Figure S7).

Various approaches are employed to model the dynamics of such systems such as, e.g., ordinary differential equations (ODEs), discrete-time and continuous-time Markov chains, master equations, etc. (see [46] for a recent review). Each strategy displays a specific trade-off in terms of expressivity and computational complexity. For instance, ODEs are very convenient from the computational perspective, but they are not suitable in certain cases, e.g., when representing low numbers of cells. Conversely, probabilistic models allow for a richer representation of the system, yet at the cost of a higher computational burden and mathematical complexity.

For sake of simplicity, the CT4TD framework employs a ODEs hierarchical model of cell population dynamics to fit longitudinal data on tumor burden [6, 7, 12, 8, 46, 13, 14]. On the one hand, this model provides a description of cell population dynamics in time for any given patient. On the other hand, it allows to estimate the efficacy of the therapy in each patient, on the basis of the observed cancer subpopulation decay, which is then used to estimate patient-specific PD models (see Materials and Methods for further details).

## Results

### Imatinib administration in Chronic Myeloid Leukemia – CML

We here show the application of CT4TD to the specific case of Imatinib mesylate administration in patients with CML. The final goal is to determine the drug optimized dosage and schedule in two distinct scenarios: (*i*) in the first case the goal is to optimize personalized therapeutic strategies to reach given target concentrations, as those commonly used in clinical protocols, and by assuming to be at diagnosis time; (*ii*) in the second case, we employ the population dynamics models, as retrieved by fitting longitudinal data on single patients under standard treatment, to deliver patient-specific therapies that are most effective in reducing/eradicating the tumor subpopulation after the major molecular response, on the basis of PK/PD personalized models.

Imatinib is an inhibitor of the BCR-ABL tyrosine kinase, which is known to bind to the inactive form of BCR-ABL at nanomolar concentration, competing with the ATP for its binding pocket and hindering the switch of the fusion kinase to the active form, therefore impairing the catalytic activity of the enzyme [47]. The therapy is in most cases long-life [6, 7, 48, 8, 9] and the treatment is expensive (≈ 30,000 US$ per year) [49]. Therefore, the impact of an optimized and personalized administration would be two-fold: on the one hand, it could be effective in optimizing the performance, while reducing the toxicity and minimizing the adverse effects for long-term therapies [50, 51, 52]; on the other hand, it could help in reducing the overall economical costs, which currently limit the access to therapy, hence making long-term health care more sustainable [27, 25].

### Datasets

We applied the CT4TD framework to a longitudinal dataset from [6], in which 29 CML patients have been monitored with a peripheral blood draw taken every 90 days, from the time of diagnosis up to a maximum time of about 3500 days (average time ≈ 2659 ± 938 days). For these patients, the administration schedule has been 400 mg Imatinib/day for the whole considered period.

In particular the fraction of cancer cells in blood – that will be referred to as *tumour burden* from now on – can be reliably estimated by analyzing the expression level of the fusion gene BCR-ABL, thus providing an easy way to monitor the disease progression, as well the response to therapy. As BCR-ABL transcript is solely expressed by the leukemic cells, its measurement by mean of quantitative PCR (Q-PCR) is considered one of the most sensitive and specific techniques to indirectly assess the tumor burden, and is the standard de facto for monitoring minimal residual disease in CML.

More in detail, we selected a subset of the dataset provided in [6], by removing all the patients that displayed too few data points (i.e., < 3), or that were characterized by resistant mutations, i.e., specific DNA alterations that render the therapy via Imatinib ineffective, usually due to steric impediments [53]. In such cases, it is common practice to employ an alternative therapy, based either on Dasatinib, Nilotinib, Ponatinib or Bosutinib [53]. We decided to leave resistant patients out of the analysis for two distinct technical reasons. First, this scenario would require a more complex population dynamics model – i.e., with more subopulations –, characterized by many more parameters, often impossible to estimate. Second, in this case the identification of an optimized therapy should involve two distinct controls, and even if theoretically possible, this would require to obtain data concerning the effect of Dasatinib/Nilotinib/Ponatinib/Bosutinib on tumor burden, which are not present in the used dataset.

We eventually selected 22 (out of 29) patients, for which the therapy led to a successful major molecular response (MMR), i.e., the ratio of cells with BCR-ABL mutation is ≤ 0.1 on the International Scale [54].

### Patient-specific PK models

We first use patient-specific PK models by incorporating demographic factors - i.e., body weight, age and sex – in the clearance and in the volume of the distribution, as per Eqs. (3) and (4) [19](see Materials and Methods).

In Figure 2 the PK curves corresponding to the 22 patients (in blue) and the average PK model (in red) over a selected time window ([0, 48] hours) are displayed.

**Figure 2:**
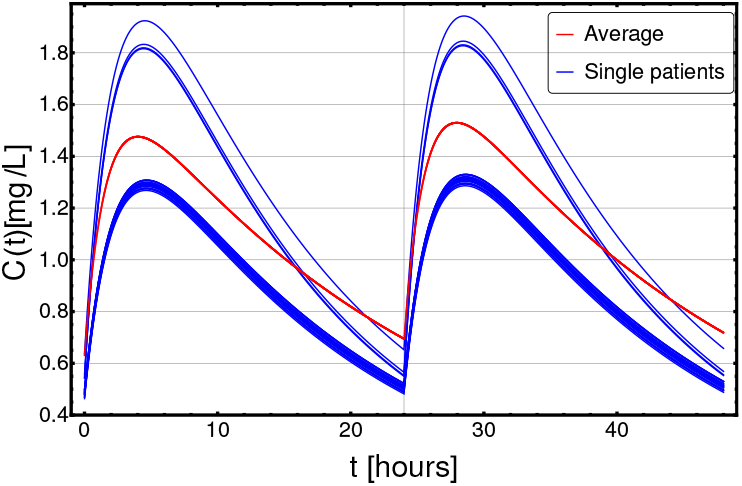
Patient-specific PK models. Personalized pharmacokinetic curves (blue solid lines), as estimated from demographic factors, such as age, sex and body weight, as per Eqs. (3) and (4), in the range *t* ∈ [0, 48] hours (x axis); y axis describes the drug concentration *C*(*t*) in [*mg/L*]. The blue curves correspond to the 22 distinct patients included in the dataset, whereas the dashed red curve represents the population-average pharmacokinetics.

### Defining personalized optimized administration at diagnosis time

CT4TD can be employed when CML is diagnosed, in order to identify optimized therapeutic strategies that lead to drug concentrations as close as possible to given targets, as. We here present the application of CT4TD to two distinct targets.

The first target concentration is *C*_*targ*_(*t*) = 0.57 [*mg/L*], which is currently the most widely employed in the clinic [17]. It is hypothesized that any effective therapy should ensure a drug concentration close to, but strictly larger than this value, in order to lead to a good performance, while minimizing the AEs [55, 56]. In this case, we consider this concentration as a *lower-bound* target, and the goal of the CT4TD framework will be to design an optimized therapy to be close to, but strictly larger than this concentration value, by employing an opportune distance notion (see Materials and Methods).

The second target concentration is *C*_*targ*_(*t*) = 1 [*mg/L*] and is supposed to provide a more effective therapy, but at the cost of an increased likelihood of AEs and toxicity [16, 57, 18]. From common practice, an effective therapy is that leading to values of drug concentrations *around* this target. For this reason, CT4TD will return an optimized therapy ensuring a drug concentration *as close as possible* to this *optimal* target (see Materials and Methods). Notice that one could select any arbitrary target concentration, or even combinations of targets, and this would not affect the validity of our approach.

For both targets we tested distinct settings, in which we considered, respectively, 1 and 3 doses per day at fixed times (i.e., 1 dose each 24 and 8 hours, respectively)^1^.

In Figure 3 we present the application of CT4TD to a selected patient (n. 0001 00004 AJR, male), with respect to the *lower-bound* target *C*_*targ*_(*t*) = 0.57 [*mg/L*] (left panels), and the *optimal* target *C*_*targ*_(*t*) = 1 [*mg/L*] (right panels). In particular, we compared the standard administration (red), the 1-dose optimized therapy (blue) and the 3-doses optimized therapy (green), on a temporal window of 14 days, with respect to drug dosage (panels A–E),drug concentration in blood (B–F), cumulative (Euclidean) distance with respect to the target concentration (C–E), and AUC (D–H).

**Figure 3:**
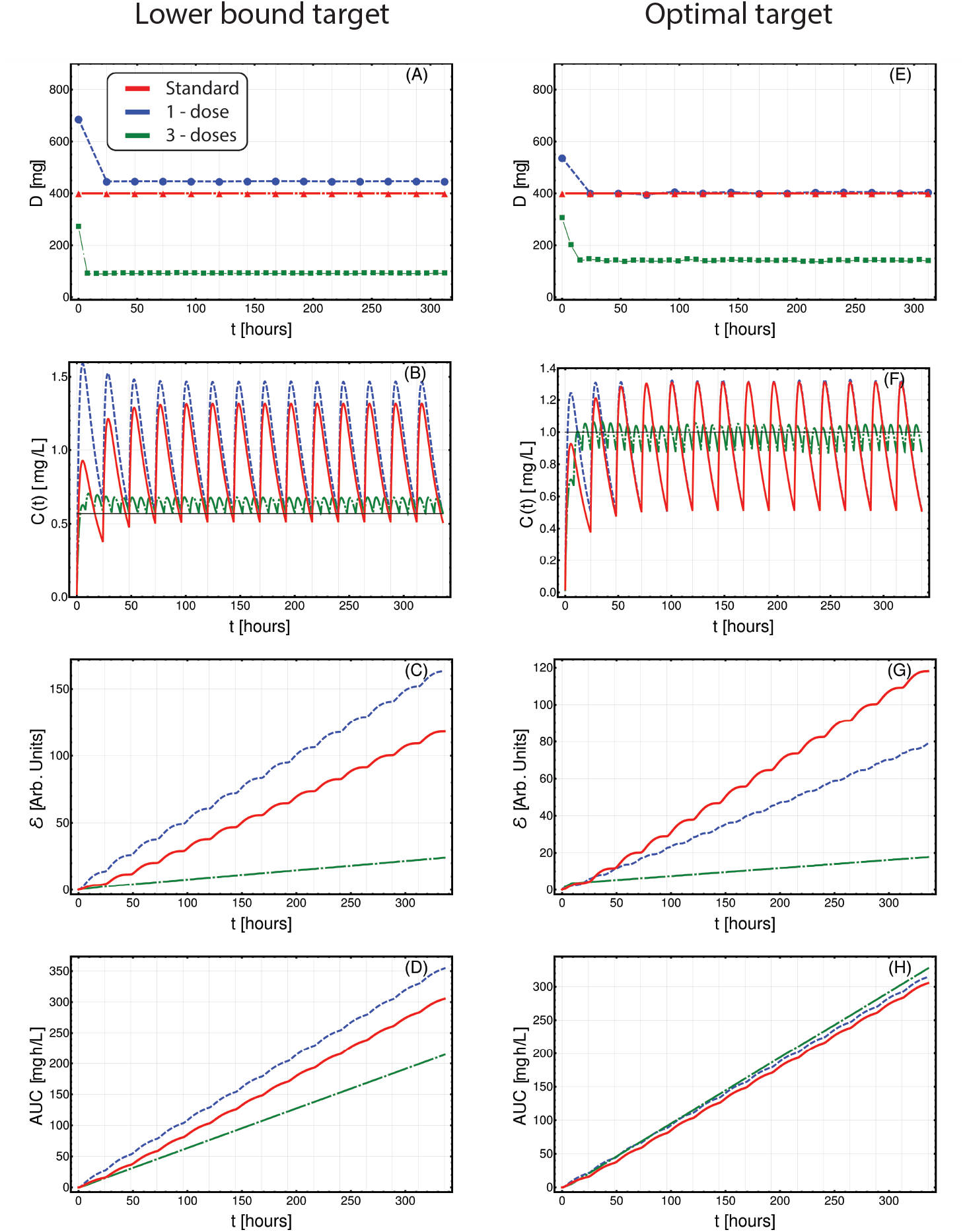
Patient-specific optimized therapy with fixed target drug concentrations. Optimized Imatinib administration returned by CT4TD for patient 0001 00004 AJR from [6], in the cases of: 1-dose/day (blue) and 3-doses/day (green), with respect to: *lower-bound* target concentration *C*_*targ*_ = 0.57 [*mg/L*] (left panels: A–D), and *optimal* target concentration *C*_*targ*_ = 1 [*mg/L*] (right panels: E–H). Standard administration – i.e., 400 mg Imatinib/day – is shown with a red dashed line. In this case, the optimization is obtained on patient-specific PK parameters, without considering the PD models. **(A,E)** Imatinib scheduled dosage in *mg* (y axis), displayed on 14 days (x axis). **(B-F)** Imatinib concentration in blood in [*mg/L*] (y axis). **(C-G)** Variation of the cumulative distances between the observed concentration and the selected target in time. **(D-H)** Temporal variation of the AUC in [*mg* · *h/L*].

When assessing the goodness of a therapy in the lower-bound scenario – i.e., *C*_*targ*_ = 0.57 [*mg/L*] – it is important to look at both the distance to the target *and* the overall time in which the drug concentration is above such target. In Figure 3A-D one can see that the optimized 1-dose strategy displays higher cumulative distance and area under the curve – AUC – with respect to the standard schedule, due to the fact that drug concentration is always strictly larger than the lower-bound target. This is proven by the proportion of time spent above the target (computed on the whole period), which is 100%, 100% and 88.6%, for the 1-dose, the 3-doses, and the standard administrations, respectively. Notably, the 3-doses optimized strategy displays a remarkable improvement also with respect to cumulative distance and AUC, proving to be an effective therapeutic choice for this specific patient.

With respect to the optimal target – i.e., *C*_*targ*_(*t*) = 1 [*mg/L*] –, an effective therapy should ensure a drug concentration as close as possible to the target, thus reducing the drug surpluses, while minimizing the cases of insufficient dosage. In this case, the 1-dose optimized scenario almost overlaps with the standard administration (yet, this is not always the case as one can see, for example, with respect to 0006 0007 AJR in Figure S20), whereas the 3-doses optimized strategy displays a significant improvement in terms of cumulative distance, as the drug concentration is constantly kept much closer to the desired target, thus importantly reducing under- and over-dosing (Figure 3E-H).

We also assessed the robustness of CT4TD via synthetic tests, (*i*) by introducing a stochastic and uniformly distributed noise over distinct PK parameters, and (*ii*) by generating an optimized schedule for a given set of PK parameters and then applying such schedule to a simulated patient with distinct parameters (i.e., inter-patient variability). The analyses can be found in the Supplementary Information. In both cases, the results produced by CT4TD proved to be robust with respect to noise and possible technical/measurement errors (see Supplementary Figures S8-S9).

### Adjusting treatment for tumor burden reduction

The CT4TD framework can be employed in order to identify optimized therapeutic strategies for patients that are currently treated with a standard regime, and for which longitudinal data on tumor burden variation are available. In this case, in order to estimate personalized PD models from experimental data – which describe the individual therapeutic response to identical drug concentrations –, CT4TD employs a module which fits longitudinal data on tumor burden with a hierarchical model of cancer population dynamics. In particular, we fitted each patient’s data with a biphasic exponential, which in log-scale describes the presence of two straight lines with distinct slopes, as proposed in [6, 7, 8] (see Materials and Methods for further details). With a few assumptions, the slope of such lines can be used to estimate the parameters of a 2-compartment population dynamics model of CML and, in particular, the (stem) cancer subpopulation death rate in presence of a standard Imatinib therapy – i.e., 400 *mg* per day – in each patient. This allows to estimate the patient-specific parameters of the PD model. The results of the data analysis on all patients are presented in Supplementary Table S3 and in Supplementary Figures S3-S6. In Figure 4-A one can see the personalized PD curves for the 22 patients, computed via (9), as compared to the average one.

**Figure 4:**
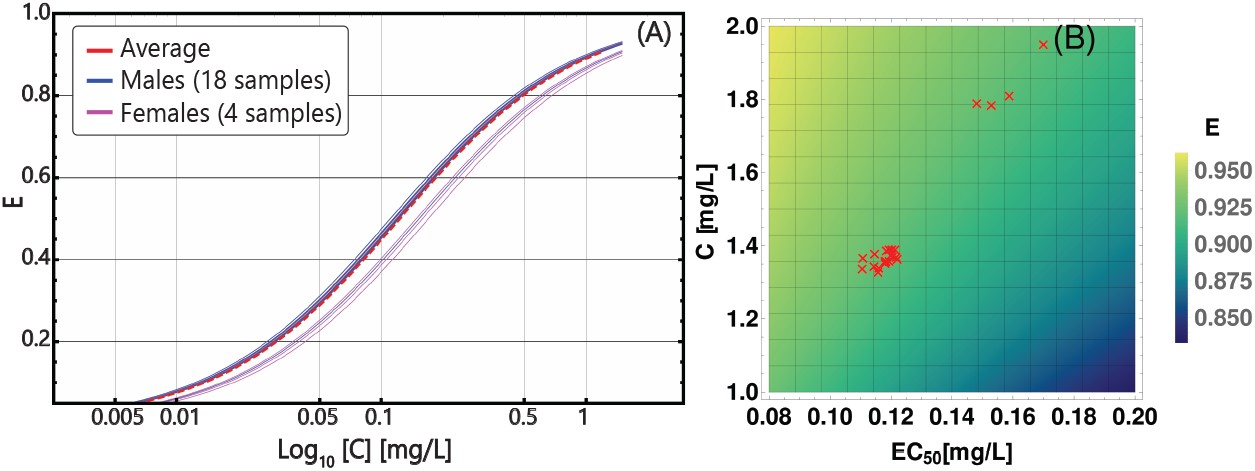
Patient-specific PD models. **(A)** Personalized PD curves obtained from (9) by using *E*_*max*_ = 1 and *n* = 1 for all patients, and distinct values of *EC*_50_, based on the death rate of cancer stem cells, as estimated from longitudinal data on tumor burden; x axis (*Log*_10_ scale) describes the concentration in the range *C* ∈ [0, 1.2] [*mg/L*], on y axis the efficiency *E* is displayed. The solid blue curves correspond to the 18 distinct male patients included in the dataset and the solid pink curves correspond to the 4 distinct female patients and the dashed red curve represents the population-average pharmacodynamics. **(B)** Heat-map returning the variation of efficiency *E*, computed via (9), with respect to distinct parameters of the PK model – i.e., patient-specific time-average concentration *C̄* (y axis) – and of the PD model – i.e., patient-specific *EC*_50_ (x axis). Red triangles represent the 22 patients in the dataset.

We also assessed the relative relevance of the personalized parameters of the PD and PK models with respect to the efficacy of the therapy. The heat-map in Figure 4-B returns the variation of the efficacy with respect to combination of time-average concentration *C̄* and *EC*_50_, highlighting the personalized parameters of the 22 patients. As a first result, one can see that much of the variance in our dataset is due to differences in PK, rather than to PD, which however is still relevant. Notice also that the two visible clusters basically overlap with the male and the female groups, providing a possible explanation of the distinct therapeutic response observed in clinical studies [48]. We remark that the estimation of personalized PD models from experimental data of patients under treatment is one of the major novelties of our approach, and, in combination with the demographics-based PK models, allow to identify patient-specific therapeutic regimes that are optimized to minimize the tumor burden.

In order to identify personalized optimized therapies, we finally defined a cost function with the goals of: (*i*) maximizing the reduction of the tumor burden, and (*ii*) minimizing the toxicity and possible AEs, in terms of AUC (see Materials and Methods for further details). Such cost function requires to set opportune weights *W*_1_ and *W*_2_ for the two terms, respectively. In particular, a parameter 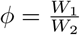 is defined, which can be opportunely tuned to favor either the first or the second term. However, the choice of a specific value for *ϕ* is arbitrary and depends on subjective research and clinical criteria.

To investigate the sensitivity of our framework to the variation of this parameter, we repeatedly applied the CT4TD framework to the CML dataset, by scanning various values of *ϕ*, and eventually assessed the differences in: (*i*) the time-average AUC as computed on a 14-days temporal window, and (*ii*) the efficiency computed on the time-average concentration in the same period, with respect to a 1-dose optimization scenario (the 3-doses scenario can be found in Supplementary Figure S51). In Figure 5 one can see the distribution of both quantities with respect to the 22 patients in the dataset, divided in males (blue) and females (pink), as compared to the average AUC and efficiency for the standard administration case (red).

**Figure 5:**
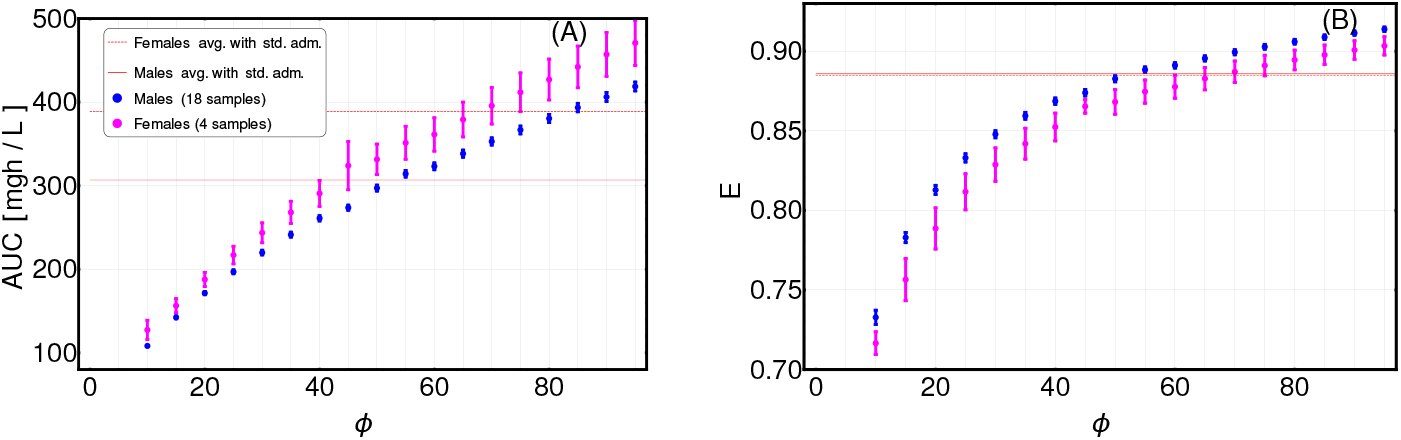
Assessment of term weights in cost function definition. The definition of the cost function for the adjusting treatment scenario requires to set the weights of the different terms. We here considered two terms, in order to: (*i*) minimize the tumor burden (weight *W*_1_), and (*ii*) minimize the AUC (weight *W*_2_) (see Materials and Methods for further details). We scanned the values of 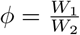 in the range [10, 100], by repeatedly applying the CT4TD framework to the 22-patients CML dataset from [6]. **(A)**. Distribution of the value of the AUC after 14-days of the optimized therapy retrieved by CT4TD (1-dose case), for distinct values of *ϕ*, with respect to the 22 samples in the datasets, divided in males (blue) and females (pink), and compared to the average AUC values returned by standard administration (400 mg Imatinib/day) in males (red solid line) and females (red dashed line). **(B)** Distribution of efficiency computed via (9) on the time-average concentration over 14 days of the optimized therapy retrieved by CT4TD (1-dose case), for distinct values of *ϕ*, and compared to the average efficiency in the standard administration scenario (solid and dashed red lines overlap).

It is first important to notice, that the results are highly sensitive with respect to the choice of *ϕ*, and can either display improvements (e.g., higher AUC and/or lower efficiency) or worsening with respect to the standard case in distinct cases. Moreover, male and female groups show significantly different distributions, thus pointing at physiological differences that should be considered in therapy design. As a rule-of-thumb, we suggest to select a value of *ϕ* for which a slightly larger value of efficiency is observed, while not inducing a too high increase in AUC. In our case, we selected a value of *ϕ* equal to 60 for men and of 75 for women.

In Figure 6A-C we show the comparison among the actual therapeutic regime administered to a selected patient (n.0001 00004 AJR, male – code ID in Supplementary Table S3) and the optimized therapies identified via CT4TD by setting *ϕ* = 60, in both 1-dose (blue) and 3-doses (green) scenarios, in terms of: (*i*) drug dosage, (*ii*) drug concentration, (*iii*) AUC and (*iv*) variation of the tumor burden in time. In particular, the temporal evolution of the tumor burden from diagnosis to the present is displayed by showing the experimental data points (purple) and the best fit (red), whereas the predicted future evolution is shown with respect to the 1-dose (blue) and the 3-doses (green) optimized strategies.

**Figure 6:**
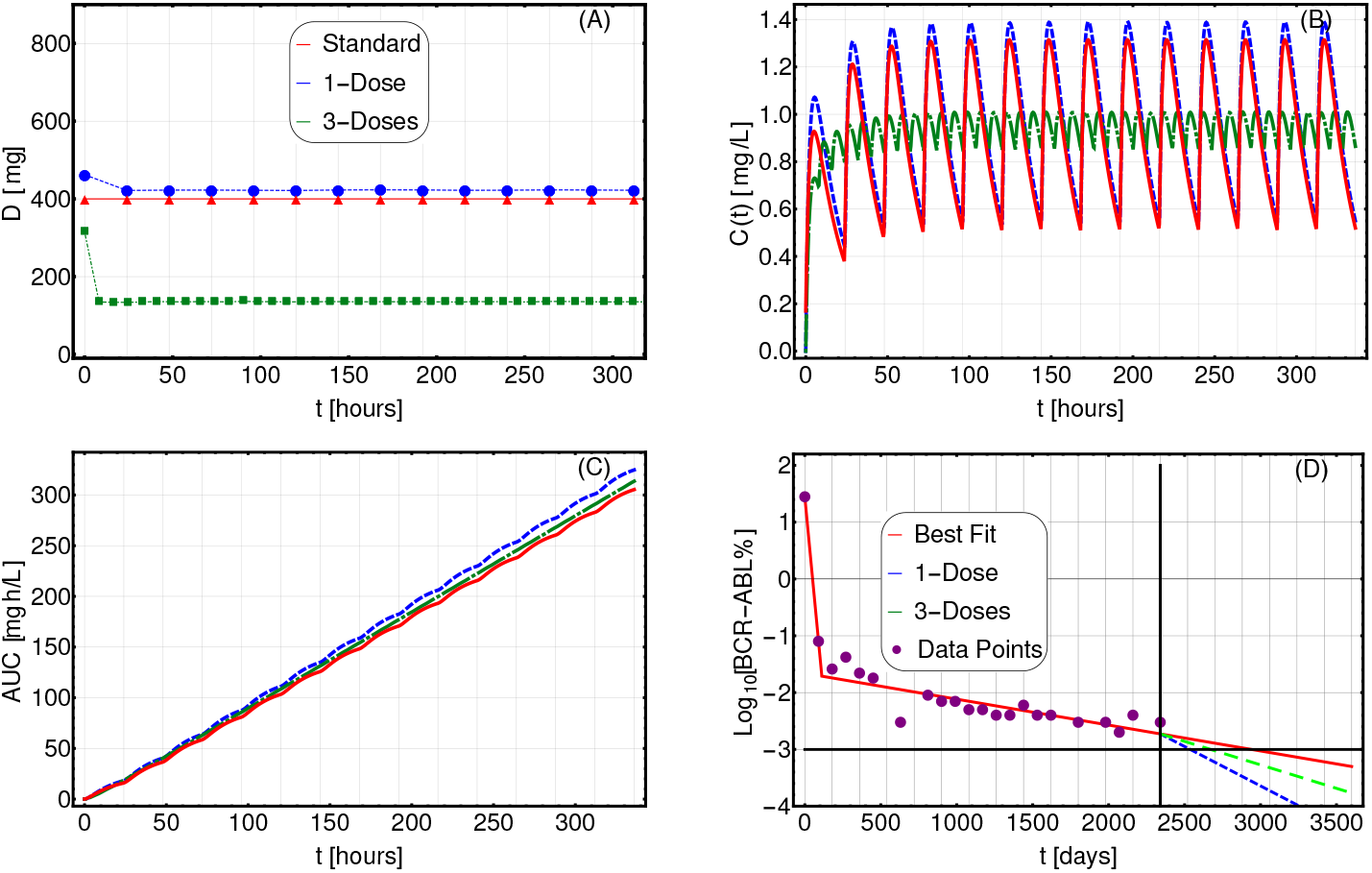
Adjusting therapy for tumor burden minimization. Imatinib administration optimized for tumor burden minimization in patient 0001 00004 AJR (male) from [6], in the cases of: 1-dose/day (purple) and 3-doses/day (green), with *ϕ* = 60, as compared to standard administration (red). In this case, the optimization is obtained on patient-specific PK and PD models. **(A)** Imatinib scheduled dosage in *mg* (y axis), displayed on 14 days (x axis). **(B)** Imatinib concentration in blood in [*mg/L*] (y axis). **(C)** Temporal variation of the AUC in [*mg · h/L*] (y axis). **(D)**Longitudinal data points on tumor burden recorded in the interval *t* ∈ [0, 2340] days (purple). The best fit is shown with red lines. The slope of the right-most line is used to determine the cancer stem cell death rate and, in turn, the patient-specific PD parameters. The blue (green) line represents the predicted cancer subpopulation decay in case the 1-dose (3-doses) optimized therapy was adopted from day 2340 to day 3500.

The most striking result is that, given similar AUC curves (i.e., similar toxicity and AEs), both the 1-dose and the 3-doses optimized strategies lead to a significantly faster predicted tumor burden decay. In particular, the tumor burden decay is respectively 3.07 and 1.82 times faster for the 1-dose and the 3-doses regimes, with respect to standard administration. This result paves the way for an automated strategy for therapy adjustment design, which might be further developed by employing closed-loop controllers.

In Supplementary Figures S31-S50 one can find the results of the analyses on all the 22 patients in the dataset.

## Discussion

The introduction of the CT4TD framework aims at filling the lack of automated and data-driven procedures for decision support in health care and personalized therapy design in cancer, especially by exploiting the increasing available computational power, which allows one to perform large-scale simulations and efficient search in the parameter space, and to deal with noisy and imperfect data.

In particular, CT4TD aims at overcoming the limitations of current control-based methods for therapeutic hypothesis generation. First, its completely general theoretical approach allows to consider: (*i*) any disease for which a PK/PD model can be derived and its parameters measured, (*ii*) any kind of administration, e.g., continuous drug infusion or discrete doses, (*iii*) any measurable term that is considered as relevant in the definition of a therapeutic *cost*. CT4TD eventually allows to evaluate in silico the outcome of the designed therapy.

Furthermore, CT4TD introduces the possibility of designing optimized therapeutic strategies based on experimental data concerning the disease progression.The identification of data-based patient-specific PK/PD models is one of the major novelties of CT4TD and has a profound impact on the characterization of tumour heterogeneity and, accordingly, on the customization of cancer therapies.

Several developments of CT4TD are underway. In particular, the possibility of tuning the PK/PD models to include information on the somatic evolutionary history of the tumors [58, 59, 60] will be essential in delivering more effective personalized therapeutic strategies. This is especially important for tumors displaying high levels of intra-tumor heterogeneity, which is known to be responsible for drug resistance, therapy failure and relapse [61].

As CT4TD relies on the RedCRAB optimization framework [10], the overall procedure could be implemented in remote, paving the way for a wireless decision support system for therapy design, to be used directly by clinicians [62]. In this respect, as a future development, an open-source computational tool will be made available to the scientific community, allowing to perform individual-specific analysis for a wide range of disease.

## Materials and Methods

### Patient-specific PK models of Imatinib in CML

We here employ the PK model of oral administration of Imatinib introduced in [19]: if *χ*_*g*_(*t*) is the amount of Imatinib in the gastrointestinal tract, *k*_*a*_ is the first order absorption rate, *f* is the bioavailability, i.e., the fraction of an administered dose of unchanged drug that reaches the circulatory system, and *D* the ingested dose, then:

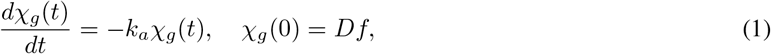

so if *v* is the volume of the distribution, i.e., the theoretical volume needed to account for the overall amount of drug in the body in case the drug was evenly distributed throughout the body, *CL* the clearance, i.e., the volume of plasma cleared of the drug per unit time, *C*(*t*) the concentration in the blood, *χ*_*b*_(*t*) the amount of Imatinib in the blood 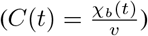, then:

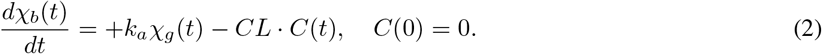

An example of the solution of (2) can be found in Supplementary Figure S2.

Both equations can be tuned to consider demographic factors as body weight, age and sex, thus providing patient-specific PK models. More in detail, such demographic factors can be incorporated in the clearance *CL* and in the volume of the distribution *v*, as initially proposed in [19]:

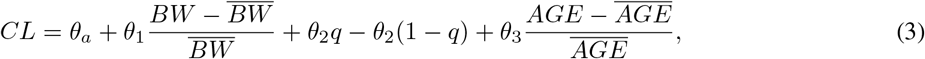

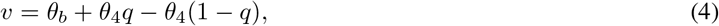

where *θ*_*i*_, for *i* = *a*, *b*, 1, 2, 3, 4 are constants, *BW* is the body weight of the patient and 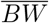 is its population-average, *AGE* is the age of the patient and 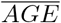 its population-average and *q* is a binary variable which takes value 1 for male and 0 for female. Estimation of such parameters is provided in [19] and shown in Supplementary Table S1 and S2. As in the dataset used in the case study, only the information about age and sex was available, we estimated the corresponding BW in each patient on the basis of average measures provided in [63]^2^.

### Population dynamics model of leukemia

CT4TD includes a data analysis module, aimed at identifying patient-specific parameters of the PD model. To this end, we define the simplest compartmental model of population dynamics for which it is possible to estimate the parameters from available experimental data.

In detail, the organization of leukemic systems is characterized by a hierarchy that is analogous to the healthy hematopoietic counterpart, and which can be modeled in the simplest case with two compartments: (*i*) cancer stem cells (CSC) and (*ii*) progressively differentiated cancer cells [6, 7, 12, 8, 13, 14]. Notice that the model could be generalized to account for *m* lineages, in order to represent the possible presence of subpopulations of tumor cells with distinct properties (e.g., therapy resistant phenotypes). However, more complex models appear not to improve the overall agreement with clinical data, due to the limited number of time points usually available.

In this case, first order differential equations are suitable to describe the population dynamics, because experimental evidences show that the proliferation of healthy cells display an exponential increase [45], whereas cancer cells under therapy display an exponential decay [6, 7, 8, 9]. Thus, we analyze the fluxes between compartments, by defining the following constants:

- *p*_*i*,*k*_ is the division rate of the cells in the *i* ensemble of the *k* lineage.
- *d*_*i*,*k*_ is the death rate of the cells in the *i* ensemble of the *k* lineage – this rate will be estimated from experimental data.
- *a*_*i*,*k*_ ∈ [0, 1] is the probability that, when a cell undergoes mitosis, both of its daughters belong to the *i* ensemble in the *k* lineage; therefore, 1 − *a*_*i*,*k*_ is the probability of belonging to the *i* + 1 ensemble. With respect to CSCs (or SC) this quantifies the self-renewal process.

Note that, we consider a symmetric differentiation scheme, according to which after mitosis both cells are of the same type, either stem or differentiated.

In a single individual, the dynamics of the healthy system – which includes stem cells, progenitors and differentiated cells – and that of the leukemic system coexist. Yet, we here assume that CML cells are cytokine-independent [13, 14], so the equations describing the dynamics of leukemic subpopulation do not include terms related to the healthy counterpart. As a consequence, the dynamics of the leukemic system can be defined as follows:

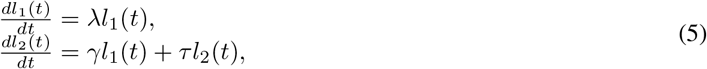

where:

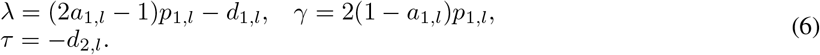

This is a typical example of a linear autonomous system, and the solution could be obtained analytically in a recursive way:

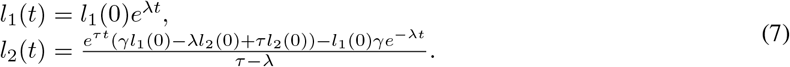

### Experimental data fitting

Once the leukemia 2-compartments model has been defined, it is possible to estimate its parameters from experimental data. In particular, we here focus on the specific case of CML. As every cancer cell in CML is characterized by the BCR-ABL mutation, it is possible to distinguish healthy from cancer cells with Q-PCR measurement, and this allows to have longitudinal experimental data returning the fraction of cancer cells over the total, i.e., the tumor burden [6, 7, 8, 9].

The CT4TD fits the longitudinal data on tumor burden in each patient with a biphasic exponential, which in log-scale this describes two distinct and intersecting lines, as proposed in [6, 7, 8]. In particular, we selected the combination of straight lines minimizing the value of *R*^2^, by scanning all the points of intersection (with a step of 1 day) and fitting data with two lines with distinct slopes, via a standard non-linear fit (see the Supplementary Table S3 for all parameter estimation). We also tried to fit data with either one or three distinct lines, yet in our case study the best fit was obtained in the two-lines case.

Once the two best fitting curves have been obtained for each patient, we adopt a simplifying assumption that allows us to estimate the parameters of the compartmental model from data. In [64] it is shown that the leftmost curve (with higher slope) is likely to represent the overall population dynamics involving cancer stem cells, cancer progenitors and cancer differentiated cells (decreasing in population size), together with that of healthy blood cells (increasing in population size). Considering that the Q-PCR measurements of the BCR-ABL fusion gene return the ratio between cancer cells and the total number of cells in the system, it would be impossible to disentangle the contribution of each subpopulation to the overall dynamics, and to reliably estimate the values of of *γ* and *τ* in (6), without ad-hoc assumptions and/or further opportune experiments.

Instead, it is possible to hypothesize that the rightmost curve (with lower slope) accounts for the dynamics involving a completely recovered healthy cell subpopulation – thus, healthy cells can be considered as constant in number – and a decaying cancer stem cell subpopulation, with no progenitors and differentiated cancer cells left in the system, as a consequence of the therapy [6, 7, 8, 9]. With this assumption, it is possible to estimate the parameters of the first compartment, and in particular, the cancer stem cell death rate *d*_1,*l*_ in (8), from experimental data. This also allows us not to explicit consider a model for the healthy hematopoietic system.

In detail, we assume that the exponential decay of the rightmost curve – given by the first equation in (7) – accounts for the dynamics of the CSC subpopulation only. In this way, it is possible to evaluate the effect of a standard Imatinib therapy – 400 mg per day – directly on the CSC decay, as estimated from any patient’s data. In fact, *β*_*j*_, i.e., the measured slope accounting for the decay of CSCs, will be given by:

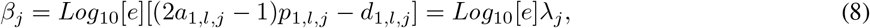

where *j* is the patient’s index.

### Estimation of patient-specific PD models from experimental data

Various PD models can be employed to estimate the efficacy of Imatinib in CML. In our case, we use a PD model based on the maximum-inhibition effect (*E*_*max*_) [17]:

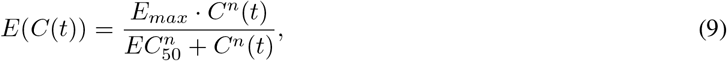

where *E*(*t*) is the effect, *E*_*max*_ is the maximum effect, *C*(*t*) is the concentration of the drug, *EC*_50_ the concentration of the drug that produces half of maximal effect, and *n* is a shape factor.

In order to identify patient-specific PD models from the parameters of the leukemia model estimated from data, we can safely suppose a linear relation between the population-average of the efficacy 〈*E*〉 and the population-average of 〈*d*_1,*l*_〉 [16]. Hence, the relation is the following:

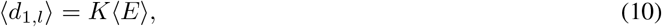

where *K* is a conversion constant.

It is possible to estimate *K* from the available dataset, by employing patient-specific PK models and an average benchmark PD model (*n* = 1, *EC*_50_ = 0.123 [*mg/L*] and *E*_*max*_ = 1). To do this, we first compute the time-average of the concentration *C̄*_*j*_ (*t*) for each patient, with respect to a 40-days standard therapy – 400 mg Imatib per day –, which we then use to compute the time-average of the efficacy *Ē*_*j*_ as per (9), by considering unique average PD parameters for all patients. Finally, we consider the population-average the efficacy (*E*), as computed on all patients.

〈*d*_1,*l*,*j*_〉 is then obtained by using formula (8), setting *a*_1,*l*_ = 0.87 and *p*_1,*l*_ = 0.45 [*days*^−1^] as proposed [14], and by taking the mean over all patients. As a result, the conversion factor for this dataset is *K* ≈ 0.377 ± 0.0007 [*days*^−1^]. At this point, we are able to estimate the personalized parameters of the PD model, and in particular of *EC*_50_, by supposing that the maximum efficacy is *E*_*max*_ = 1 and the shape factor is *n* = 1, for all patients [17, 65]. Therefore, the relation is the following:

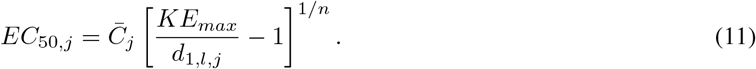

With this procedure, we can estimate the value of *EC*_50,*j*_ for each patient from individual longitudinal data on tumor burden. The results for all patients are presented in Supplementary Table S4 and in Figure 4.

Notice that, if longitudinal data on single patients are not available, the CT4TD allows to employ a unique (average) PD model for all patients, as estimated from experimental studies (see, e.g., [17, 16, 57, 18]).

### The PK/PD control problem

We formally define the *PK/PD control problem* for the administration of discrete doses as follows.

Let be *t*_*in*_ and *t*_*fin*_ the initial and the final time of the therapy, we aim at finding: (*i*) the optimal doses 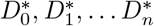, and (*ii*) the optimal schedule of administration 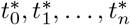, such that a functional that represents the “cost” 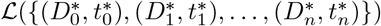 is minimized.

The definition of the *cost* functional 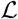 is the “core” of the PK/PD control problem and can include various weighed terms, which one should wisely select with respect the specific problem and goals. In particular, 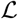 may (or may not) include distinct terms accounting for: (*i*) the efficacy *E* of the therapy, as derived via PD models, such as the Hill equation [21] or the *E*_*max*_ model [17] (if average or patient-specific parameters can be estimated); (*ii*) the toxicity of the therapy and/or the possible AEs as measured, e.g., via the Area Under the Curve (AUC); (*iii*) in case the PD model is unknown or indefinable – a distance between the optimized concentration *C*\*(*t*) and a target concentration *C*_*tg*_ as estimated, for instance, from clinical trials [18]; (*iv*) the economic cost of the therapy; (*v*) the properties and the temporal evolution of the disease, as in the case of the tumor burden estimation from longitudinal experimental data [13, 6, 46, 12, 14] (in this case the goal will be the optimization of the performance of the therapy with respect to the minimization of cancer subpopulations; (*vi*) the probability of developing resistance to the therapy [6, 7]; etc.

Obviously, some of this terms are highly correlated, as for example the AUC and the economic cost. Notice also that the choice of opportune weights is crucial in defining an effective control, if more than one term is used, and that it is necessary to fix *n a priori* when minimizing 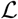. This could be useful if one wants to set the maximum amount of doses in a given temporal window. In detail, we here define cost functions 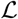 with respect to two distinct scenarios: (*i*) optimization of therapy for fixed target concentrations, (*ii*) optimization of therapy for tumor burden reduction.

### Therapy simulation

In order to simulate a therapy, we need to model a multi-dose oral administration. Let *t*_*in*_ and *t*_*fin*_ be the initial and the final time of the therapy, we suppose to give *n* + 1 doses at time *t*_0_, *t*_1_, *t*_2_, …, *t*_*n*_, with a dose amount *D*_*i*_ (*i* = 0, 1, 2, 3, … *n*), respectively. Thus, we have *n* first order differential equation like (1); by using the superposition principle the solution will be:

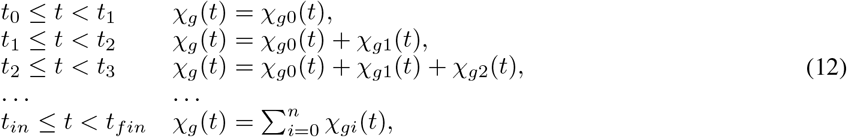

where 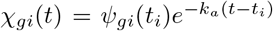 is a solution of (1), with *ψ*_*gi*_(*t*_*i*_) = *fD*_*i*_. Then, substituting *χ*_*g*_(*t*) into the (2) it is possible to study the dynamics of blood concentration of a certain drug for a multi-dose oral administration (see Supplementary Figure S2 for an example).

### Working scenario (*i*): optimal control with fixed target concentration at diagnosis time (patient-specific PK models – no PD models)

In many real-world scenarios it is not possible to retrieve or estimate the parameters of the PD models, for instance at the time of diagnosis. In this case, CT4TD can be used to find the best personalized therapeutic strategy to either: (*i*) be as close as possible to a given *optimal* target drug concentration, or (*ii*) be close to, but strictly larger than a given *lower-bound* target.

In the first case – i.e., *optimal* target concentration – CT4TD employs a simple Euclidean distance between two concentrations:

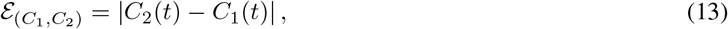

In this case, the *cost* is defined as follows:

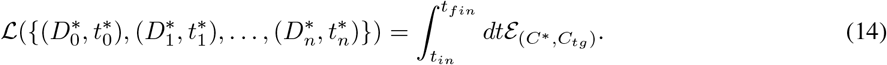

This *cost* favours the solutions which are close to the target, with no preference between above or under the target.

In the second case – i.e., *lower-bound* target concentration – CT4TD uses a “stepwise” distance in the space of concentrations. In this case the distance between two concentrations becomes:

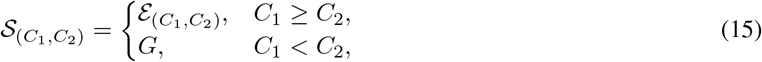

where *G* is a costant. It is then possible to define the *cost* as follows:

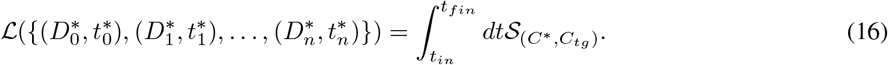

In this case, CT4TD will give solutions that display concentrations above the lower-bound target.

### Working scenario (*ii*): optimal control for tumor burden reduction (patient-specific PK models – patient-specific PD models)

When it is possible to estimate patient-specific PD parameters from longitudinal data on tumor burden variation under standard treatment, CT4TD can be used to identified an adjusted optimized therapy to reduce such burden in each patient.

In particular, our approach can use a *cost* function with the aim of: (*i*) minimizing the number of cancer stem cells (e.g., at the end of the treatment), by extremizing the exponent of (7) – this choice also allows us not to know or estimate the initial number of CSCs, *l*_1_(0); (*ii*) minimizing the AUC of the therapy, which accounts for toxicity and possible AEs.

To do this, we need to introduce two arbitrary weights *W*_1_ and *W*_2_, which account for the relative relevance of the two distinct components. Notice that we cannot consider the exponent of (7) only, as the dynamics of *l*_1_(*t*) is a monotone function and the minimization of cancer stem cells is reached for *D*_*i*_ → ∞, which implicates that if the maximum amount of drug is bounded (i.e., *D_max_*) the optimal solution is trivially reached for *D*_*i*_ = *D*_*max*_, ∀*i*.

Therefore, we have:

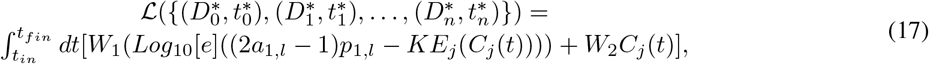

The ratio 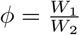 determines the overall performance of the optimal solution and should be wisely chosen. In order to provide some indications on this modeling choice, we performed an extensive scan of *ϕ* (in the range *ϕ* ∈ [10, 100]), with respect to all patients and we analyzed the variation of the time-average concentration *C̄*. The results are shown in Figure 5. From this analysis, one can see that a sound choice for *ϕ* might be in the range [60 - 65] for males and in [70 - 75] for females (note this choice depends on the units of measurement), meaning that the weight corresponding to the time evolution of CSCs is relatively more relevant than that corresponding to the toxic effects.

We finally remark that, in case longitudinal data are not available CT4TD can employ average PD models in the definition of the *cost* in (17), instead of the personalized ones.

### Optimal dosage

To solve the optimization problem with respect to dosages, we optimized a control field *D*(*t*) defined between *t*_0_ ≤ *t* ≤ *t*_*f*_ by using RedCRAB optimal control suite (See Supplemetary Information methods)

Then, we proceed by mapping the *D*(*t*) doses function into (*n* + 1)-integer values which correspond to (*n* + 1)-doses 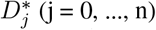, where *n* is the number of total doses given to the patient; *t*_*f*_ is the final time of the therapy and *t*_0_ = 0 the initial time. Accordingly, we can define the schedule of administration as:

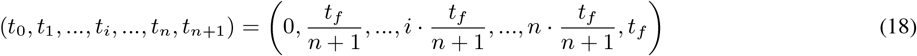

i.e. 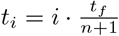 for *i* = 0, 1, …, *n* + 1.

Indeed, the n-doses 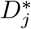 are obtained by integrating the doses function *D*(*t*) between adjacent times in the schedule administration as follows:

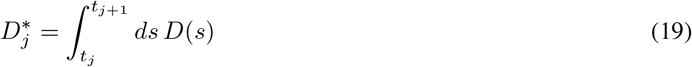

with *j* = 0, …, *n*. The optimal schedule case is described in the Supplementary Information.

## Acknowledgments

This work was partially supported by the Elixir Italian Chapter and the SysBioNet project, a Ministero dell’Istruzione, dell’Universitá e della Ricerca initiative for the Italian Roadmap of European Strategy Forum on Research Infrastructures. Furthermore we acknowledge financial support from IQST alliance. We thank Giulio Caravagna for helpful discussions.

## Footnotes

1 It would be possible to use our theoretical framework to define a free-time schedule optimization procedure. Yet, we believe that current practices in Imatinib oral administration would make a free-time schedule scarcely usable.

2 It is known that other factors can change the value of the clearance and the volume of the distribution. For instance, the concentration of the *α*_1_-acid glycoprotein affects the volume of the distribution, whereas the MDR1 genotype, the CYP3A4 activity and the creatinine clearance affect both *CL* and *v* [19]. However, we here limit to consider the aforementioned demographic measurements, because of their availability in our and in most datasets.

