## Supplementary Information for "Personalized Therapy Design for Liquid Tumors via Optimal Control Theory"

#### This PDF file includes:

Supplementary text
Figs. S1 to S51
Tables S1 to S6
References for SI reference citations

### Supporting Information Text

#### Methods.

**Optimization via RedCRAB .** In CT4TD the optimization is performed by using RedCRAB, the remote version of the dressed Chopped RAndom Basis (dCRAB) optimal control via a cloud server (1–4). Optimal control theory has been used for decades to optimize classical processes, and its quantum counterpart has been increasingly exploited in the last years (5–15). In its simplest version, optimal control drives the state of the system to a goal one, characterized by some desired properties, by using a set of time-dependent controls.

Here, the dynamics of the system is identified by the concentration of the drug in a certain compartment $C(t, D(t))$, which obeys the time evolution equation:

$$\frac{\partial C(t, D(t)))}{\partial t} = f(t, D(t), C(t, D(t))), \quad [1]$$

where $D(t)$ is the time-dependent control field, i.e. the doses function defined in section (see section **patient-specific PK models of Imatinib in CML**). The goal here is to optimize the drug administration schedule (see section **Optimal dosage**) while minimize the cost functional as defined in Sections **working scenario** *(i)* and **working scenario** *(ii)*.

Starting from the standard administration schedule $D_0(t)$, the optimization proceeds by looking for the optimal correction $g(t)$ such the the optimal administration schedule will be $D(t) = D_0(t) + g(t)$. Following (2), the correction $g(t)$ is expanded in a truncated function space, specifically in random Fourier components as:

$$g(t) = \Gamma(t) \sum_{k=1}^{n_c} [A_k \sin(\omega_k\, t) + B_k \cos(\omega_k\, t)] \quad [2]$$

where $\omega_k = 2\pi(k + r_k)/T$ and $r_k \in [-0.5, 0.5]$, $n_c$ is the total number of frequency used, $T$ is the final time, and $\Gamma(t)$ is a fixed scaling function to keep the values at initial and final times unchanged. In conclusion, the optimization problem is reformulated as the extremization of a multivariable function $\mathcal{L}(A_k, B_k)$ with fixed $\omega_k$, and can be efficiently solved numerically by searching the best combination of $\{A_k, B_k\}$ with the preferred method of choice, here a direct-search method (16). Notice that, each frequency $\omega_k$ is independently optimized: indeed, after a certain number of iterations, we move to the next $\omega_{k+1}$, by introducing an external loop on the frequencies, i.e. super-iterations. This allows the algorithm to include a high number of Fourier components and efficiently find the optimal solution (2).

In the RedCRAB optimization, the server generates and transmits a set of controls to the CT4TD, which evaluates the cost function, by interfacing with MATLAB and communicates it to the server completing one iteration. The optimization continues iteratively by providing the optimal set of controls as well as giving back the figure of merit, until the convergence is reached.

***Robustness analysis.*** In order to assess the reliability of the results produced by CT4TD, we tested its robustness with respect to intra-patient and inter-patient variability. To account for intra-patient variability, we introduced a stochastic and uniformly distributed noise, i.e., $\chi_r = [-\sigma_r, \sigma_r]$ with $r = k_a, CL, v$, to the following parameters of the PK model: $k_a$, $CL$ and $v$, for every time point in the analysis (see Materials and Methods). We performed 700 distinct PK simulations, on the average patient, in the specific scenario of a target concentration $C_{targ}(t) = 0.57 \quad [mg/L]$ and 1 dose per day. We then analyze the relative variation of the average *cost* $\Delta\mathcal{L}$, as compared to the noise-free case, with respect to the width of the distribution of noise $\sigma_r$.

In Figure S9-A, one can notice that $\Delta\mathcal{L}$ variation with respect to the noise level follows an approximately quadratic trend, which is proven by fitting the data points with a curve with equation $b + a\sigma_r^2$ (the complete results of the fit are provided in Supplementary Table S5). Notice that such results are in agreement with other works that use quantum optimal control (17–19). A robustness analysis on inter-patient variability was also performed, to assess the impact of possible mistakes in the application of the protocol, for instance due to: $(i)$ possible errors in the estimation of the patient's PK parameters , $(ii)$ scarce accuracy of the demographic study used to estimate the PK parameters, e.g., due to small sample size or sample imbalance. To this end, we generated an optimized schedule for a set of PK parameters $\{k_a, CL, v\}$ and then we applied such schedule to a simulated patient where a parameter at time is different, e.g., $\{k_a', CL, v\}$ with $k_a' = k_a + \delta_{k_a}$. We finally measured the difference of $\Delta\mathcal{L}$, as a function of $\delta_{k_a}$.

Also in this case, we show in Figure S9-B that the results produced by CT4TD are robust with respect to possible technical or measurement errors. In fact, with respect to an error of $\approx \pm 30\%$, we have a maximum difference $\approx 10\%$ in performance, as compared to the noise-free case.

***Optimal schedule.*** Analogously, we can optimize also the schedule of administration $(t_0^*, t_1^*, ..., t_n^*)$ with a new pulse $u_2(t)$. So we can introduce a new pulse $T(t) = u_2(t)$ and using a similar integral structure as before:

$$t_f^* = t_n^* + \int_{t_n}^{t_f} dt' \tilde{T}(t') = t_f \quad [3]$$

$\tilde{T}(t)$ is the normalized function with respect to the difference between the initial time $t_0$ and $t_f$, and is defined as follows:

$$\tilde{T}(t) = \frac{T(t)}{\int_{t_0}^{t_f} dt' T(t')} (t_0 - t_f) \quad [4]$$

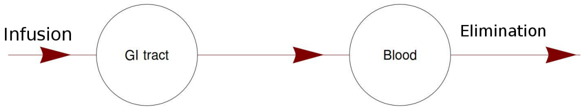


**Fig. S1.** Diagram of compartmental PK for oral administration of Imatinib.

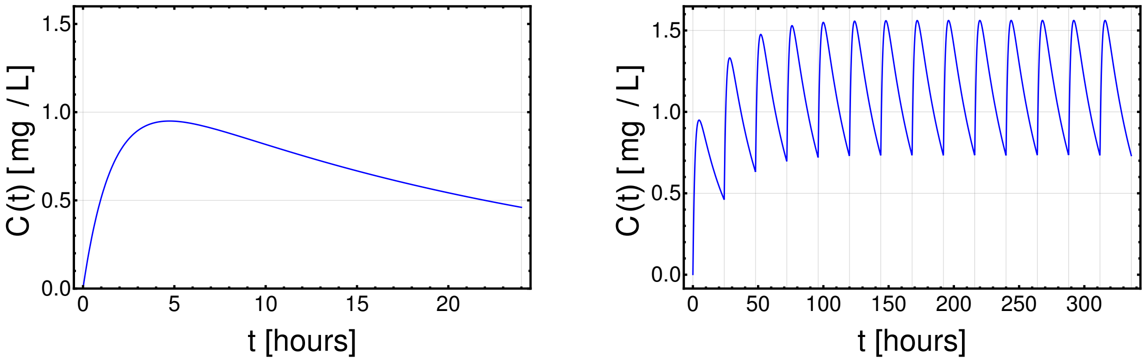


**Fig. S2.** On the left is the simulation of the concentration in the blood $C(t)$ of a single oral administration of $400mg$ of Imanitib. On the right there is a simulation of a multi-dose administration of $400mg$ of Imanitib, the drug is taken every 24 hours. Parameters of these simulation are presented in table S1

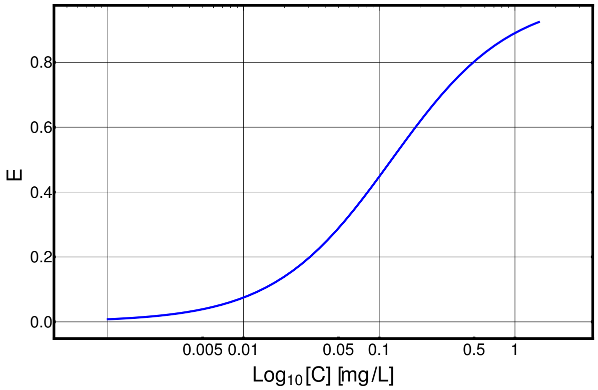


**Fig. S3.** Curve of efficacy given by Eq. (9), we impose $n = 1$, $EC_{50} = 0.123 \quad [mg/L]$ and $E_{max} = 1$.

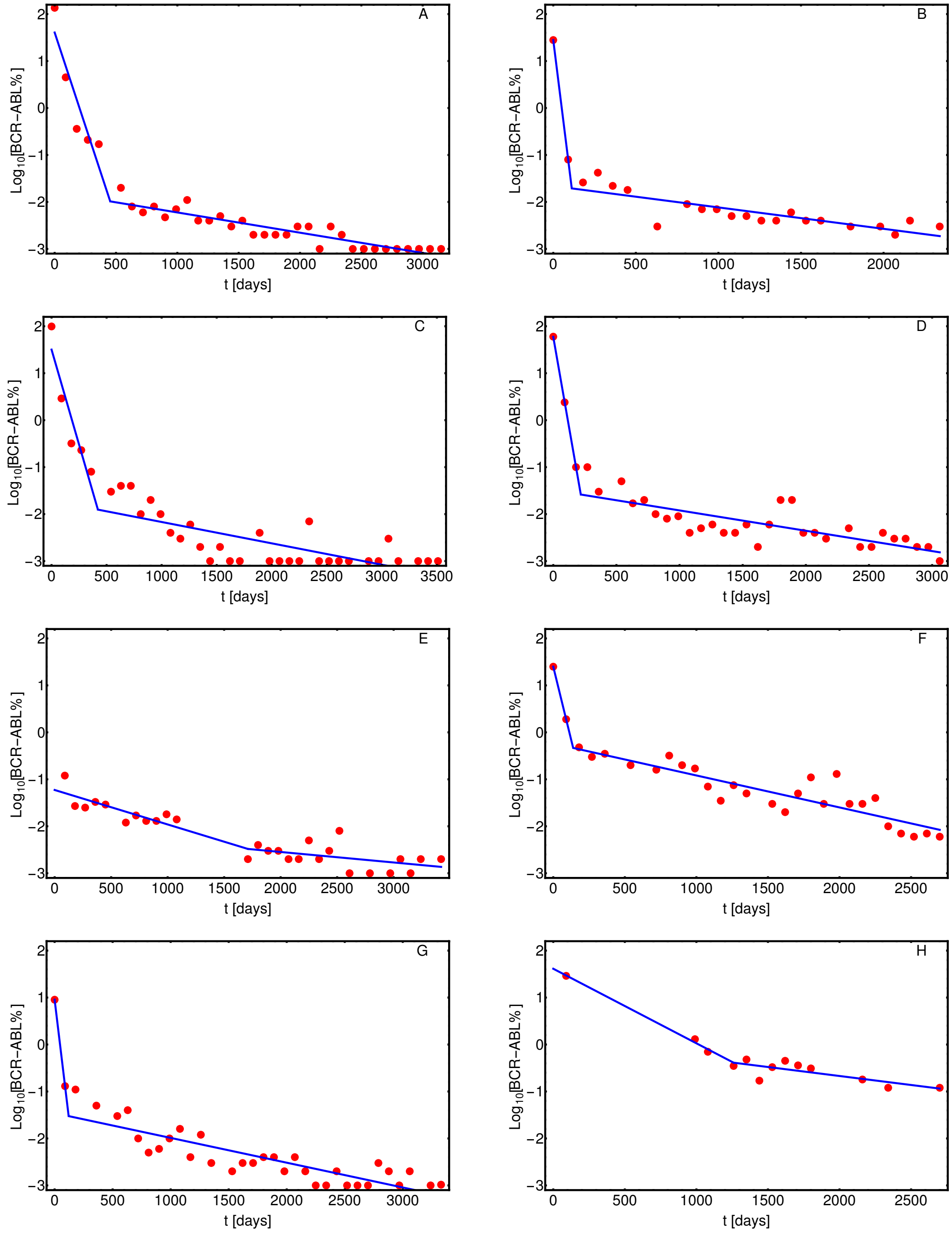


**Fig. S4.** Data analysis. The red dots are the experimental points and they represents the expression of BCR-ABL. Blue lines are the best fits. Results of the data analysis are summarized in table S3.

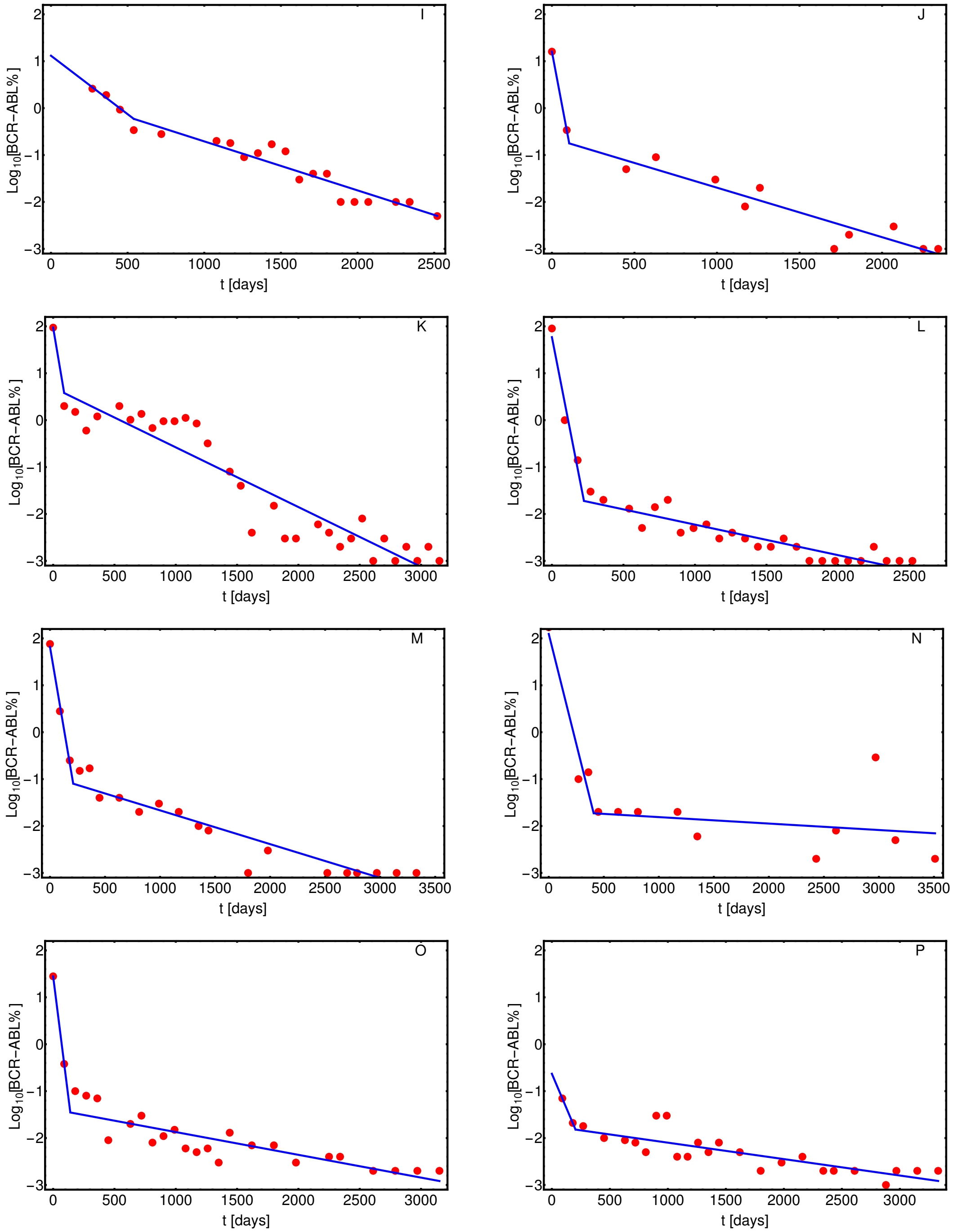


**Fig. S5.** Data analysis. The red dots are the experimental points and they represents the expression of BCR-ABL. Blue lines are the best fits. Results of the data analysis are summarized in table S3

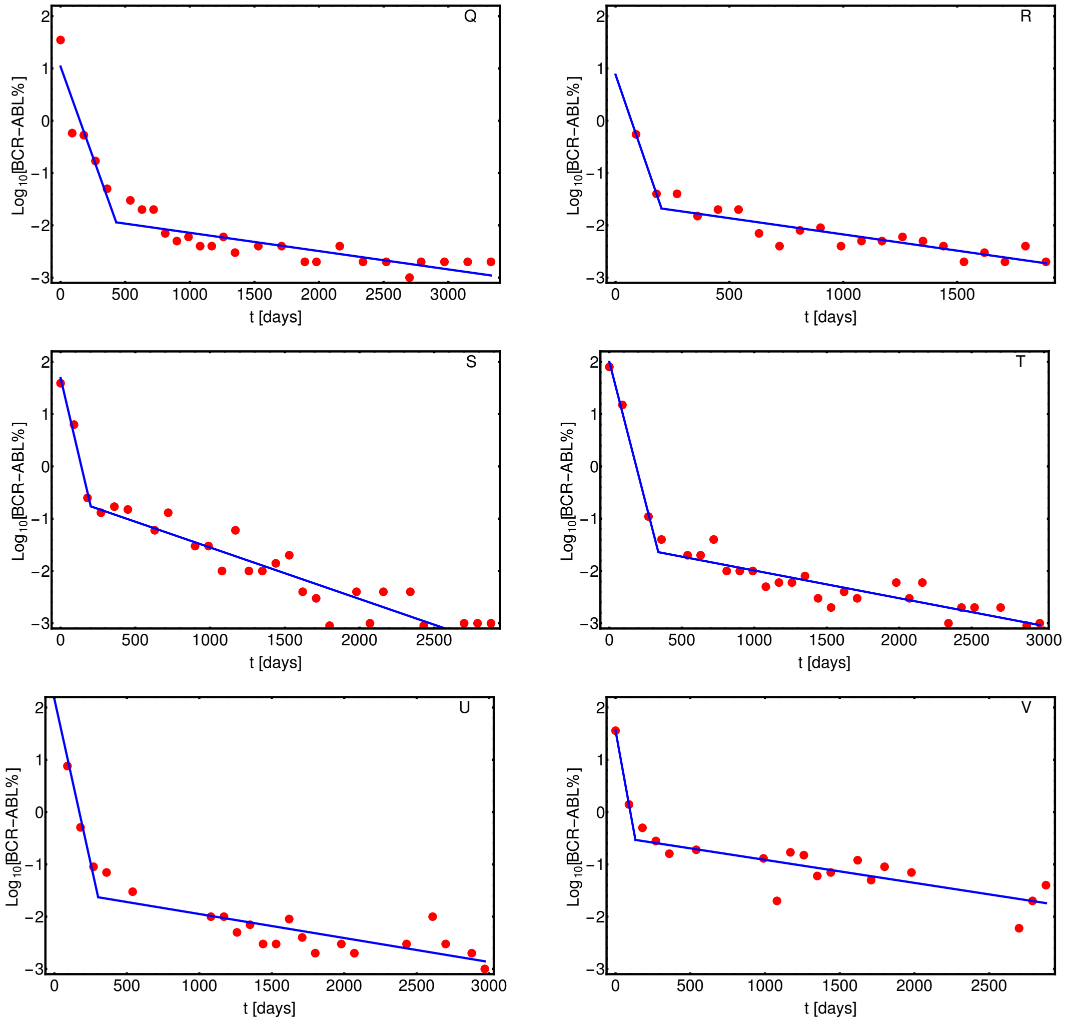


**Fig. S6.** Data analysis. The red dots are the experimental points and they represents the expression of BCR-ABL. Blue lines are the best fits. Results of the data analysis are summarized in table S3

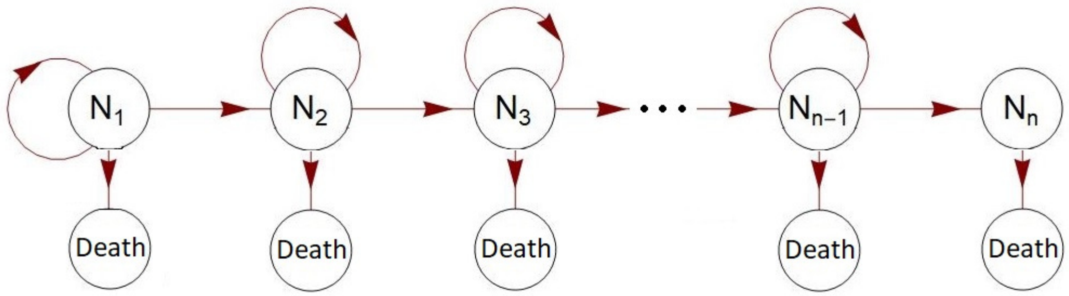


**Fig. S7.** Diagram of compartmental model for an unregulated leukemic cells lineage.

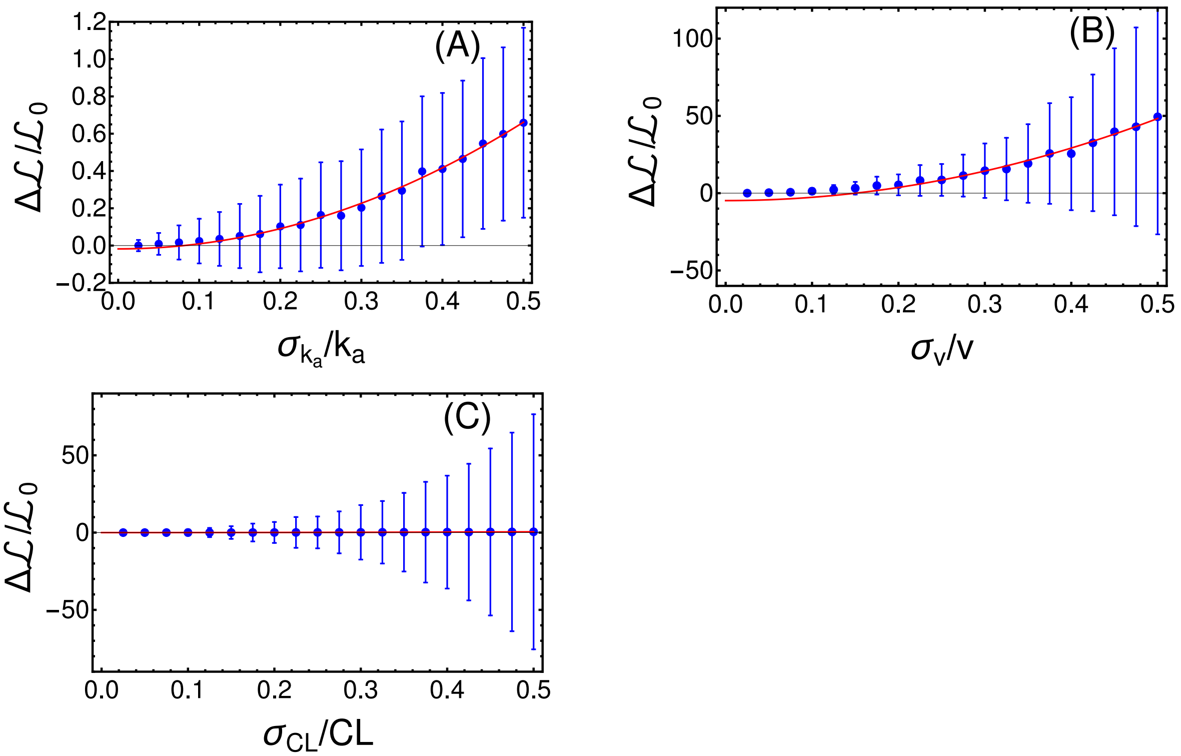


**Fig. S8.** Variation of the co respect the inter-patient noise, it is a stochastic noise picked random in the distribution $\chi_r = [-\sigma_r, \sigma_r]$. In figure $(\mathbf{A})$ we have $r = k_a$, in $(\mathbf{B})$ $r = v$ and in $(\mathbf{C})$ $r = CL$. Values of the fit are given in Table. S5

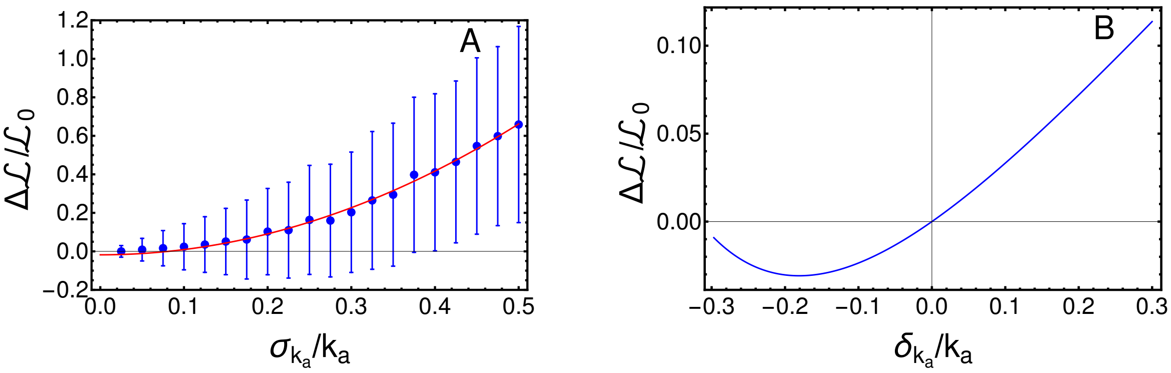


**Fig. S9. Robustness analysis with respect to intra- and inter-patient variability. (A)** Relative variation of the average *cost* with respect to the width of the distribution of intra-patient noise $\sigma_r$. $\mathcal{L}_0$ is the cost measured without noise and $\Delta\mathcal{L}$ is the difference between the cost in noise-free case and the average cost measured with distinct values of $\sigma_{k_a}$. Average values and standard deviation are displayed in blue, whereas the red solid line is the best fit $\Delta\mathcal{L}/\mathcal{L}_0 = 2.72 - 0.0178\sigma^2_{k_a}/k_a$. **(B)** Relative variation of the *cost* $\Delta\mathcal{L}/\mathcal{L}_0$ with respect to the relative value of the systematic error $\delta_{k_a}/k_a$

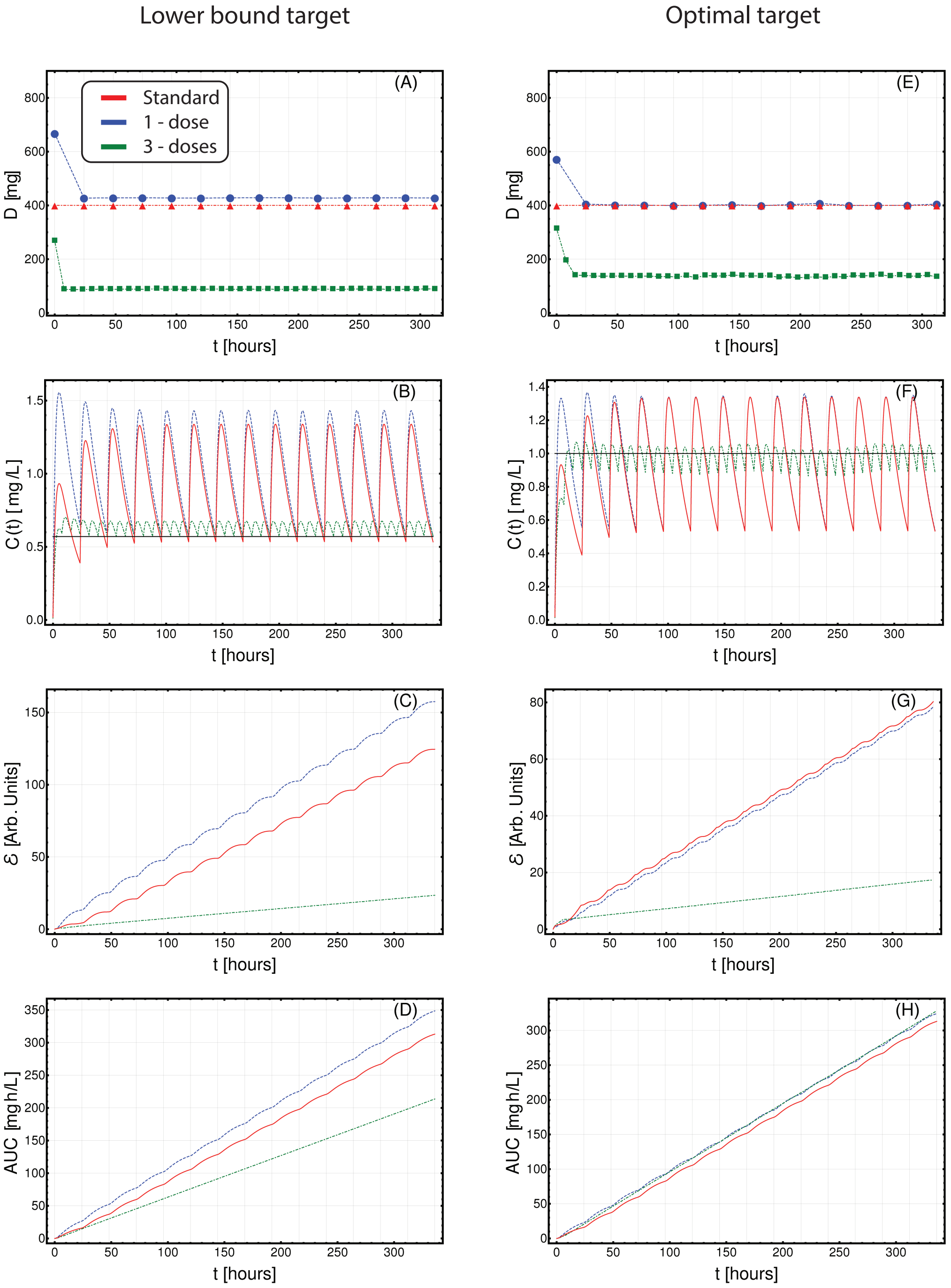


**Fig. S10.** Optimized Imatinib administration returned by CT4TD for patient 0001 00002 RH from (20), in the cases of: 1-dose/day (blue) and 3-doses/day (green), with respect to: $lower\text{-}bound$ target concentration $C_{targ} = 0.57\ [mg/L]$ (left panels: A–D) and $optimal$ target concentration $C_{targ} = 1\ [mg/L]$ (right panels: E–H). Standard administration – i.e., 400 mg Imatinib/day – is shown with a red dashed line. In this case, the optimization of PK/PD models is obtained on patient-specific PK parameters, without considering the PD models. **(A,D)** Imatinib scheduled dosage in $mg$ (y axis), displayed on $14$ days (x axis). **(B-E)** Imatinib concentration in blood in $[mg/L]$ (y axis). **(C-F)** Variation of the cumulative distance between the observed concentration and the target in time.**(D-H)** The AUC in $[mg \cdot h/L]$

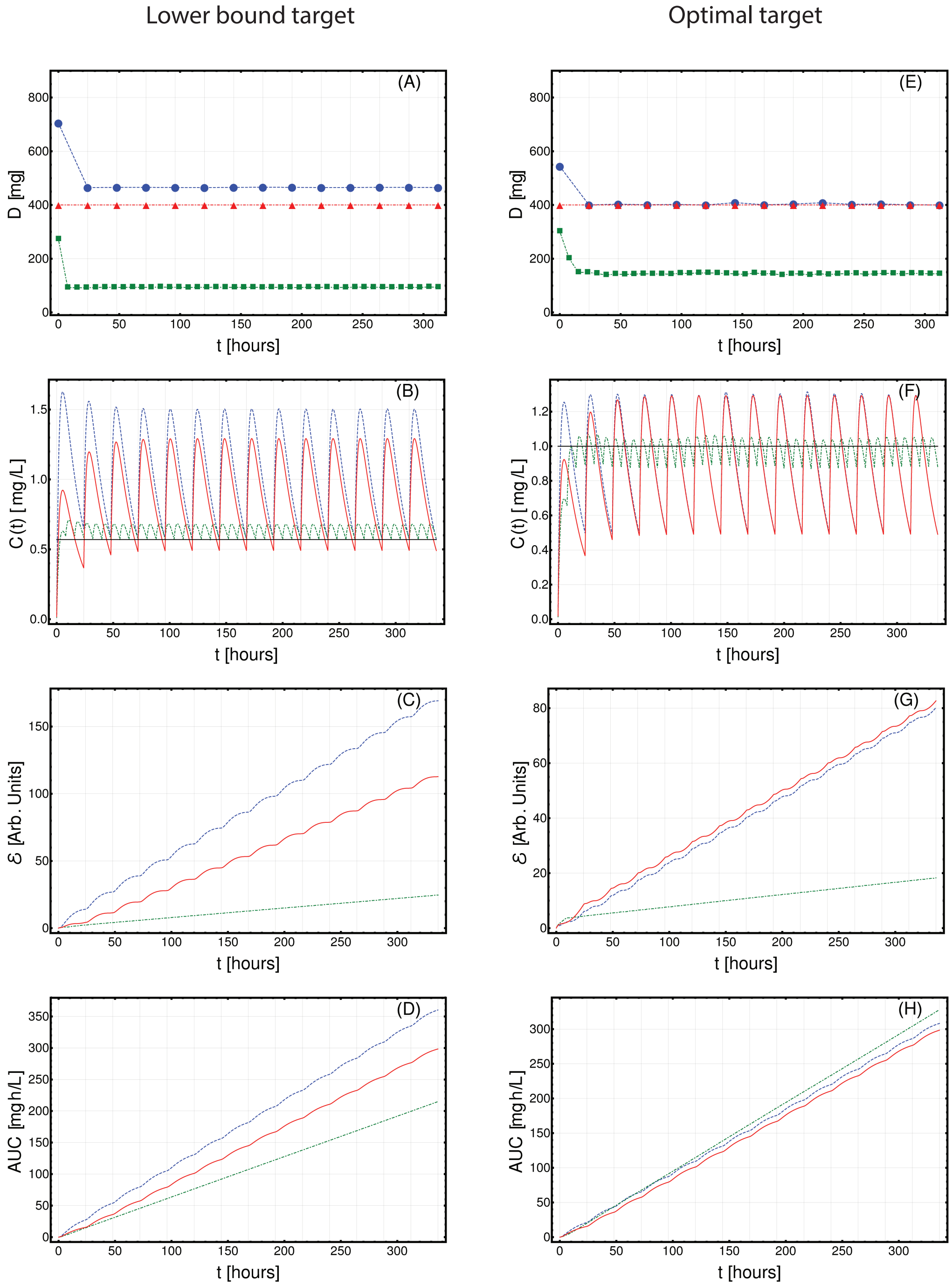


**Fig. S11.** Optimized Imatinib administration returned by CT4TD for patient 0001 00006 GMC from (20), in the cases of: 1-dose/day (blue) and 3-doses/day (green), with respect to: $lower\text{-}bound$ target concentration $C_{targ} = 0.57\ [mg/L]$ (left panels: A–D) and $optimal$ target concentration $C_{targ} = 1\ [mg/L]$ (right panels: E–H). Standard administration – i.e., 400 mg Imatinib/day – is shown with a red dashed line. In this case, the optimization of PK/PD models is obtained on patient-specific PK parameters, without considering the PD models. **(A,D)** Imatinib scheduled dosage in $mg$ (y axis), displayed on $14$ days (x axis). **(B-E)** Imatinib concentration in blood in $[mg/L]$ (y axis). **(C-F)** Variation of the cumulative distance between the observed concentration and the target in time.**(D-H)** The AUC in $[mg \cdot h/L]$

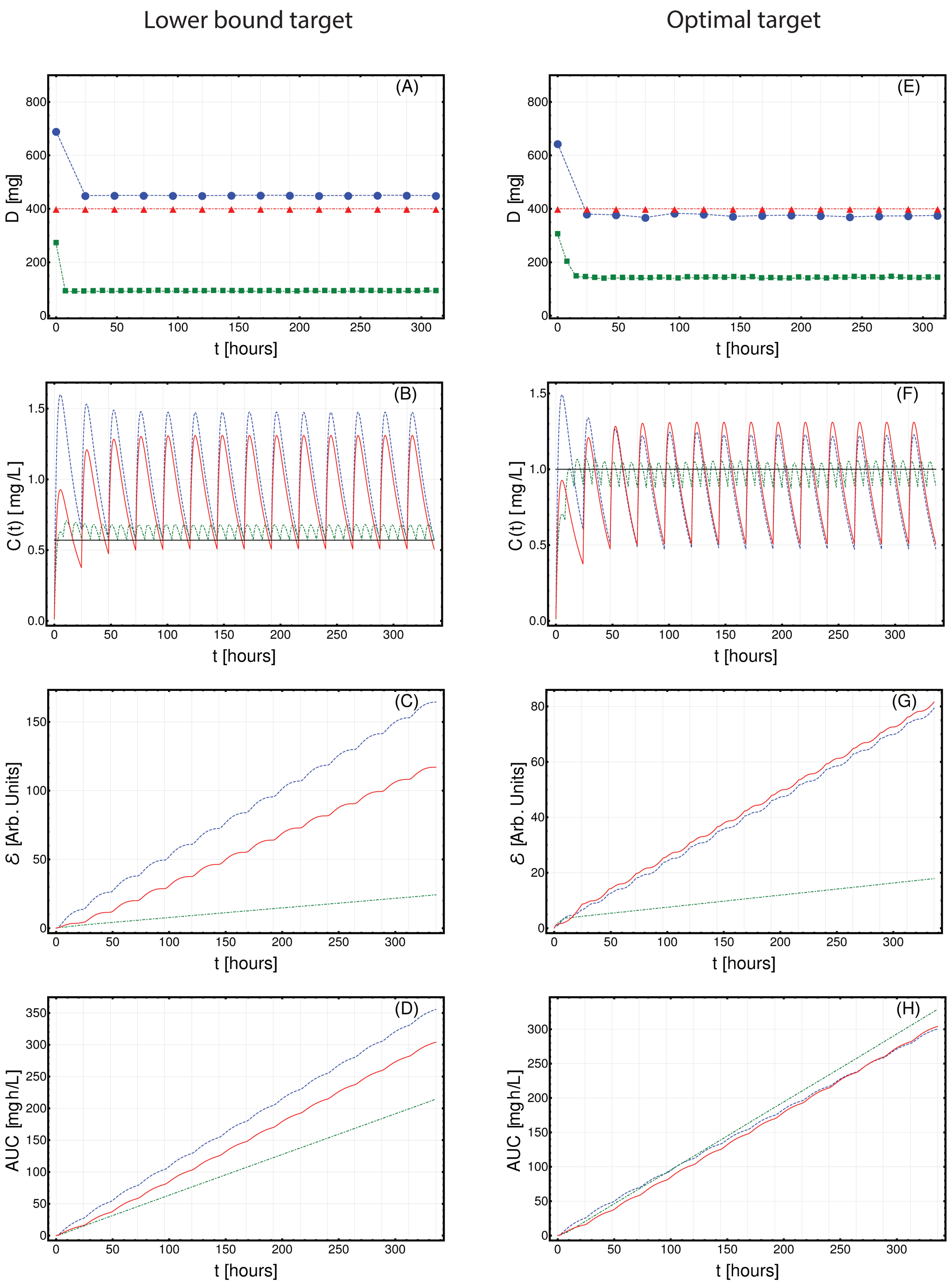


**Fig. S12.** Optimized Imatinib administration returned by CT4TD for patient 0001 00009 MJG from (20), in the cases of: 1-dose/day (blue) and 3-doses/day (green), with respect to: $lower\text{-}bound$ target concentration $C_{targ} = 0.57$ $[mg/L]$ (left panels: A–D) and $optimal$ target concentration $C_{targ} = 1$ $[mg/L]$ (right panels: E–H). Standard administration – i.e., 400 mg Imatinib/day – is shown with a red dashed line. In this case, the optimization of PK/PD models is obtained on patient-specific PK parameters, without considering the PD models. **(A,D)** Imatinib scheduled dosage in $mg$ (y axis), displayed on $14$ days (x axis). **(B-E)** Imatinib concentration in blood in $[mg/L]$ (y axis). **(C-F)** Variation of the cumulative distance between the observed concentration and the target in time.**(D-H)** The AUC in $[mg \cdot h/L]$

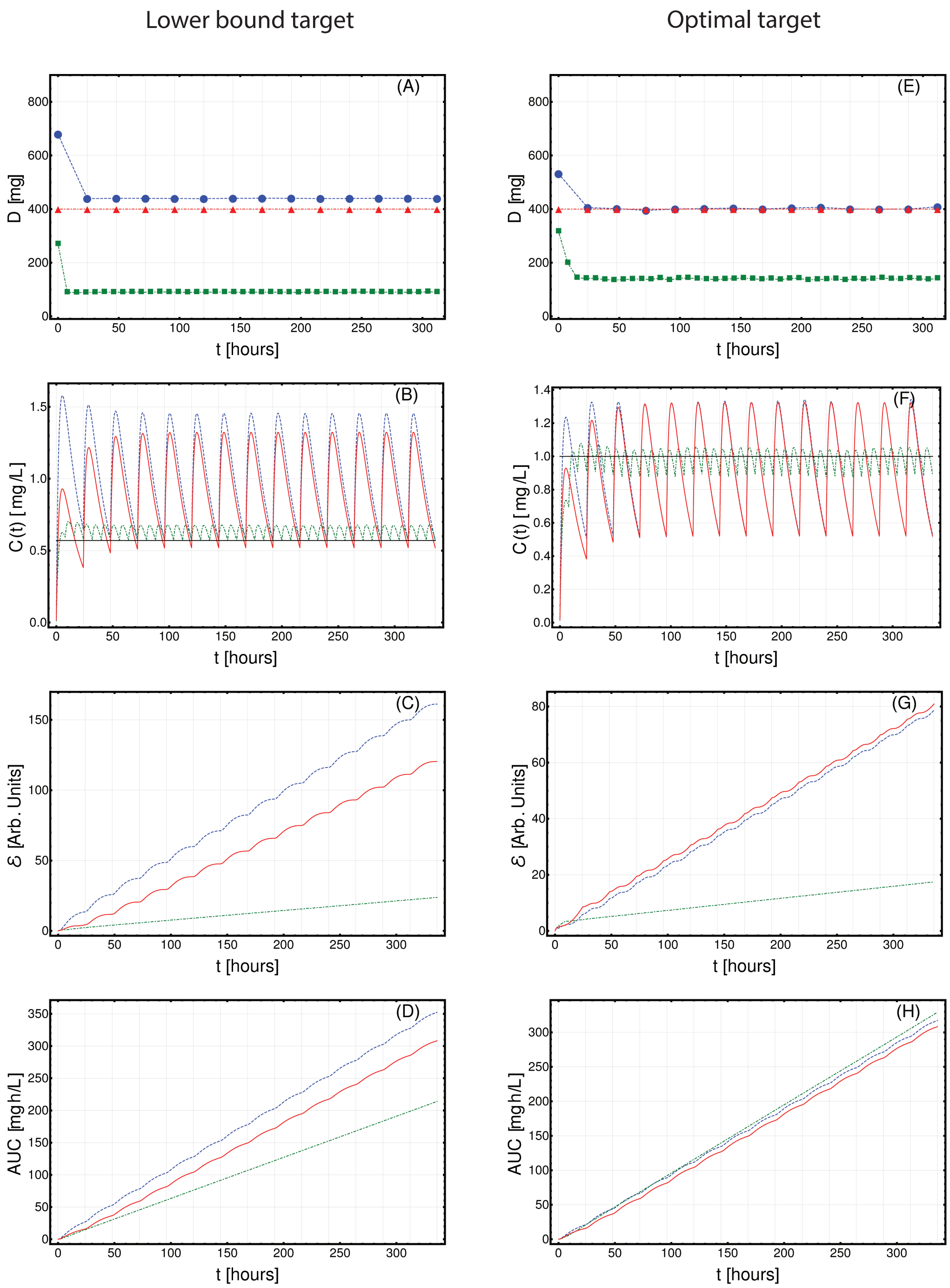


**Fig. S13.** Optimized Imatinib administration returned by CT4TD for patient 0002 00003 SMV from (20), in the cases of: 1-dose/day (blue) and 3-doses/day (green), with respect to: $lower\text{-}bound$ target concentration $C_{targ} = 0.57\ [mg/L]$ (left panels: A–D) and $optimal$ target concentration $C_{targ} = 1\ [mg/L]$ (right panels: E–H). Standard administration – i.e., 400 mg Imatinib/day – is shown with a red dashed line. In this case, the optimization of PK/PD models is obtained on patient-specific PK parameters, without considering the PD models. **(A,D)** Imatinib scheduled dosage in $mg$ (y axis), displayed on $14$ days (x axis). **(B-E)** Imatinib concentration in blood in $[mg/L]$ (y axis). **(C-F)** Variation of the cumulative distance between the observed concentration and the target in time. **(D-H)** The AUC in $[mg \cdot h/L]$

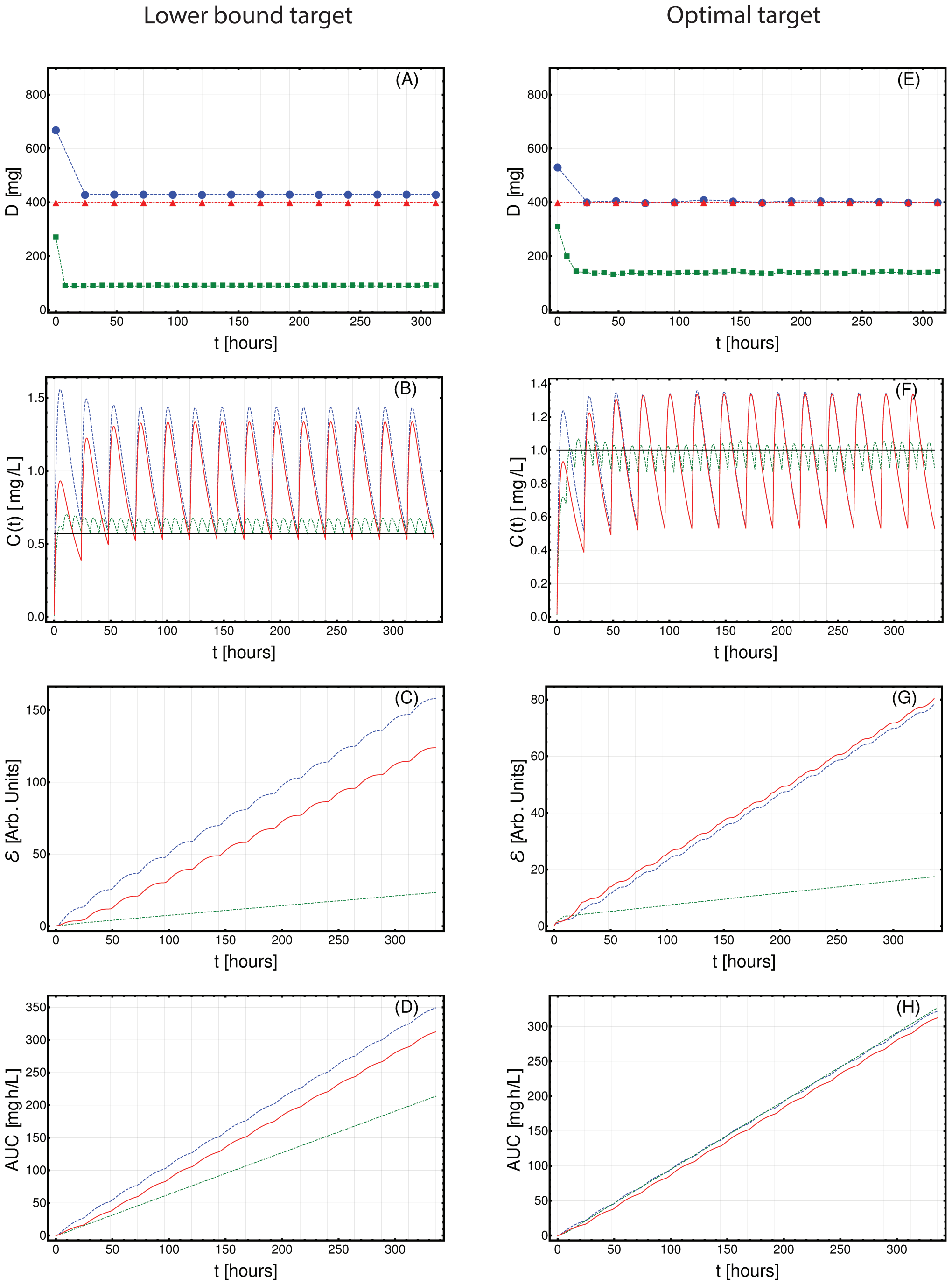


**Fig. S14.** Optimized Imatinib administration returned by CT4TD for patient 0002 00003 PMG from (20), in the cases of: 1-dose/day (blue) and 3-doses/day (green), with respect to: $lower\text{-}bound$ target concentration $C_{targ} = 0.57$ $[mg/L]$ (left panels: A–D) and $optimal$ target concentration $C_{targ} = 1$ $[mg/L]$ (right panels: E–H). Standard administration – i.e., 400 mg Imatinib/day – is shown with a red dashed line. In this case, the optimization of PK/PD models is obtained on patient-specific PK parameters, without considering the PD models. **(A,D)** Imatinib scheduled dosage in $mg$ (y axis), displayed on $14$ days (x axis). **(B-E)** Imatinib concentration in blood in $[mg/L]$ (y axis). **(C-F)** Variation of the cumulative distance between the observed concentration and the target in time. **(D-H)** The AUC in $[mg \cdot h/L]$

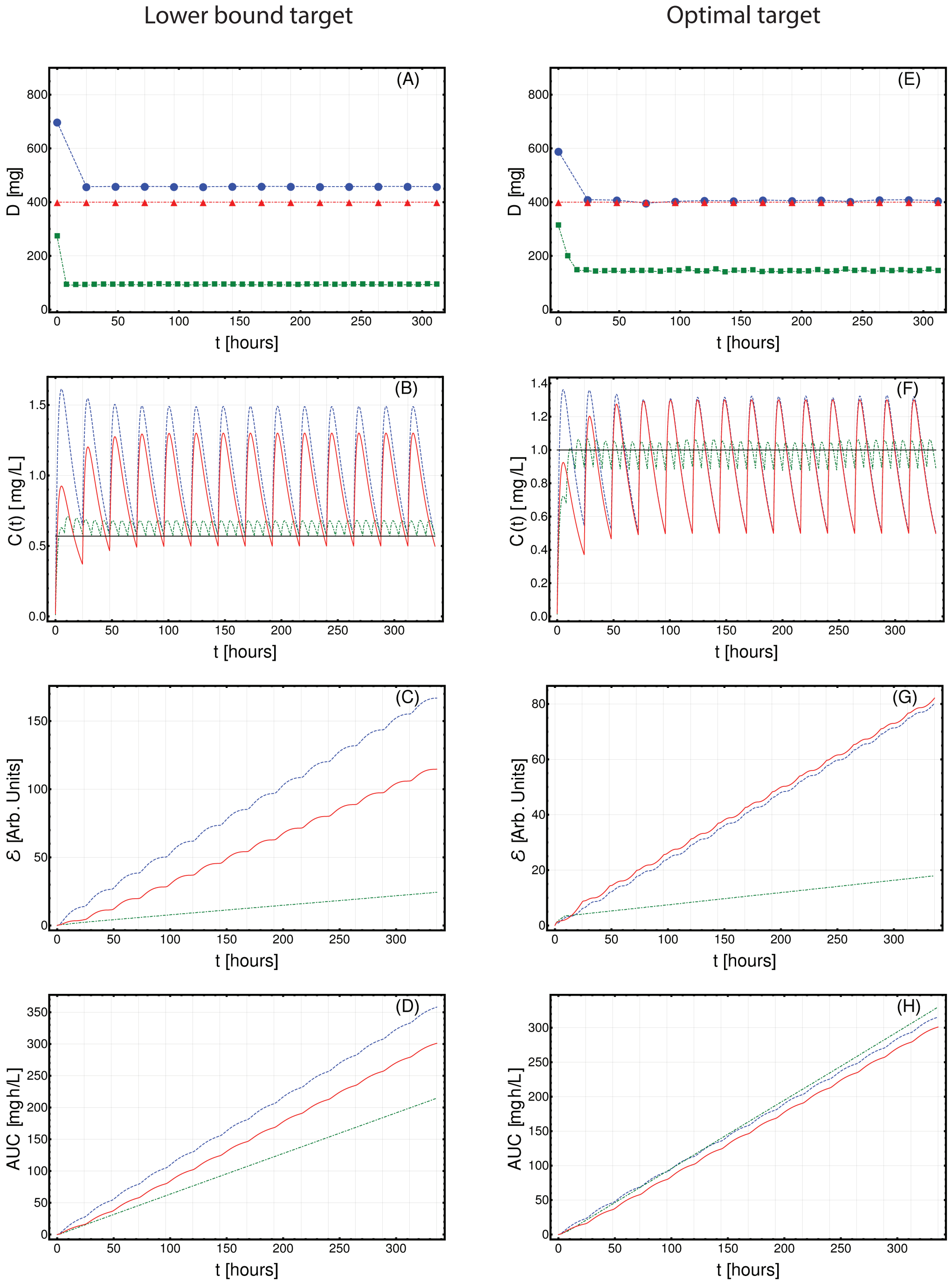


**Fig. S15.** Optimized Imatinib administration returned by CT4TD for patient 0002 00008 SAT from (20), in the cases of: 1-dose/day (blue) and 3-doses/day (green), with respect to: $lower\text{-}bound$ target concentration $C_{targ} = 0.57\ [mg/L]$ (left panels: A–D) and $optimal$ target concentration $C_{targ} = 1\ [mg/L]$ (right panels: E–H). Standard administration – i.e., 400 mg Imatinib/day – is shown with a red dashed line. In this case, the optimization of PK/PD models is obtained on patient-specific PK parameters, without considering the PD models. **(A,D)** Imatinib scheduled dosage in $mg$ (y axis), displayed on $14$ days (x axis). **(B-E)** Imatinib concentration in blood in $[mg/L]$ (y axis). **(C-F)** Variation of the cumulative distance between the observed concentration and the target in time.AF **(D-H)** The AUC in $[mg \cdot h/L]$

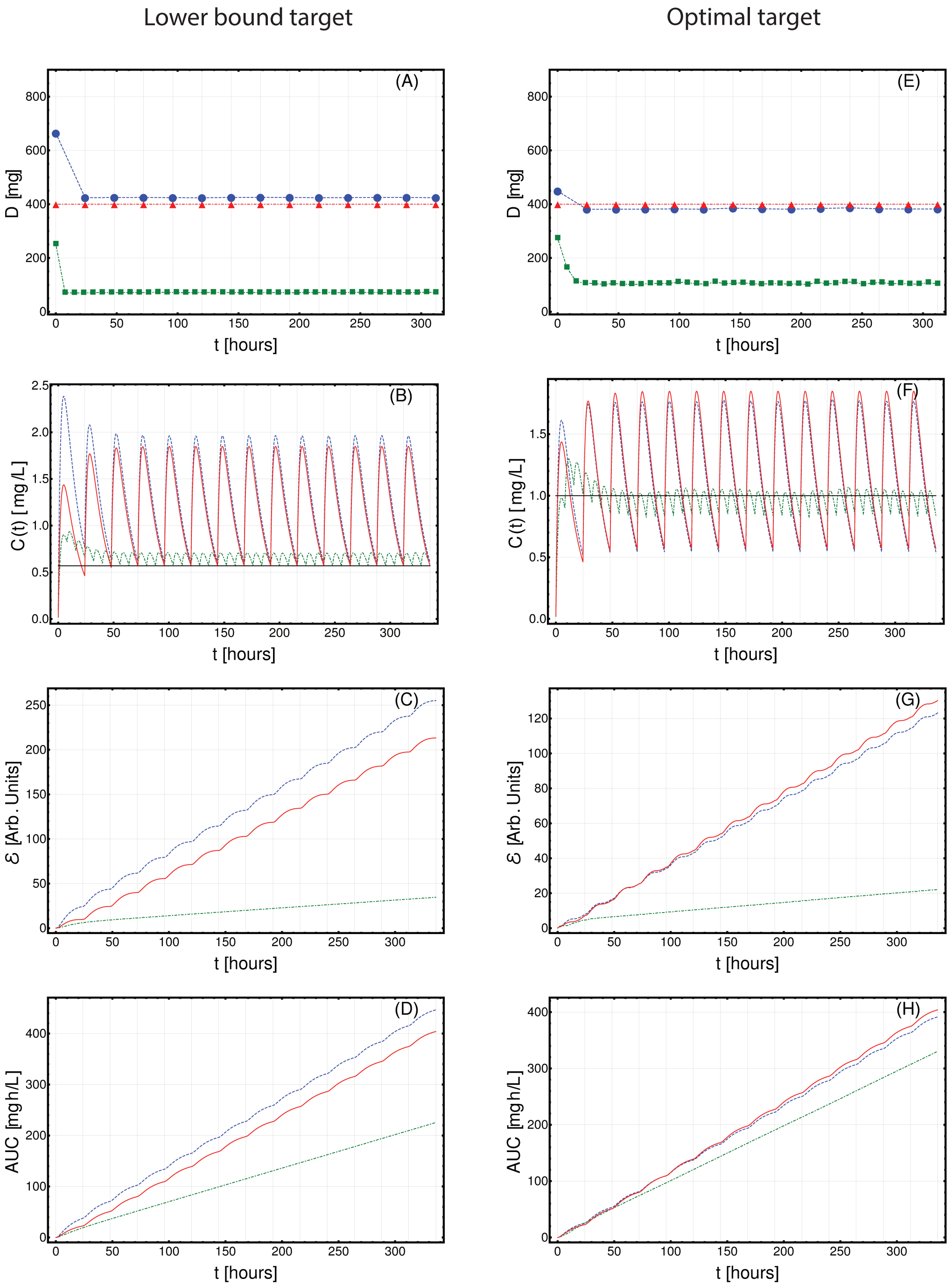


**Fig. S16.** Optimized Imatinib administration returned by CT4TD for patient 0003 00002 CL from (20), in the cases of: 1-dose/day (blue) and 3-doses/day (green), with respect to: $lower\text{-}bound$ target concentration $C_{targ} = 0.57$ $[mg/L]$ (left panels: A–D) and $optimal$ target concentration $C_{targ} = 1$ $[mg/L]$ (right panels: E–H). Standard administration – i.e., 400 mg Imatinib/day – is shown with a red dashed line. In this case, the optimization of PK/PD models is obtained on patient-specific PK parameters, without considering the PD models. **(A,D)** Imatinib scheduled dosage in $mg$ (y axis), displayed on $14$ days (x axis). **(B-E)** Imatinib concentration in blood in $[mg/L]$ (y axis). **(C-F)** Variation of the cumulative distance between the observed concentration and the target in time.**(D-H)** The AUC in $[mg \cdot h/L]$

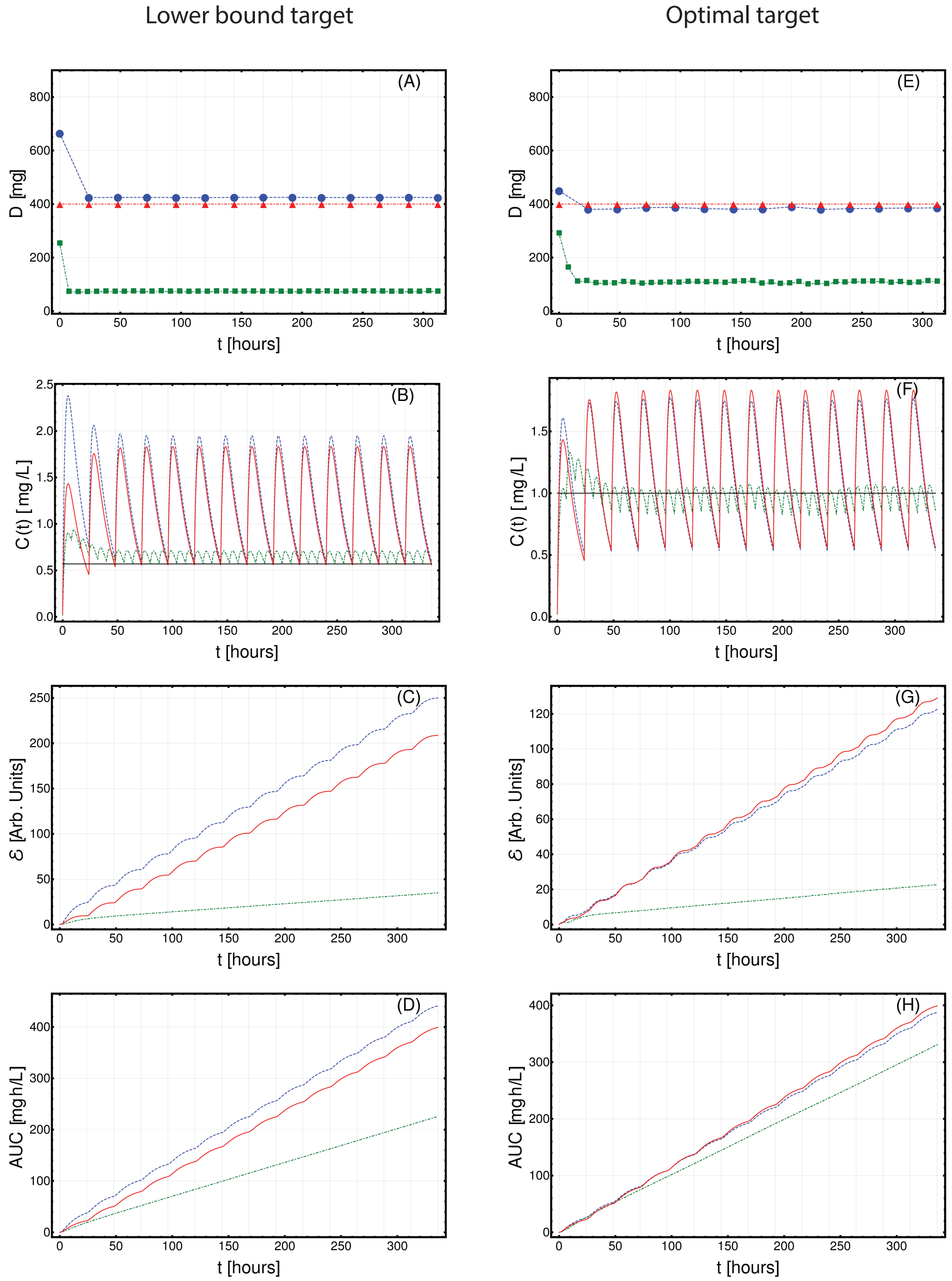


**Fig. S17.** Optimized Imatinib administration returned by CT4TD for patient 0004 00003 CAR from (20), in the cases of: 1-dose/day (blue) and 3-doses/day (green), with respect to: $lower\text{-}bound$ target concentration $C_{targ} = 0.57\ [mg/L]$ (left panels: A–D) and $optimal$ target concentration $C_{targ} = 1\ [mg/L]$ (right panels: E–H). Standard administration – i.e., 400 mg Imatinib/day – is shown with a red dashed line. In this case, the optimization of PK/PD models is obtained on patient-specific PK parameters, without considering the PD models. **(A,D)** Imatinib scheduled dosage in $mg$ (y axis), displayed on $14$ days (x axis). **(B-E)** Imatinib concentration in blood in $[mg/L]$ (y axis). **(C-F)** Variation of the cumulative distance between the observed concentration and the target in time.**(D-H)** The AUC in $[mg \cdot h/L]$

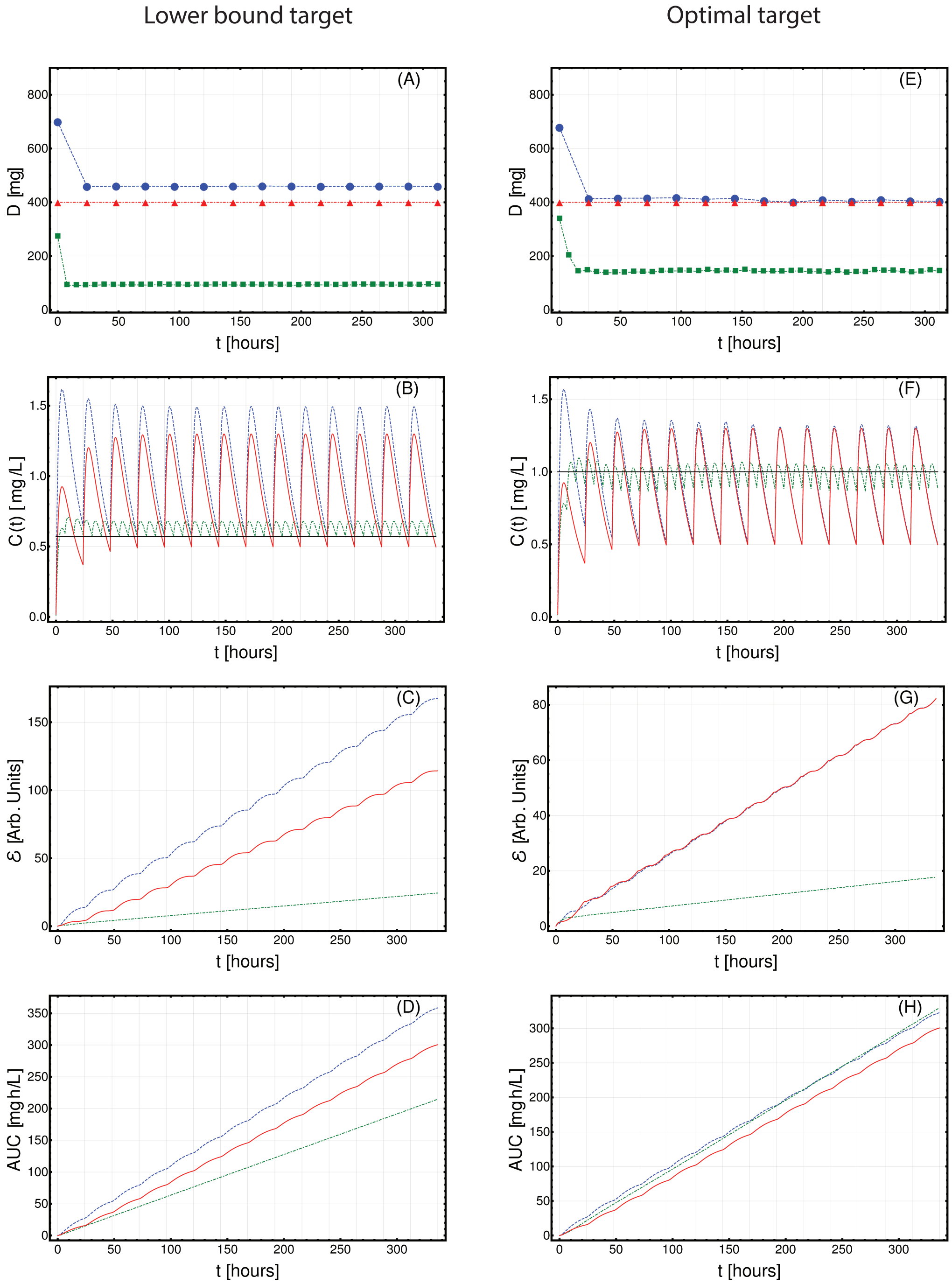


**Fig. S18.** Optimized Imatinib administration returned by CT4TD for patient 0004 00006 GM from (20), in the cases of: 1-dose/day (blue) and 3-doses/day (green), with respect to: $lower\text{-}bound$ target concentration $C_{targ} = 0.57\ [mg/L]$ (left panels: A–D) and $optimal$ target concentration $C_{targ} = 1\ [mg/L]$ (right panels: E–H). Standard administration – i.e., 400 mg Imatinib/day – is shown with a red dashed line. In this case, the optimization of PK/PD models is obtained on patient-specific PK parameters, without considering the PD models. **(A,D)** Imatinib scheduled dosage in $mg$ (y axis), displayed on $14$ days (x axis). **(B-E)** Imatinib concentration in blood in $[mg/L]$ (y axis). **(C-F)** Variation of the cumulative distance between the observed concentration and the target in time.**(D-H)** The AUC in $[mg \cdot h/L]$

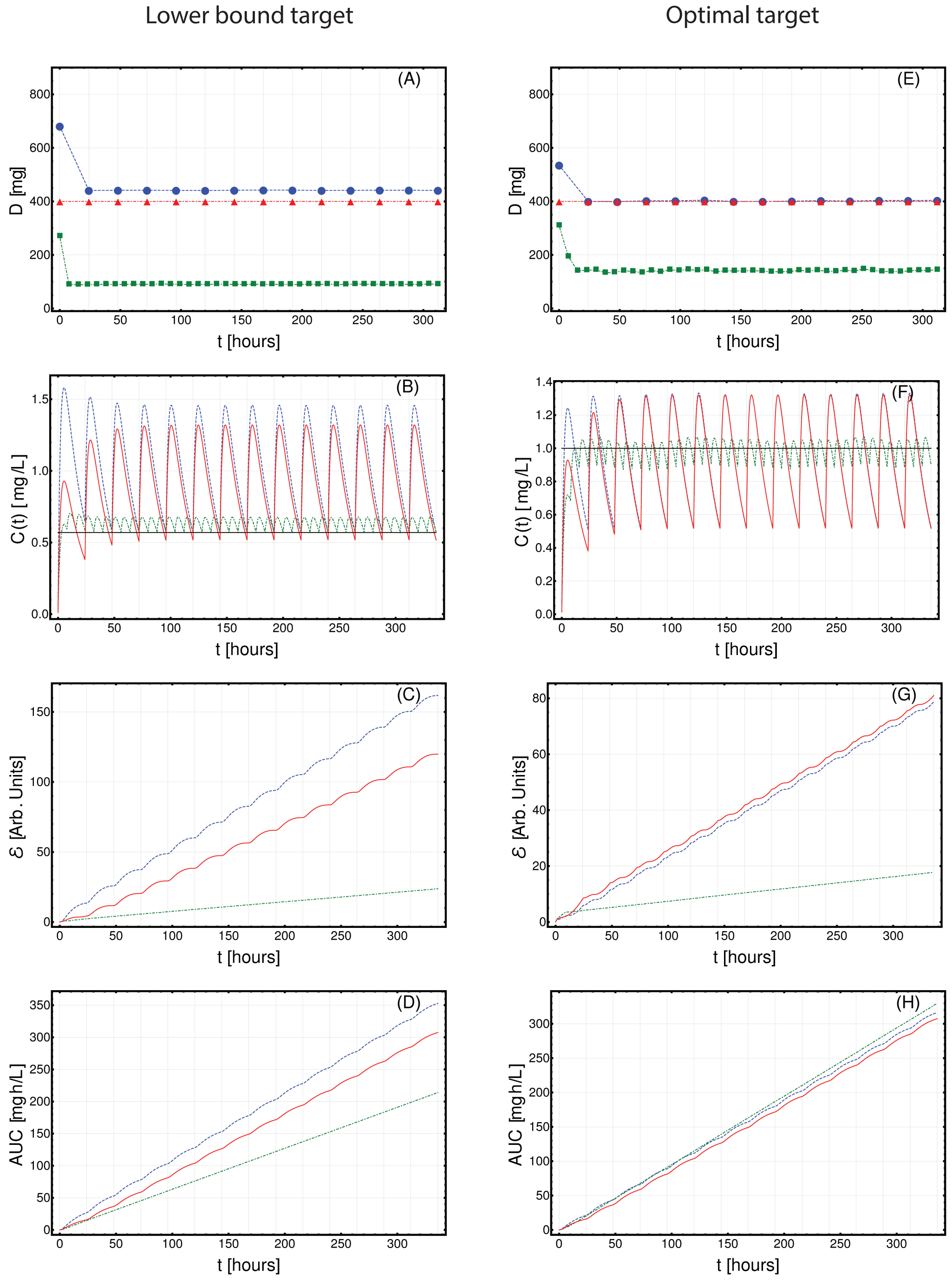


**Fig. S19.** Optimized Imatinib administration returned by CT4TD for patient 0006 00006 ML from (20), in the cases of: 1-dose/day (blue) and 3-doses/day (green), with respect to: $lower\text{-}bound$ target concentration $C_{targ} = 0.57\ [mg/L]$ (left panels: A–D) and $optimal$ target concentration $C_{targ} = 1\ [mg/L]$ (right panels: E–H). Standard administration – i.e., 400 mg Imatinib/day – is shown with a red dashed line. In this case, the optimization of PK/PD models is obtained on patient-specific PK parameters, without considering the PD models. **(A,D)** Imatinib scheduled dosage in $mg$ (y axis), displayed on $14$ days (x axis). **(B-E)** Imatinib concentration in blood in $[mg/L]$ (y axis). **(C-F)** Variation of the cumulative distance between the observed concentration and the target in time.**(D-H)** The AUC in $[mg \cdot h/L]$

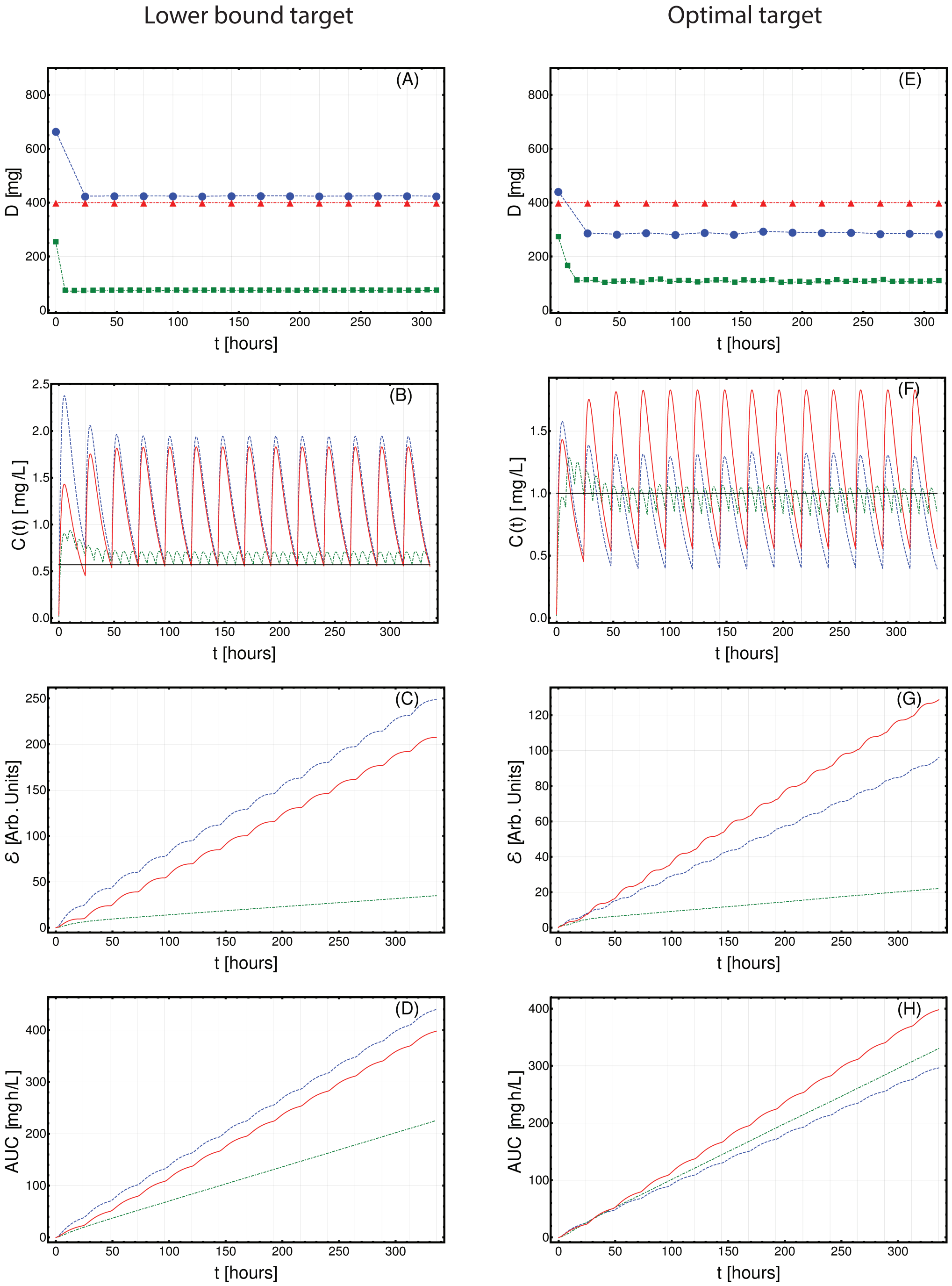


**Fig. S20.** Optimized Imatinib administration returned by CT4TD for patient 0006 00007 RJW from (20), in the cases of: 1-dose/day (blue) and 3-doses/day (green), with respect to: $lower\text{-}bound$ target concentration $C_{targ} = 0.57\ [mg/L]$ (left panels: A–D) and $optimal$ target concentration $C_{targ} = 1\ [mg/L]$ (right panels: E–H). Standard administration – i.e., 400 mg Imatinib/day – is shown with a red dashed line. In this case, the optimization of PK/PD models is obtained on patient-specific PK parameters, without considering the PD models. **(A,D)** Imatinib scheduled dosage in $mg$ (y axis), displayed on $14$ days (x axis). **(B-E)** Imatinib concentration in blood in $[mg/L]$ (y axis). **(C-F)** Variation of the cumulative distance between the observed concentration and the target in time.**(D-H)** The AUC in $[mg \cdot h/L]$

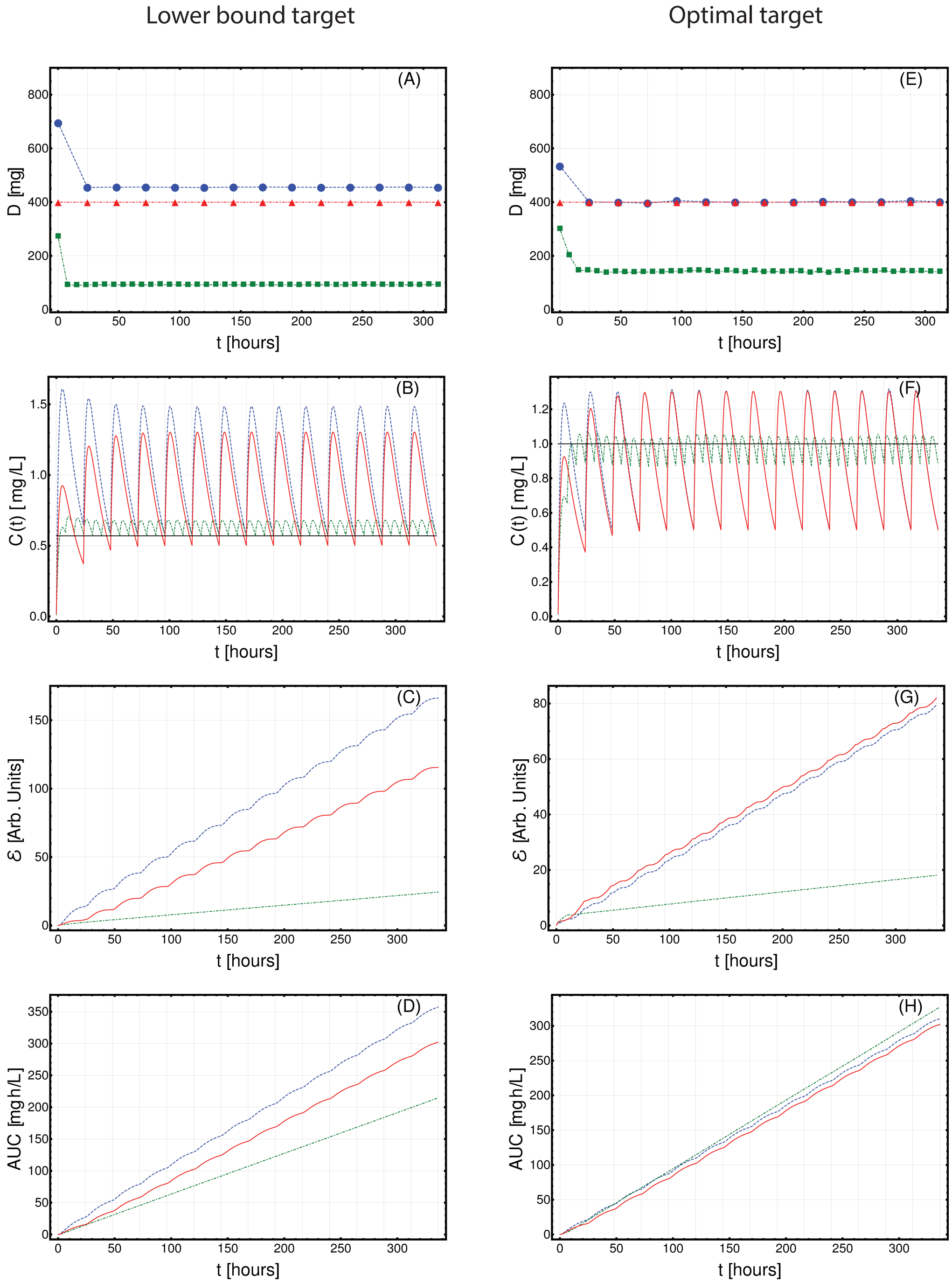


**Fig. S21.** Optimized Imatinib administration returned by CT4TD for patient 0007 00001 AJV from (20), in the cases of: 1-dose/day (blue) and 3-doses/day (green), with respect to: $lower\text{-}bound$ target concentration $C_{targ} = 0.57$ $[mg/L]$ (left panels: A–D) and $optimal$ target concentration $C_{targ} = 1$ $[mg/L]$ (right panels: E–H). Standard administration – i.e., 400 mg Imatinib/day – is shown with a red dashed line. In this case, the optimization of PK/PD models is obtained on patient-specific PK parameters, without considering the PD models. **(A,D)** Imatinib scheduled dosage in $mg$ (y axis), displayed on $14$ days (x axis). **(B-E)** Imatinib concentration in blood in $[mg/L]$ (y axis). **(C-F)** Variation of the cumulative distance between the observed concentration and the target in time.**(D-H)** The AUC in $[mg \cdot h/L]$

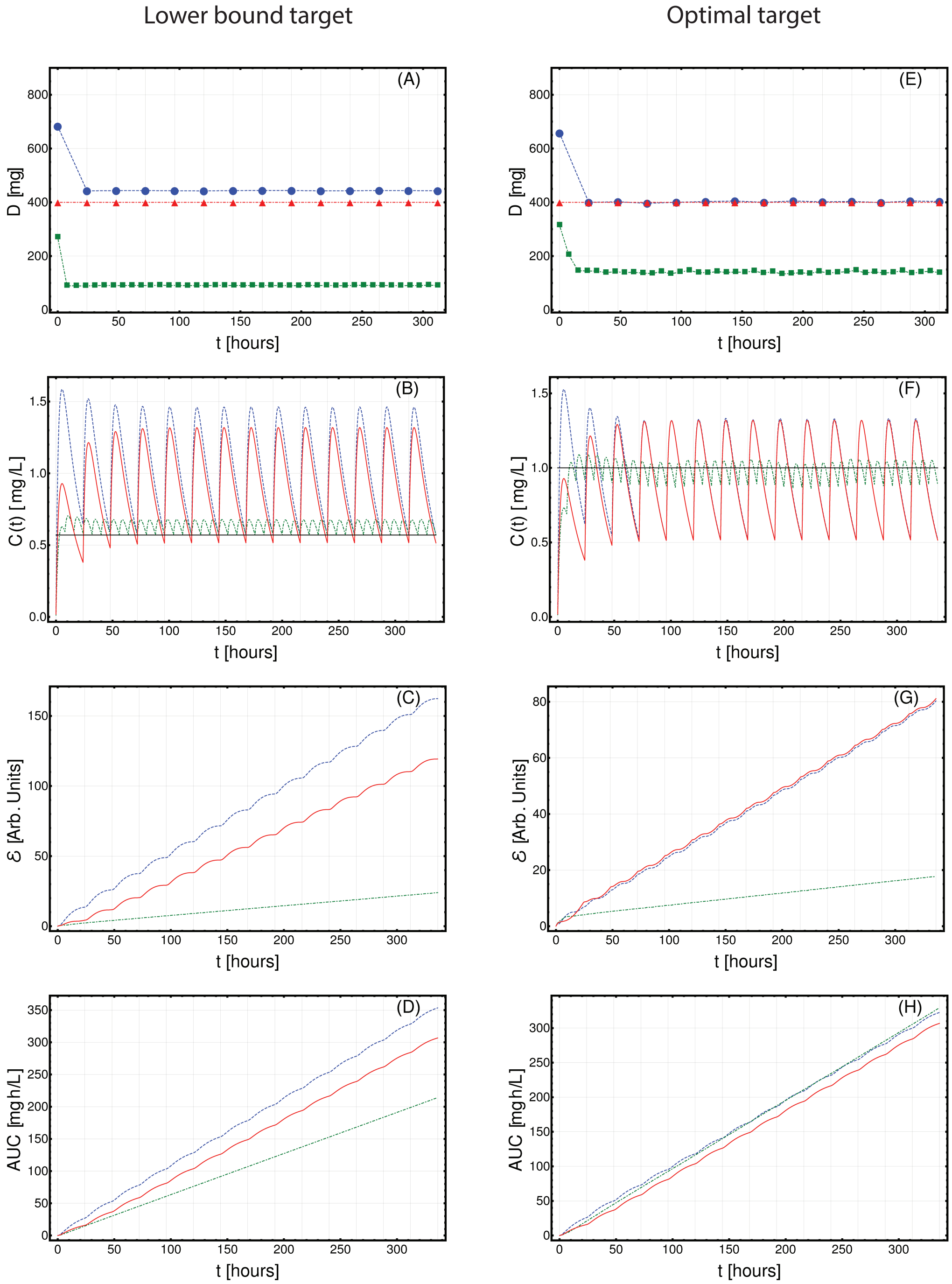


**Fig. S22.** Optimized Imatinib administration returned by CT4TD for patient 0007 00002 DPS from (20), in the cases of: 1-dose/day (blue) and 3-doses/day (green), with respect to: $lower\text{-}bound$ target concentration $C_{targ} = 0.57$ $[mg/L]$ (left panels: A–D) and $optimal$ target concentration $C_{targ} = 1$ $[mg/L]$ (right panels: E–H). Standard administration – i.e., 400 mg Imatinib/day – is shown with a red dashed line. In this case, the optimization of PK/PD models is obtained on patient-specific PK parameters, without considering the PD models. **(A,D)** Imatinib scheduled dosage in $mg$ (y axis), displayed on $14$ days (x axis). **(B-E)** Imatinib concentration in blood in $[mg/L]$ (y axis). **(C-F)** Variation of the cumulative distance between the observed concentration and the target in time. **(D-H)** The AUC in $[mg \cdot h/L]$

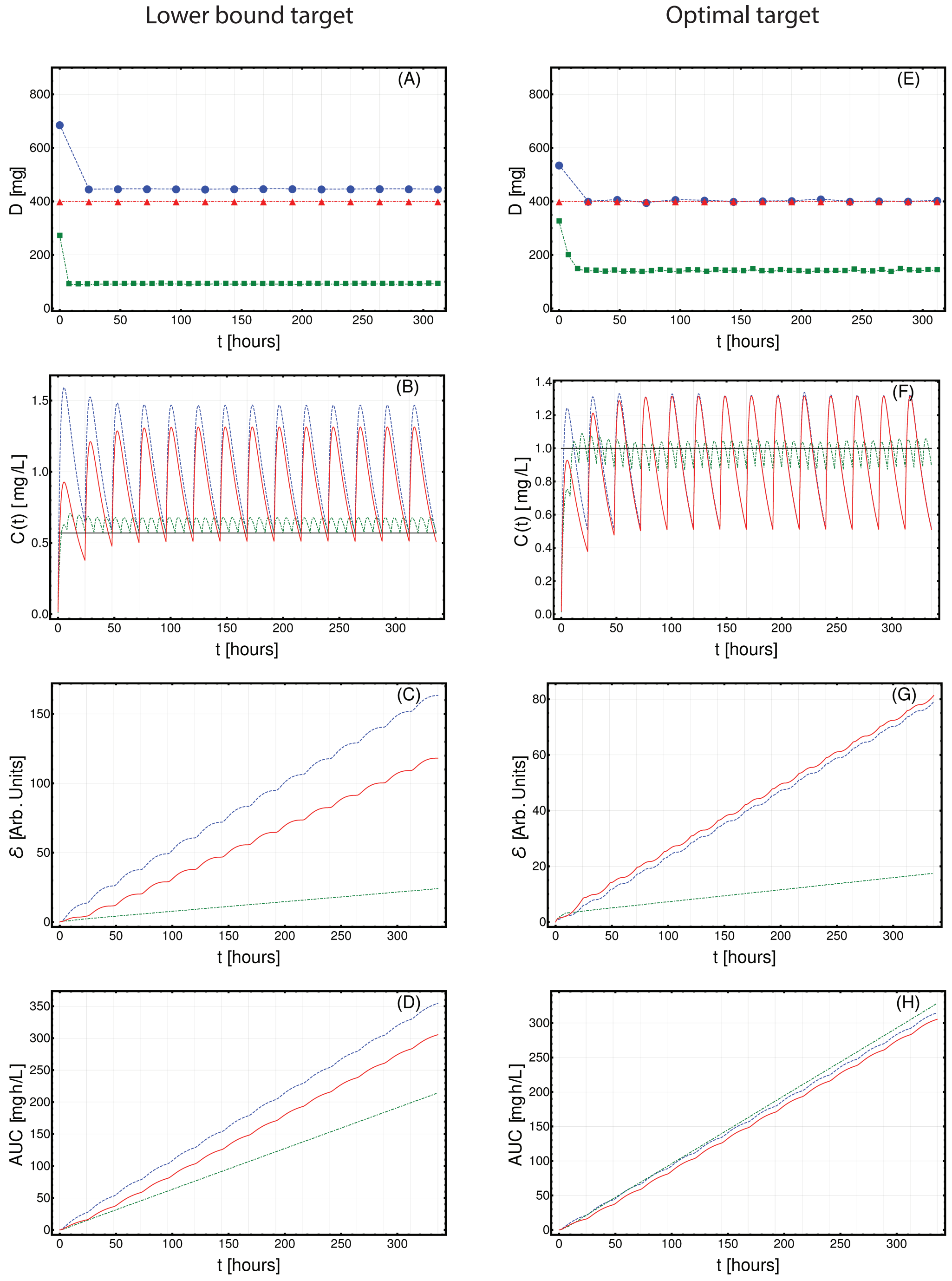


**Fig. S23.** Optimized Imatinib administration returned by CT4TD for patient 0008 00002 PWR from (20), in the cases of: 1-dose/day (blue) and 3-doses/day (green), with respect to: $lower\text{-}bound$ target concentration $C_{targ} = 0.57\ [mg/L]$ (left panels: A–D) and $optimal$ target concentration $C_{targ} = 1\ [mg/L]$ (right panels: E–H). Standard administration – i.e., 400 mg Imatinib/day – is shown with a red dashed line. In this case, the optimization of PK/PD models is obtained on patient-specific PK parameters, without considering the PD models. **(A,D)** Imatinib scheduled dosage in $mg$ (y axis), displayed on $14$ days (x axis). **(B-E)** Imatinib concentration in blood in $[mg/L]$ (y axis). **(C-F)** Variation of the cumulative distance between the observed concentration and the target in time.**(D-H)** The AUC in $[mg \cdot h/L]$

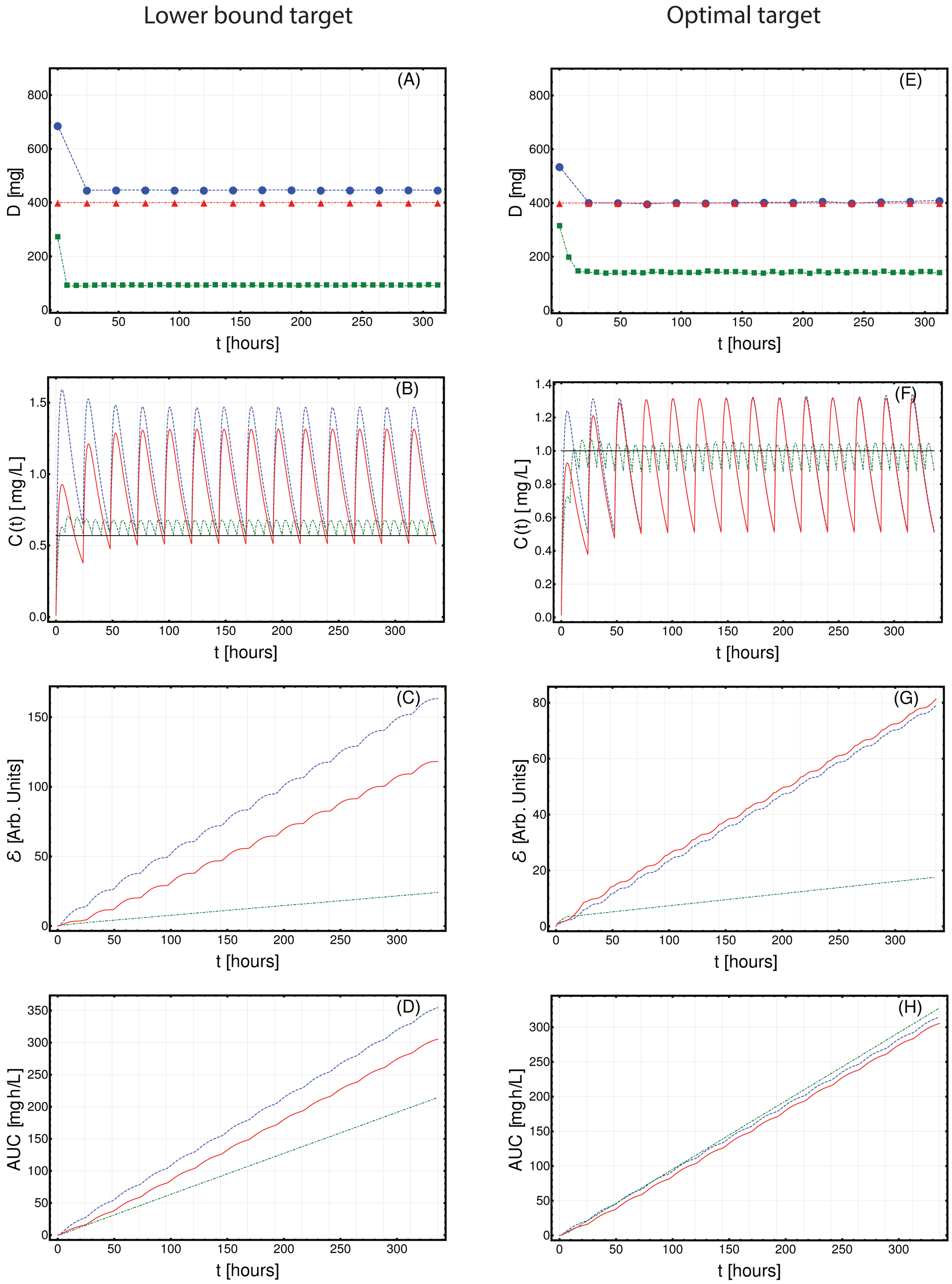


**Fig. S24.** Optimized Imatinib administration returned by CT4TD for patient 0008 00003 LJG from (20), in the cases of: 1-dose/day (blue) and 3-doses/day (green), with respect to: $lower\text{-}bound$ target concentration $C_{targ} = 0.57\ [mg/L]$ (left panels: A–D) and $optimal$ target concentration $C_{targ} = 1\ [mg/L]$ (right panels: E–H). Standard administration – i.e., 400 mg Imatinib/day – is shown with a red dashed line. In this case, the optimization of PK/PD models is obtained on patient-specific PK parameters, without considering the PD models. **(A,D)** Imatinib scheduled dosage in $mg$ (y axis), displayed on $14$ days (x axis). **(B-E)** Imatinib concentration in blood in $[mg/L]$ (y axis). **(C-F)** Variation of the cumulative distance between the observed concentration and the target in time.**(D-H)** The AUC in $[mg \cdot h/L]$

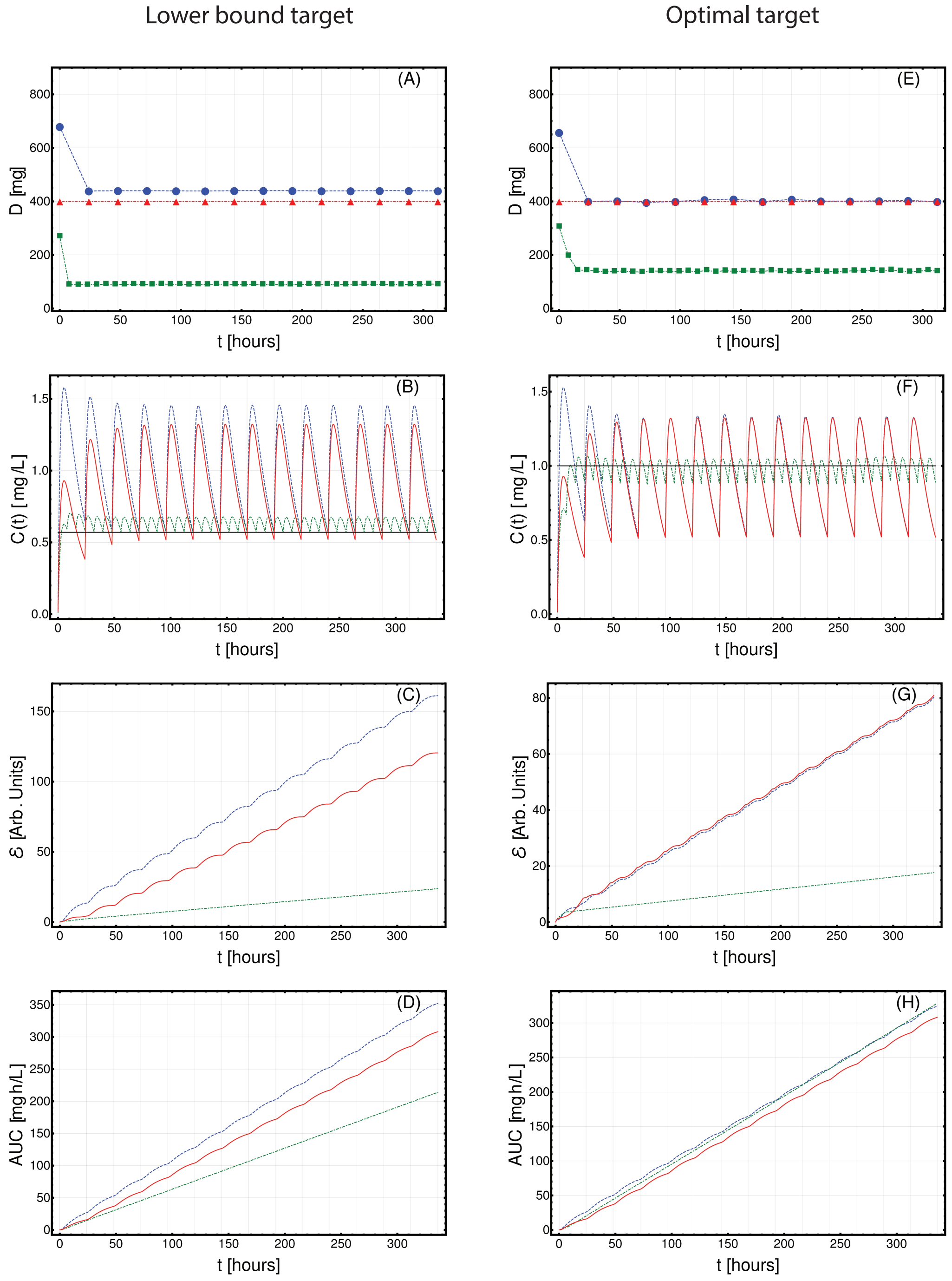


**Fig. S25.** Optimized Imatinib administration returned by CT4TD for patient 0008 00005 BAT from (20), in the cases of: 1-dose/day (blue) and 3-doses/day (green), with respect to: $lower\text{-}bound$ target concentration $C_{targ} = 0.57\ [mg/L]$ (left panels: A–D) and $optimal$ target concentration $C_{targ} = 1\ [mg/L]$ (right panels: E–H). Standard administration – i.e., 400 mg Imatinib/day – is shown with a red dashed line. In this case, the optimization of PK/PD models is obtained on patient-specific PK parameters, without considering the PD models. **(A,D)** Imatinib scheduled dosage in $mg$ (y axis), displayed on $14$ days (x axis). **(B-E)** Imatinib concentration in blood in $[mg/L]$ (y axis). **(C-F)** Variation of the cumulative distance between the observed concentration and the target in time.**(D-H)** The AUC in $[mg \cdot h/L]$

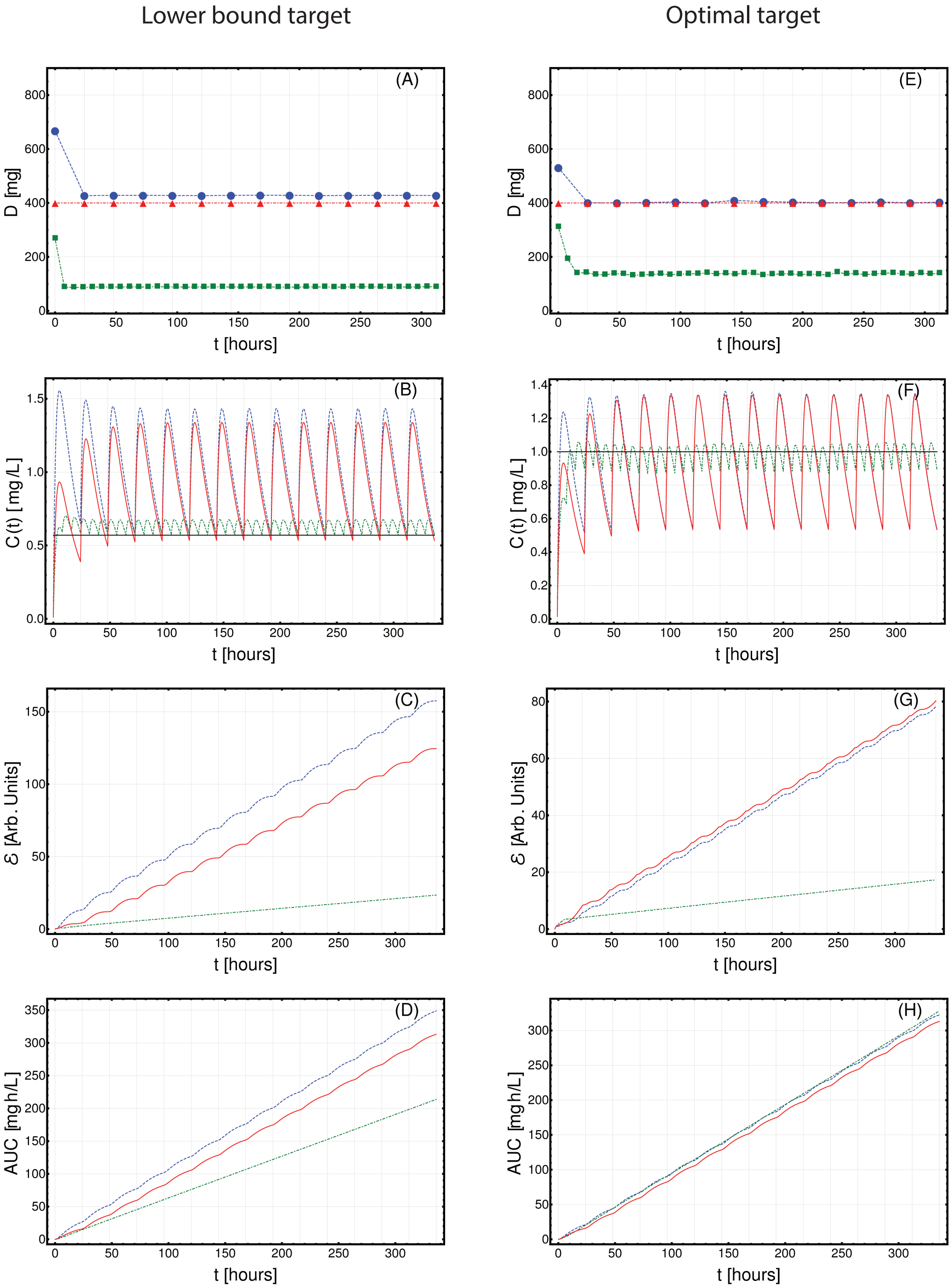


**Fig. S26.** Optimized Imatinib administration returned by CT4TD for patient 0009 00003 LH from (20), in the cases of: 1-dose/day (blue) and 3-doses/day (green), with respect to: $lower\text{-}bound$ target concentration $C_{targ} = 0.57\ [mg/L]$ (left panels: A–D) and $optimal$ target concentration $C_{targ} = 1\ [mg/L]$ (right panels: E–H). Standard administration – i.e., 400 mg Imatinib/day – is shown with a red dashed line. In this case, the optimization of PK/PD models is obtained on patient-specific PK parameters, without considering the PD models. **(A,D)** Imatinib scheduled dosage in $mg$ (y axis), displayed on $14$ days (x axis). **(B-E)** Imatinib concentration in blood in $[mg/L]$ (y axis). **(C-F)** Variation of the cumulative distance between the observed concentration and the target in time.**(D-H)** The AUC in $[mg \cdot h/L]$

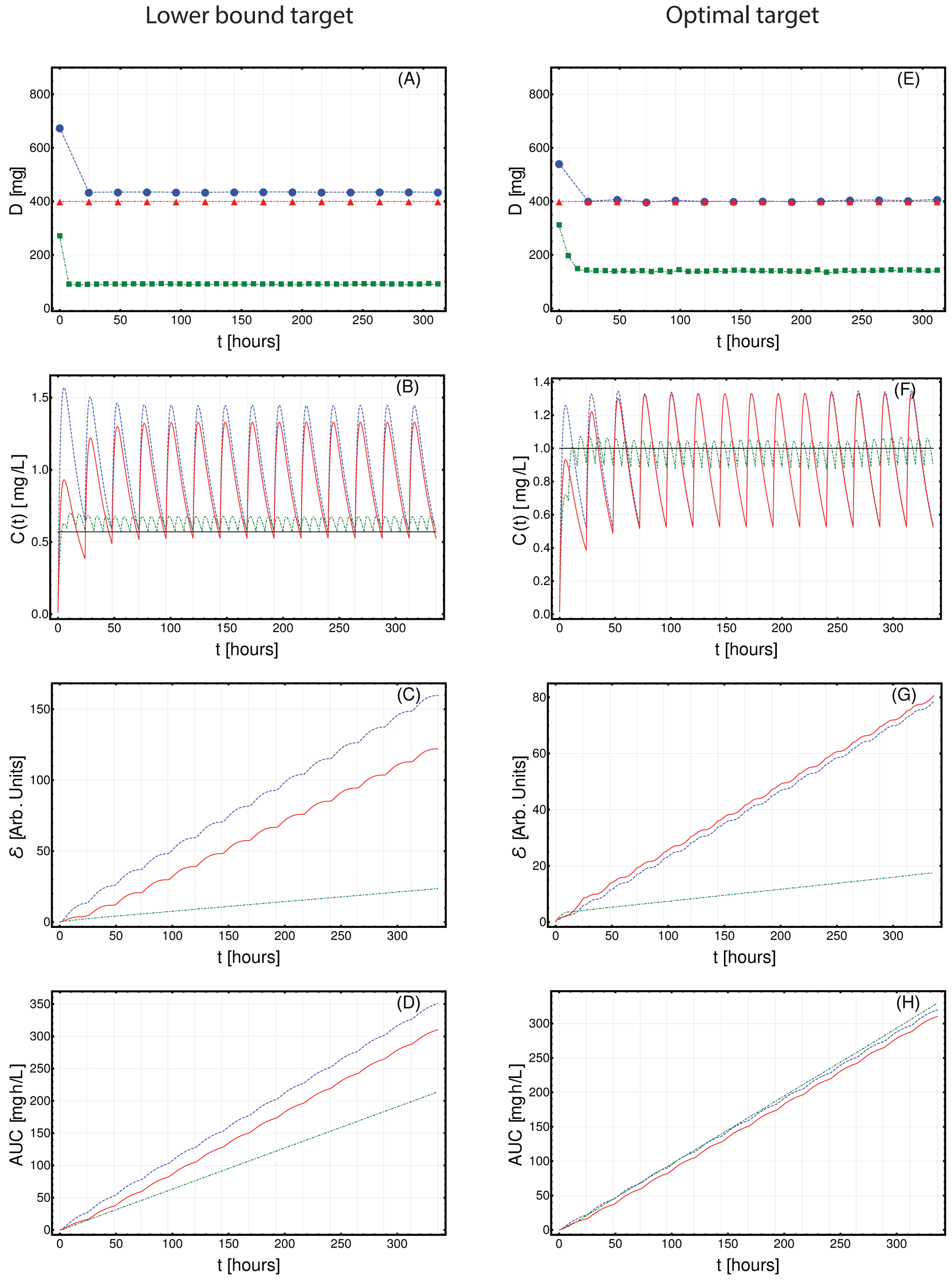


**Fig. S27.** Optimized Imatinib administration returned by CT4TD for patient 0010 00001 HJ from (20), in the cases of: 1-dose/day (blue) and 3-doses/day (green), with respect to: $lower\text{-}bound$ target concentration $C_{targ} = 0.57$ $[mg/L]$ (left panels: A–D) and $optimal$ target concentration $C_{targ} = 1$ $[mg/L]$ (right panels: E–H). Standard administration – i.e., 400 mg Imatinib/day – is shown with a red dashed line. In this case, the optimization of PK/PD models is obtained on patient-specific PK parameters, without considering the PD models. **(A,D)** Imatinib scheduled dosage in $mg$ (y axis), displayed on $14$ days (x axis). **(B-E)** Imatinib concentration in blood in $[mg/L]$ (y axis). **(C-F)** Variation of the cumulative distance between the observed concentration and the target in time.**(D-H)** The AUC in $[mg \cdot h/L]$

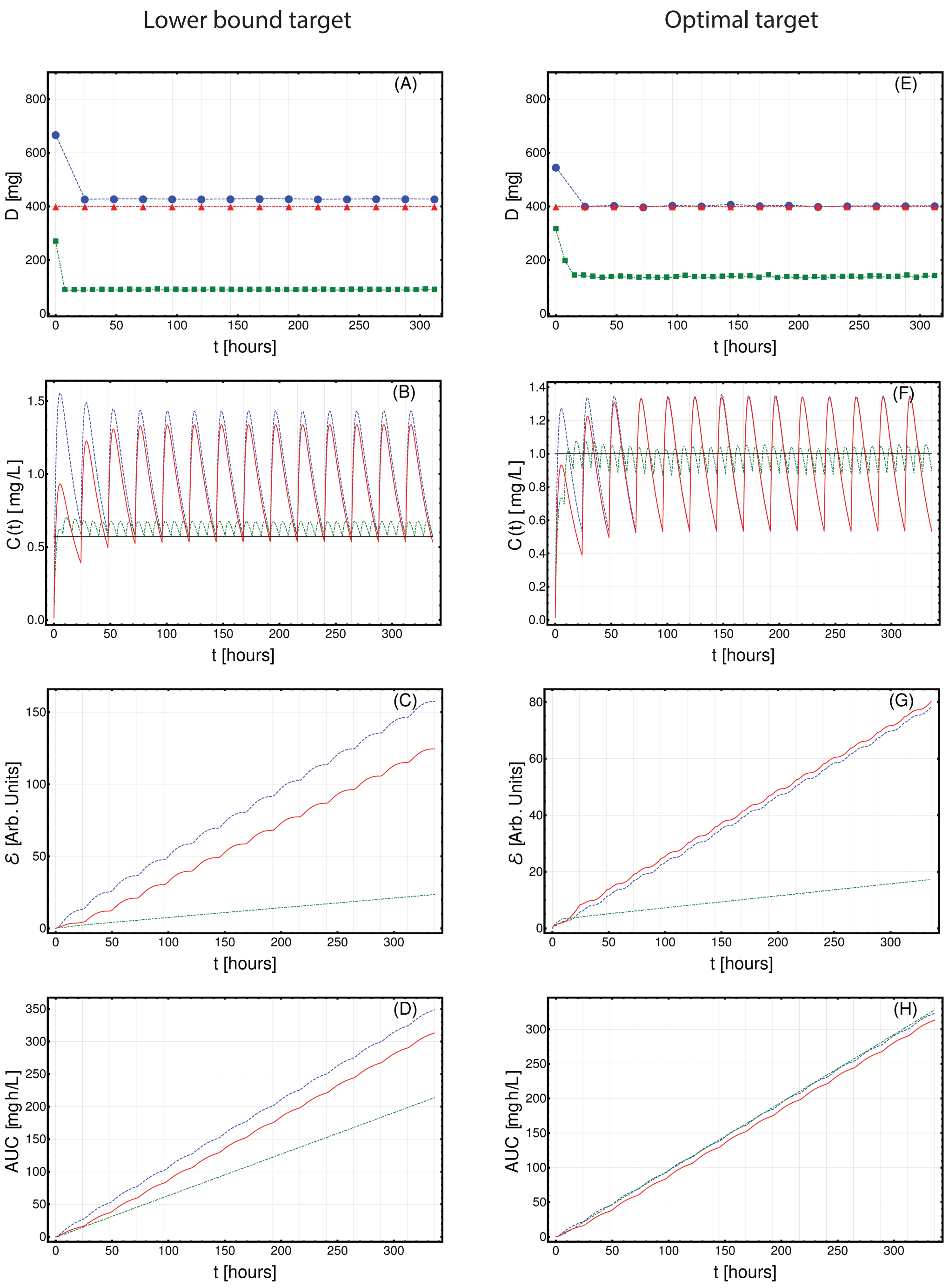


**Fig. S28.** Optimized Imatinib administration returned by CT4TD for patient 0010 00002 GR from (20), in the cases of: 1-dose/day (blue) and 3-doses/day (green), with respect to: $lower\text{-}bound$ target concentration $C_{targ} = 0.57\ [mg/L]$ (left panels: A–D) and $optimal$ target concentration $C_{targ} = 1\ [mg/L]$ (right panels: E–H). Standard administration – i.e., 400 mg Imatinib/day – is shown with a red dashed line. In this case, the optimization of PK/PD models is obtained on patient-specific PK parameters, without considering the PD models. **(A,D)** Imatinib scheduled dosage in $mg$ (y axis), displayed on $14$ days (x axis). **(B-E)** Imatinib concentration in blood in $[mg/L]$ (y axis). **(C-F)** Variation of the cumulative distance between the observed concentration and the target in time.**(D-H)** The AUC in $[mg \cdot h/L]$

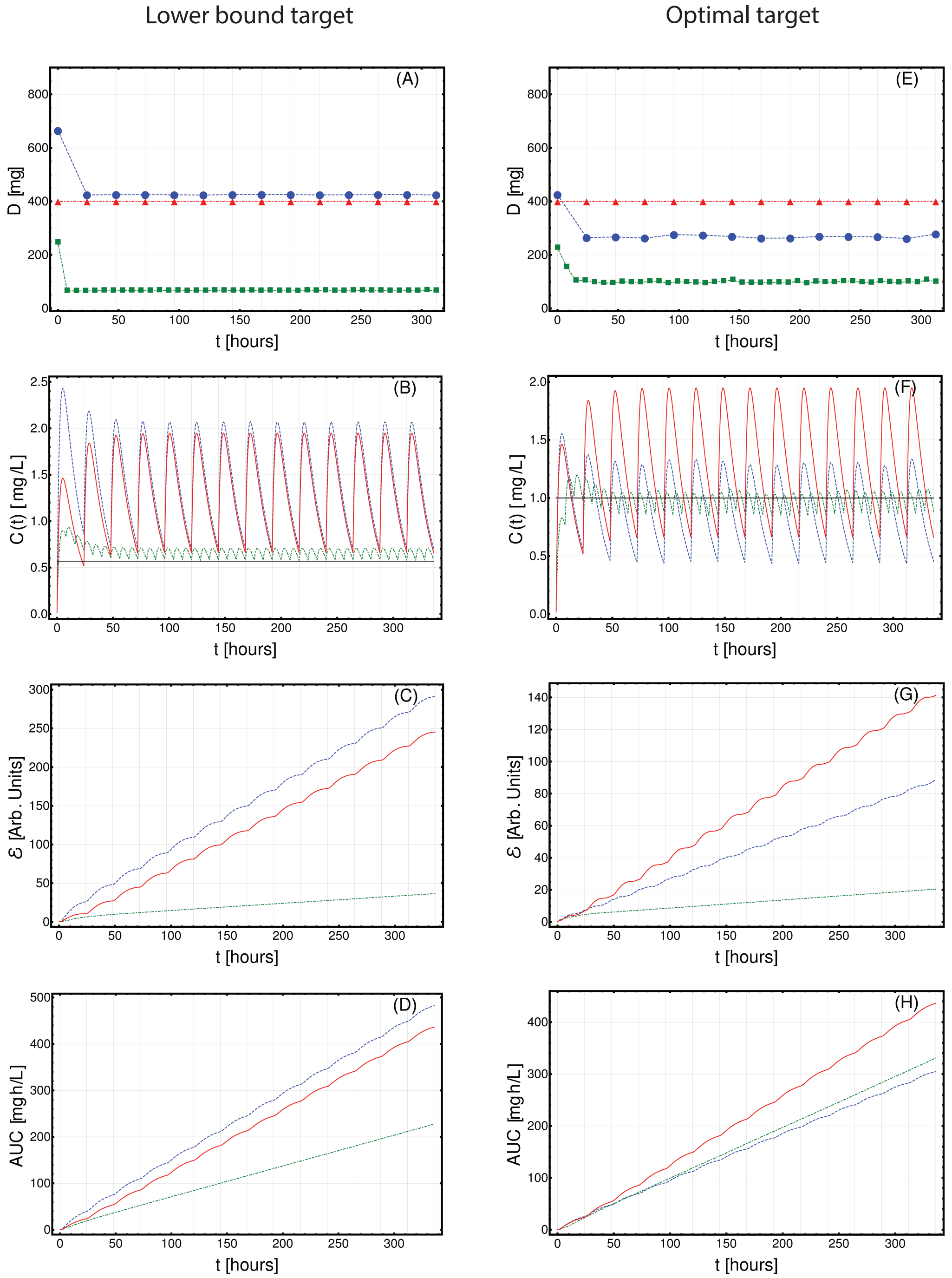


**Fig. S29.** Optimized Imatinib administration returned by CT4TD for patient 0011 00001 CMC from (20), in the cases of: 1-dose/day (blue) and 3-doses/day (green), with respect to: $lower\text{-}bound$ target concentration $C_{targ} = 0.57$ $[mg/L]$ (left panels: A–D) and $optimal$ target concentration $C_{targ} = 1$ $[mg/L]$ (right panels: E–H). Standard administration – i.e., 400 mg Imatinib/day – is shown with a red dashed line. In this case, the optimization of PK/PD models is obtained on patient-specific PK parameters, without considering the PD models. **(A,D)** Imatinib scheduled dosage in $mg$ (y axis), displayed on $14$ days (x axis). **(B-E)** Imatinib concentration in blood in $[mg/L]$ (y axis). **(C-F)** Variation of the cumulative distance between the observed concentration and the target in time.**(D-H)** The AUC in $[mg \cdot h/L]$

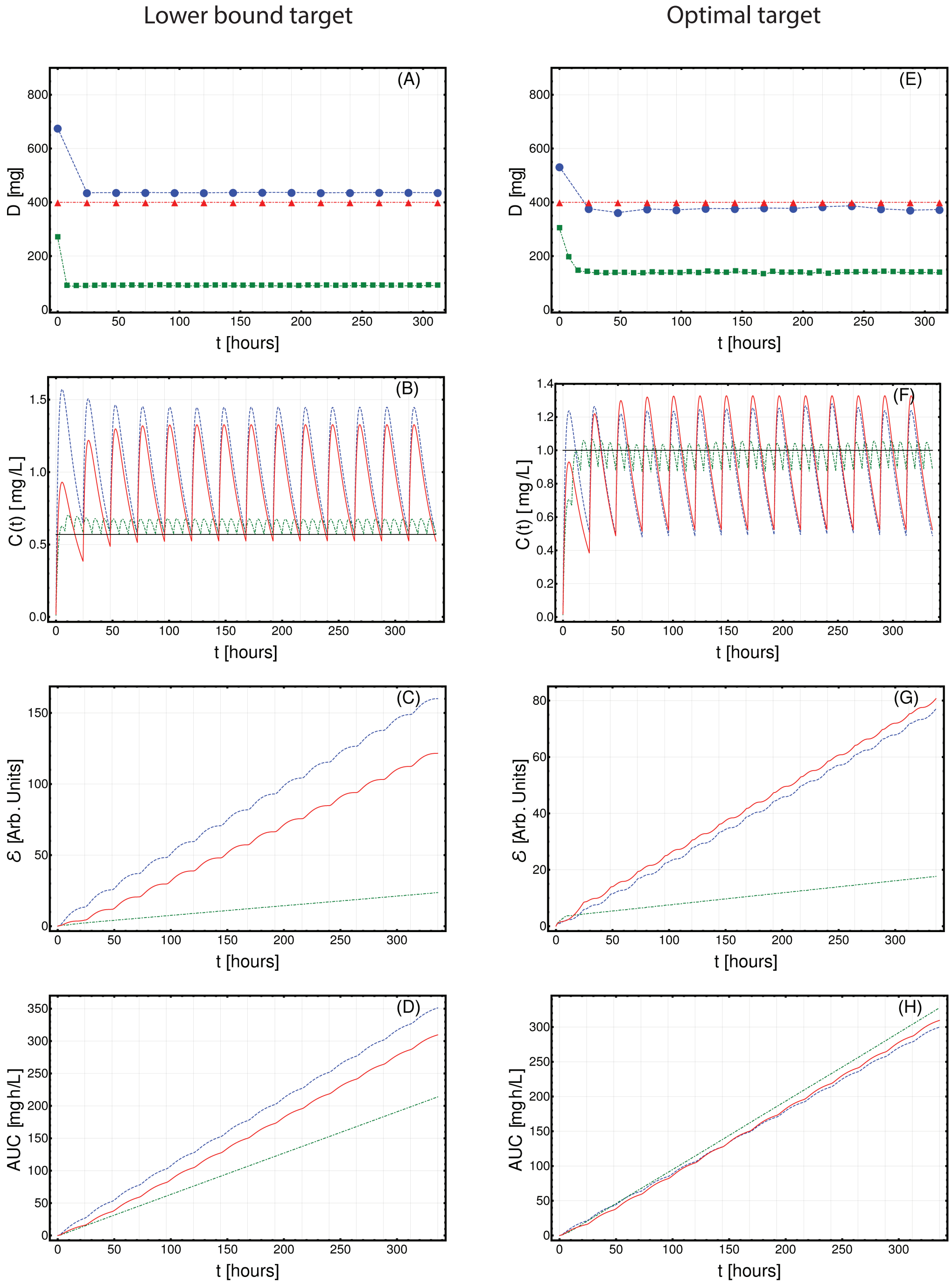


**Fig. S30.** Optimized Imatinib administration returned by CT4TD for patient 0011 00005 SCW from (20), in the cases of: 1-dose/day (blue) and 3-doses/day (green), with respect to: $lower\text{-}bound$ target concentration $C_{targ} = 0.57\ [mg/L]$ (left panels: A–D) and $optimal$ target concentration $C_{targ} = 1\ [mg/L]$ (right panels: E–H). Standard administration – i.e., 400 mg Imatinib/day – is shown with a red dashed line. In this case, the optimization of PK/PD models is obtained on patient-specific PK parameters, without considering the PD models. **(A,D)** Imatinib scheduled dosage in $mg$ (y axis), displayed on $14$ days (x axis). **(B-E)** Imatinib concentration in blood in $[mg/L]$ (y axis). **(C-F)** Variation of the cumulative distance between the observed concentration and the target in time.**(D-H)** The AUC in $[mg \cdot h/L]$

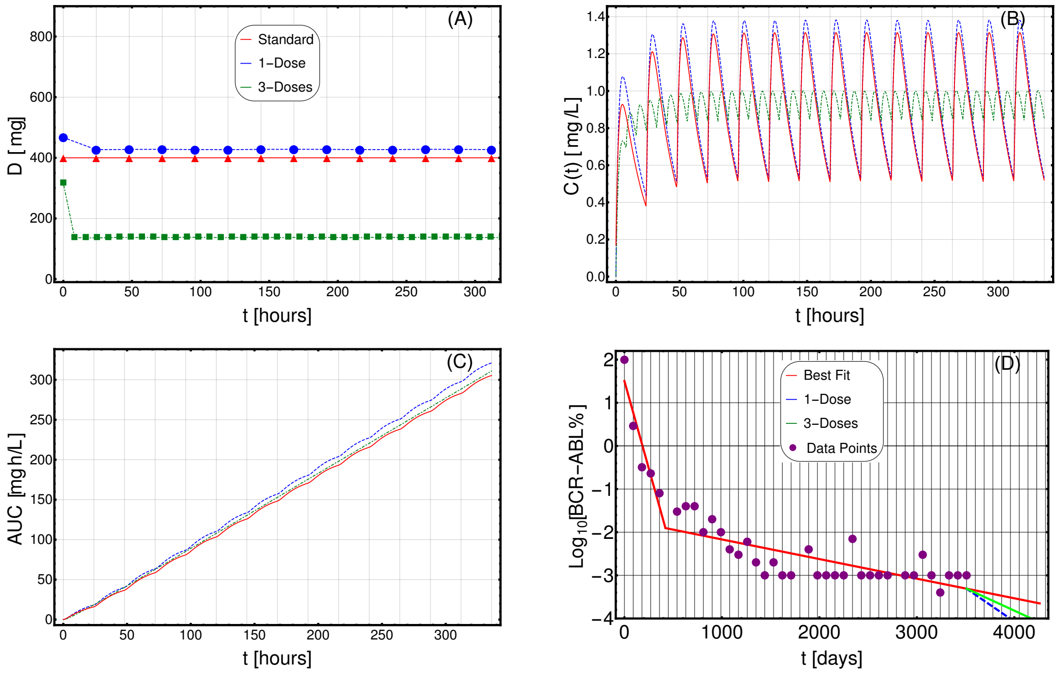


**Fig. S31. Adjusting therapy for tumor burden minimization.** Imatinib administration optimized for tumor burden minimization in patient 0001 00006 GMC (male) from (20), in the cases of: 1-dose/day (purple) and 3-doses/day (green), with $\phi = 60$, as compared to standard administration (red). In this case, the optimization is obtained on patient-specific PK and PD models. **(A)** Imatinib scheduled dosage in $mg$ (y axis), displayed on $14$ days (x axis). **(B)** Imatinib concentration in blood in $[mg/L]$ (y axis). **(C)** Temporal variation of the AUC in $[mg \cdot h/L]$ (y axis). **(D)** Longitudinal data points on tumor burden recorded are presented in purple. The best fit is shown with red lines. The slope of the right-most line is used to determine the cancer stem cell death rate and, in turn, the patient-specific PD parameters. The blue (green) line represents the predicted cancer subpopulation decay in case the 1-dose (3-doses) optimal administration

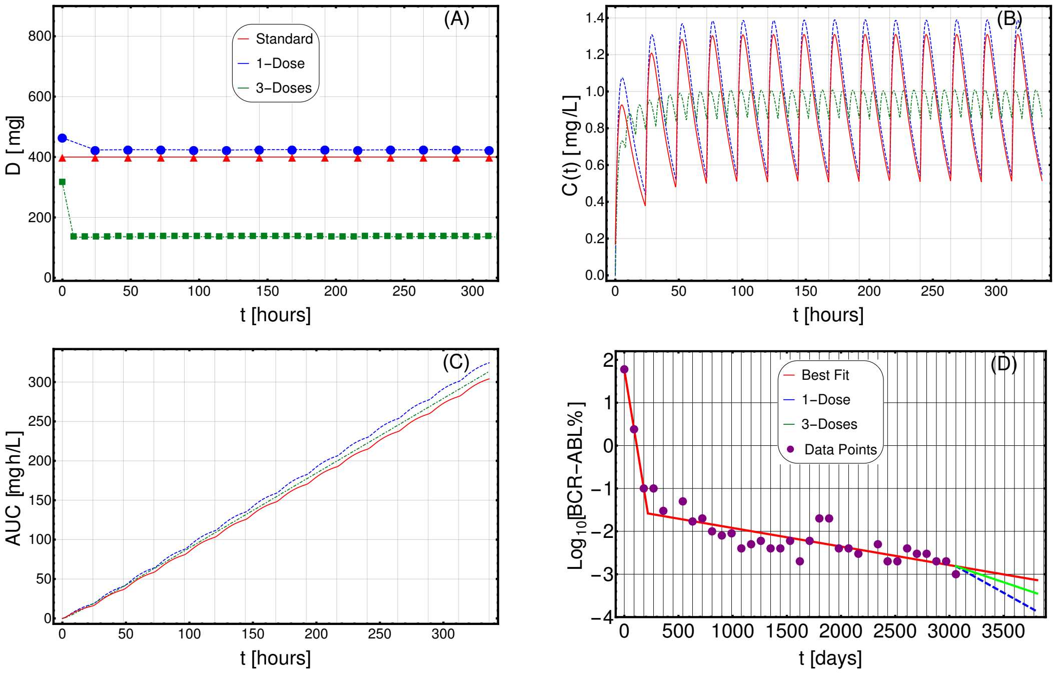


**Fig. S32. Adjusting therapy for tumor burden minimization.** Imatinib administration optimized for tumor burden minimization in patient 0001 00009 MJG (male) from (20), in the cases of: 1-dose/day (purple) and 3-doses/day (green), with $\phi = 60$, as compared to standard administration (red). In this case, the optimization is obtained on patient-specific PK and PD models. **(A)** Imatinib scheduled dosage in $mg$ (y axis), displayed on $14$ days (x axis). **(B)** Imatinib concentration in blood in $[mg/L]$ (y axis). **(C)** Temporal variation of the AUC in $[mg \cdot h/L]$ (y axis). **(D)** Longitudinal data points on tumor burden recorded are presented in purple. The best fit is shown with red lines. The slope of the right-most line is used to determine the cancer stem cell death rate and, in turn, the patient-specific PD parameters. The blue (green) line represents the predicted cancer subpopulation decay in case the 1-dose (3-doses) optimal administration

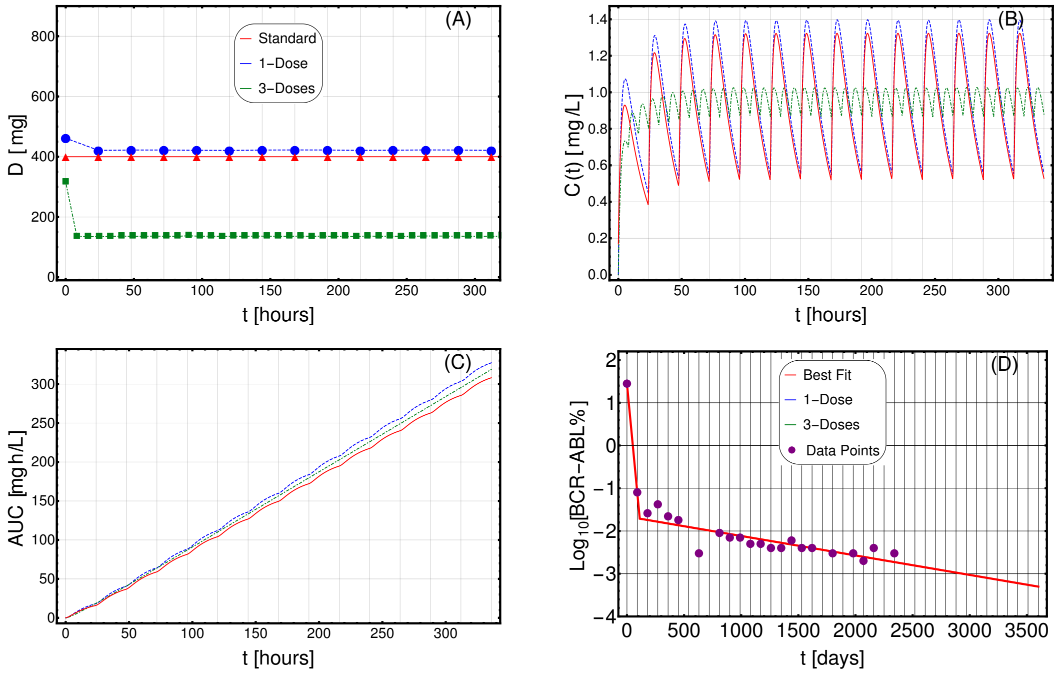


**Fig. S33. Adjusting therapy for tumor burden minimization.** Imatinib administration optimized for tumor burden minimization in patient 0002 00003 SMV (male) from (20), in the cases of: 1-dose/day (purple) and 3-doses/day (green), with $\phi = 60$, as compared to standard administration (red). In this case, the optimization is obtained on patient-specific PK and PD models. **(A)** Imatinib scheduled dosage in $mg$ (y axis), displayed on $14$ days (x axis). **(B)** Imatinib concentration in blood in $[mg/L]$ (y axis). **(C)** Temporal variation of the AUC in $[mg \cdot h/L]$ (y axis). **(D)** Longitudinal data points on tumor burden recorded are presented in purple. The best fit is shown with red lines. The slope of the right-most line is used to determine the cancer stem cell death rate and, in turn, the patient-specific PD parameters. The blue (green) line represents the predicted cancer subpopulation decay in case the 1-dose (3-doses) optimal administration

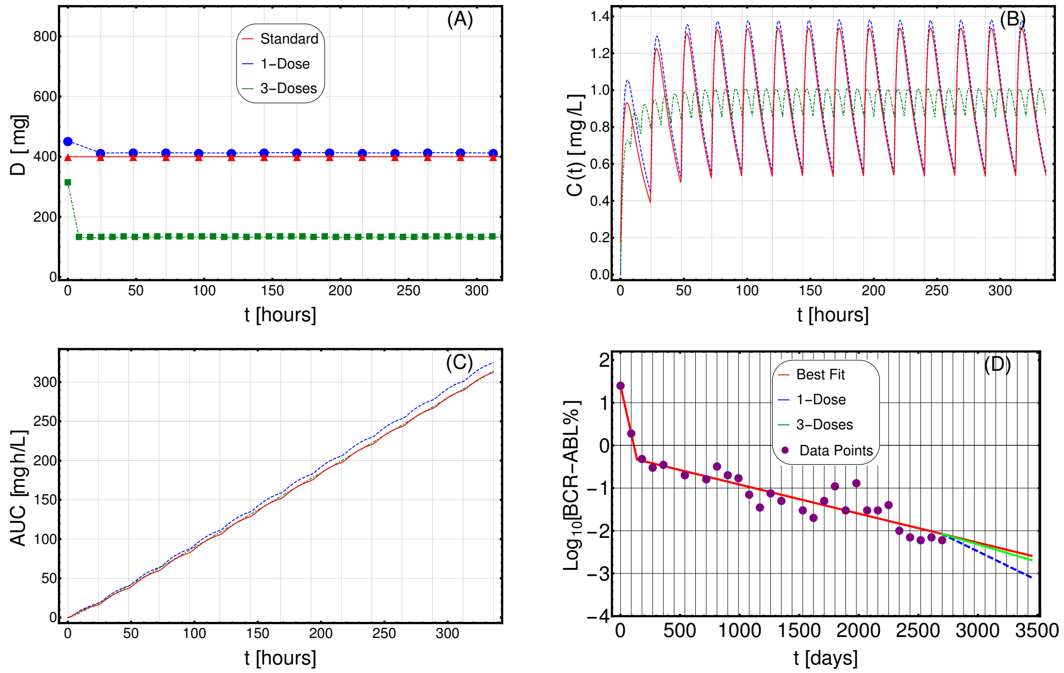


**Fig. S34. Adjusting therapy for tumor burden minimization.** Imatinib administration optimized for tumor burden minimization in patient 0002 00007 PMG (male) from (20), in the cases of: 1-dose/day (purple) and 3-doses/day (green), with $\phi = 60$, as compared to standard administration (red). In this case, the optimization is obtained on patient-specific PK and PD models. **(A)** Imatinib scheduled dosage in $mg$ (y axis), displayed on $14$ days (x axis). **(B)** Imatinib concentration in blood in $[mg/L]$ (y axis). **(C)** Temporal variation of the AUC in $[mg \cdot h/L]$ (y axis). **(D)** Longitudinal data points on tumor burden recorded are presented in purple. The best fit is shown with red lines. The slope of the right-most line is used to determine the cancer stem cell death rate and, in turn, the patient-specific PD parameters. The blue (green) line represents the predicted cancer subpopulation decay in case the 1-dose (3-doses) optimal administration

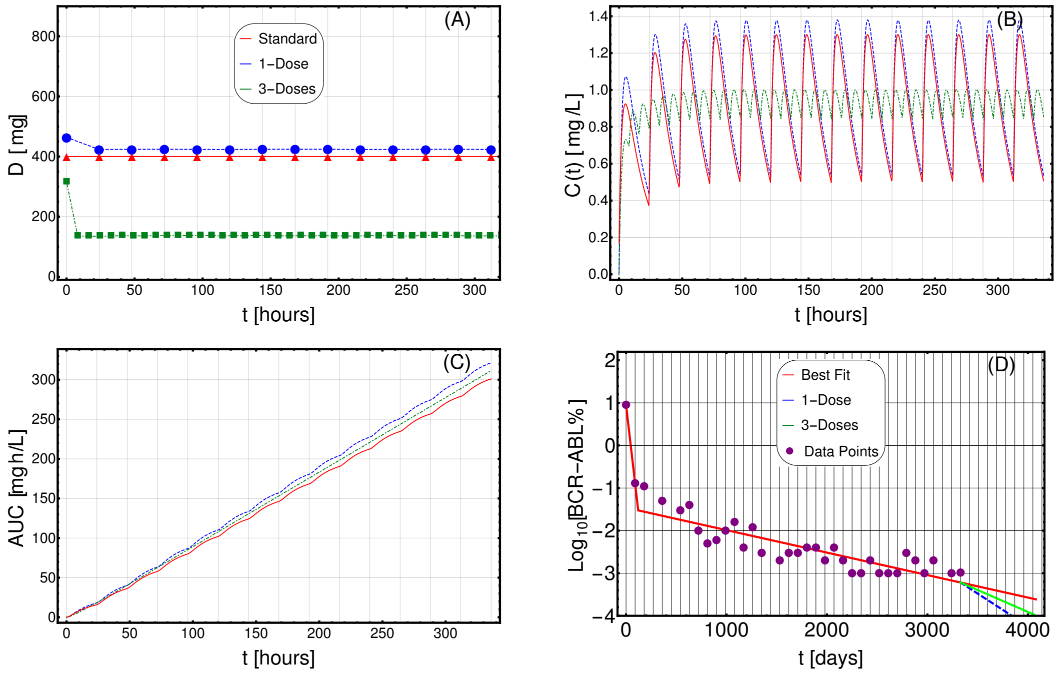


**Fig. S35. Adjusting therapy for tumor burden minimization.** Imatinib administration optimized for tumor burden minimization in patient 0002 00008 SAT (male) from (20), in the cases of: 1-dose/day (purple) and 3-doses/day (green), with $\phi = 60$, as compared to standard administration (red). In this case, the optimization is obtained on patient-specific PK and PD models. **(A)** Imatinib scheduled dosage in $mg$ (y axis), displayed on $14$ days (x axis). **(B)** Imatinib concentration in blood in $[mg/L]$ (y axis). **(C)** Temporal variation of the AUC in $[mg \cdot h/L]$ (y axis). **(D)** Longitudinal data points on tumor burden recorded are presented in purple. The best fit is shown with red lines. The slope of the right-most line is used to determine the cancer stem cell death rate and, in turn, the patient-specific PD parameters. The blue (green) line represents the predicted cancer subpopulation decay in case the 1-dose (3-doses) optimal administration

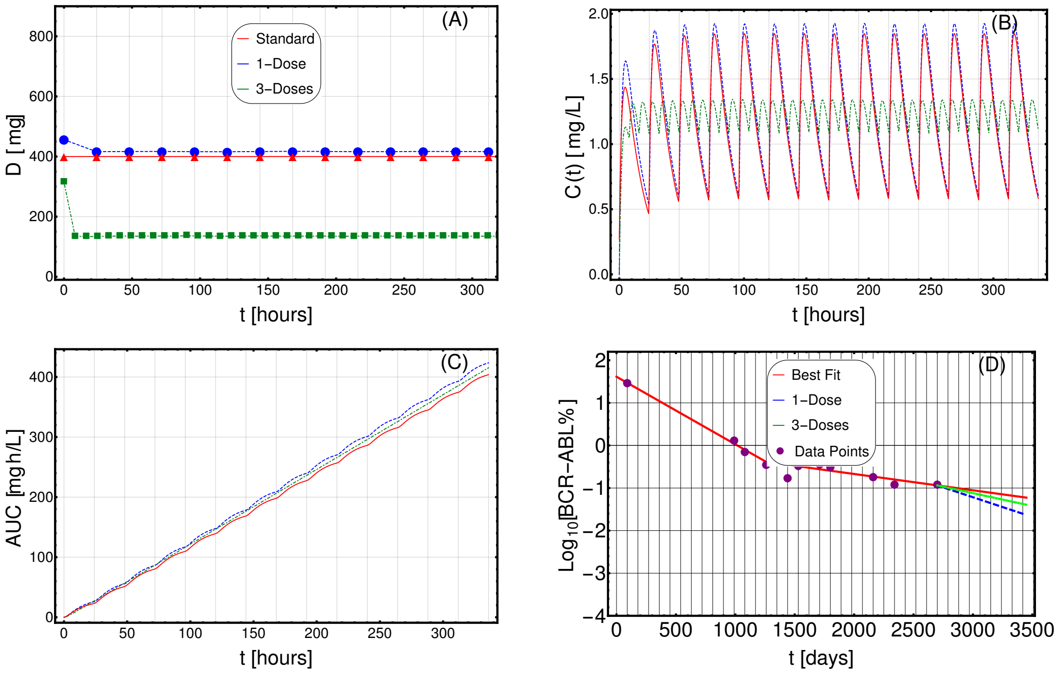


**Fig. S36. Adjusting therapy for tumor burden minimization.** Imatinib administration optimized for tumor burden minimization in patient 0002 00008 CL (female) from (20), in the cases of: 1-dose/day (purple) and 3-doses/day (green), with $\phi = 75$, as compared to standard administration (red). In this case, the optimization is obtained on patient-specific PK and PD models. **(A)** Imatinib scheduled dosage in $mg$ (y axis), displayed on $14$ days (x axis). **(B)** Imatinib concentration in blood in $[mg/L]$ (y axis). **(C)** Temporal variation of the AUC in $[mg \cdot h/L]$ (y axis). **(D)** Longitudinal data points on tumor burden recorded are presented in purple. The best fit is shown with red lines. The slope of the right-most line is used to determine the cancer stem cell death rate and, in turn, the patient-specific PD parameters. The blue (green) line represents the predicted cancer subpopulation decay in case the 1-dose (3-doses) optimal administration

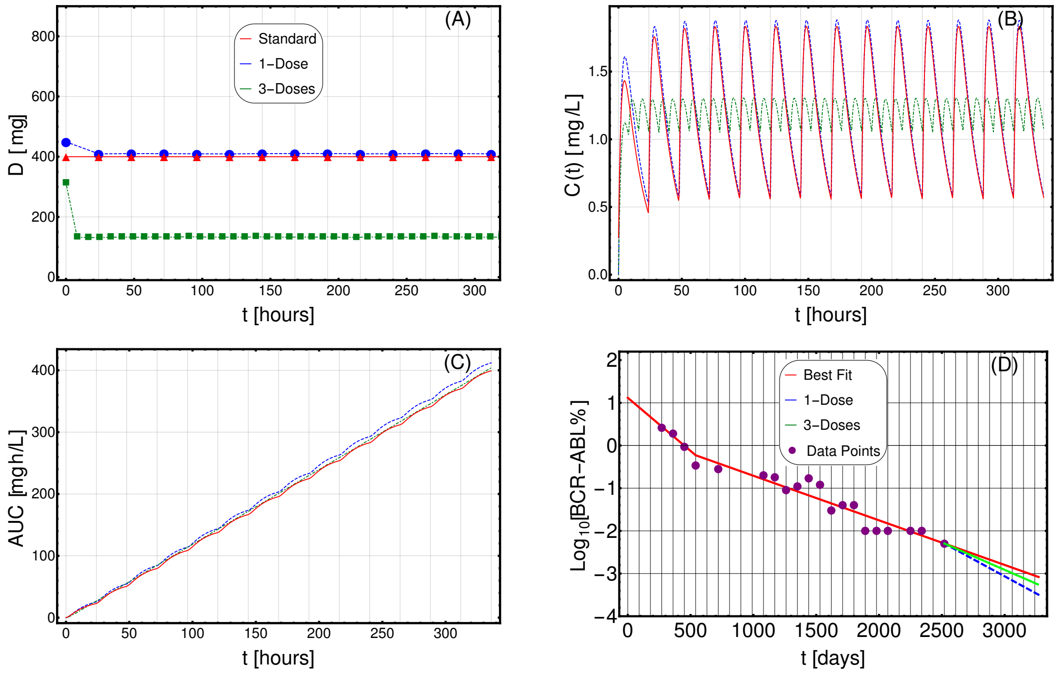


**Fig. S37. Adjusting therapy for tumor burden minimization.** Imatinib administration optimized for tumor burden minimization in patient 0004 00003 CAR (male) from (20), in the cases of: 1-dose/day (purple) and 3-doses/day (green), with $\phi = 75$, as compared to standard administration (red). In this case, the optimization is obtained on patient-specific PK and PD models. **(A)** Imatinib scheduled dosage in $mg$ (y axis), displayed on $14$ days (x axis). **(B)** Imatinib concentration in blood in $[mg/L]$ (y axis). **(C)** Temporal variation of the AUC in $[mg \cdot h/L]$ (y axis). **(D)** Longitudinal data points on tumor burden recorded are presented in purple. The best fit is shown with red lines. The slope of the right-most line is used to determine the cancer stem cell death rate and, in turn, the patient-specific PD parameters. The blue (green) line represents the predicted cancer subpopulation decay in case the 1-dose (3-doses) optimal administration

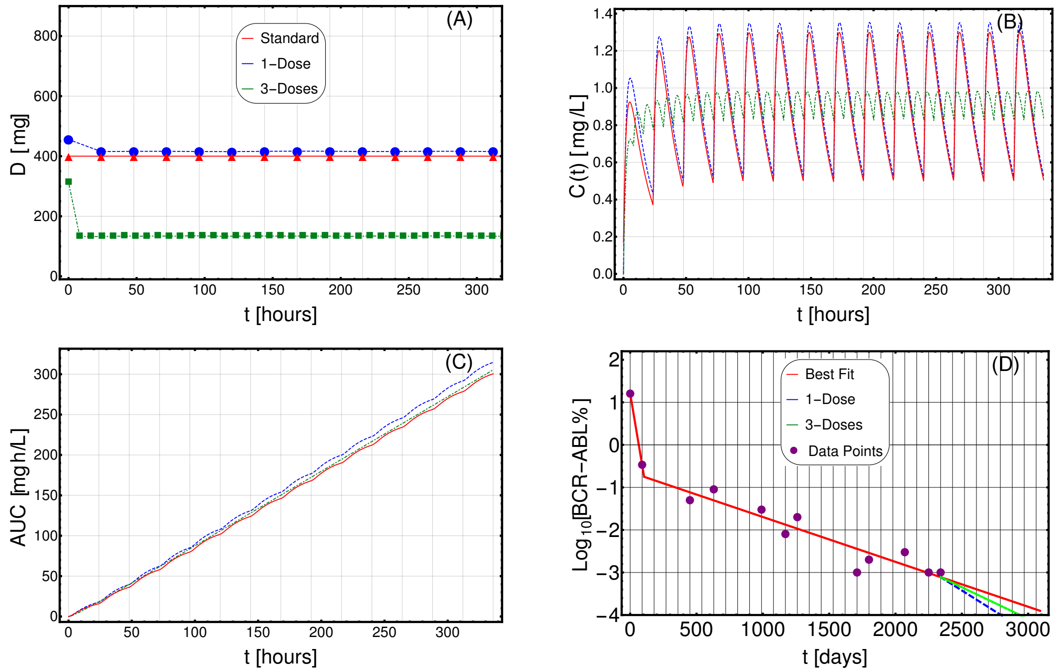


**Fig. S38. Adjusting therapy for tumor burden minimization.** Imatinib administration optimized for tumor burden minimization in patient 0004 00006 GM (male) from (20), in the cases of: 1-dose/day (purple) and 3-doses/day (green), with $\phi = 60$, as compared to standard administration (red). In this case, the optimization is obtained on patient-specific PK and PD models. **(A)** Imatinib scheduled dosage in $mg$ (y axis), displayed on $14$ days (x axis). **(B)** Imatinib concentration in blood in $[mg/L]$ (y axis). **(C)** Temporal variation of the AUC in $[mg \cdot h/L]$ (y axis). **(D)** Longitudinal data points on tumor burden recorded are presented in purple. The best fit is shown with red lines. The slope of the right-most line is used to determine the cancer stem cell death rate and, in turn, the patient-specific PD parameters. The blue (green) line represents the predicted cancer subpopulation decay in case the 1-dose (3-doses) optimal administration

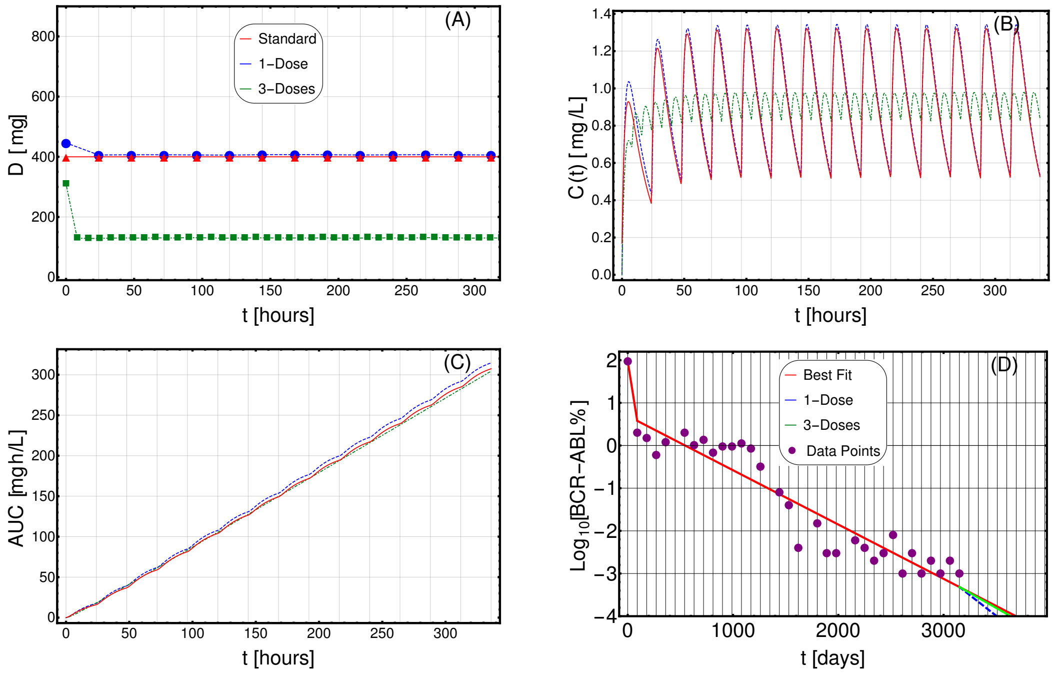


**Fig. S39. Adjusting therapy for tumor burden minimization.** Imatinib administration optimized for tumor burden minimization in patient 0006 00006 ML (male) from (20), in the cases of: 1-dose/day (purple) and 3-doses/day (green), with $\phi = 60$, as compared to standard administration (red). In this case, the optimization is obtained on patient-specific PK and PD models. **(A)** Imatinib scheduled dosage in $mg$ (y axis), displayed on $14$ days (x axis). **(B)** Imatinib concentration in blood in $[mg/L]$ (y axis). **(C)** Temporal variation of the AUC in $[mg \cdot h/L]$ (y axis). **(D)** Longitudinal data points on tumor burden recorded are presented in purple. The best fit is shown with red lines. The slope of the right-most line is used to determine the cancer stem cell death rate and, in turn, the patient-specific PD parameters. The blue (green) line represents the predicted cancer subpopulation decay in case the 1-dose (3-doses) optimal administration

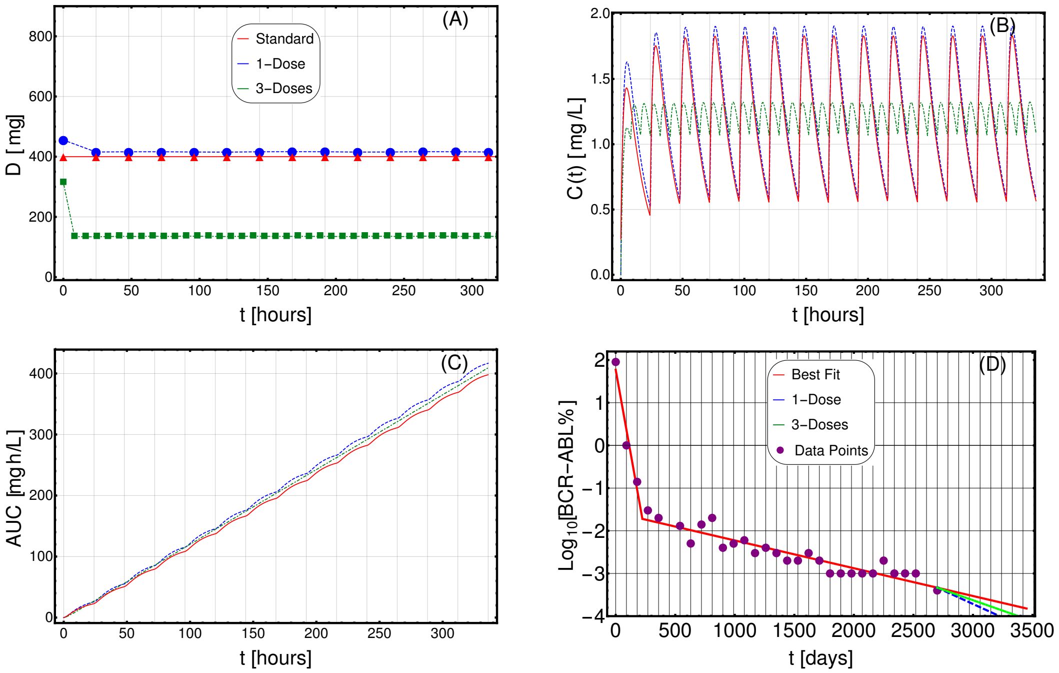


**Fig. S40. Adjusting therapy for tumor burden minimization.** Imatinib administration optimized for tumor burden minimization in patient 0006 00007 RJW (female) from (20), in the cases of: 1-dose/day (purple) and 3-doses/day (green), with $\phi = 75$, as compared to standard administration (red). In this case, the optimization is obtained on patient-specific PK and PD models. **(A)** Imatinib scheduled dosage in $mg$ (y axis), displayed on $14$ days (x axis). **(B)** Imatinib concentration in blood in $[mg/L]$ (y axis). **(C)** Temporal variation of the AUC in $[mg \cdot h/L]$ (y axis). **(D)** Longitudinal data points on tumor burden recorded are presented in purple. The best fit is shown with red lines. The slope of the right-most line is used to determine the cancer stem cell death rate and, in turn, the patient-specific PD parameters. The blue (green) line represents the predicted cancer subpopulation decay in case the 1-dose (3-doses) optimal administration

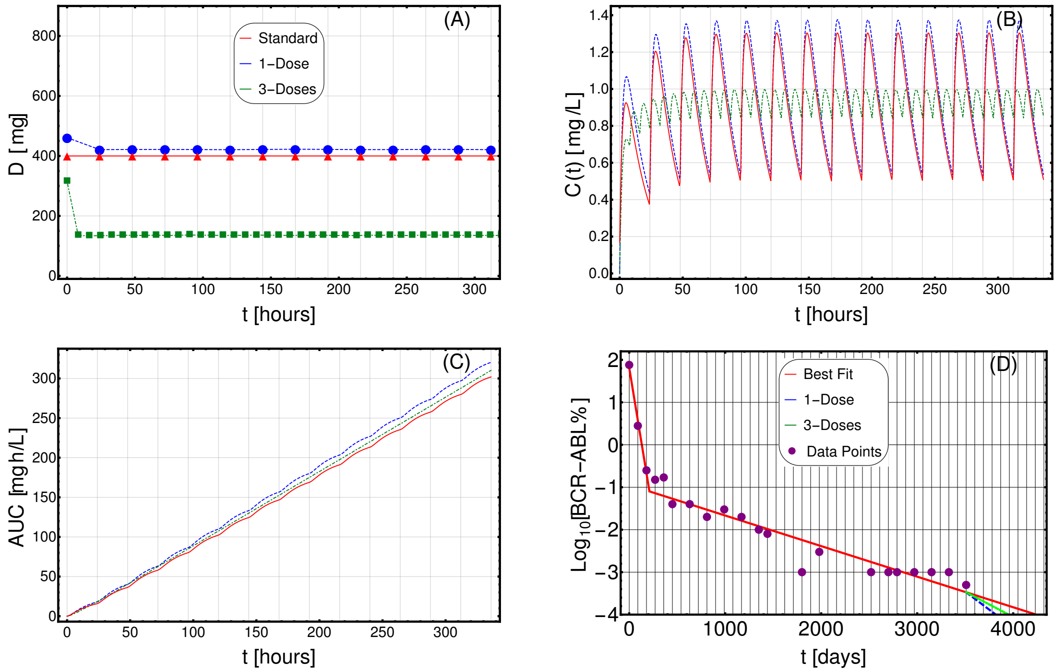


**Fig. S41. Adjusting therapy for tumor burden minimization.** Imatinib administration optimized for tumor burden minimization in patient 0007 00001 AJV (male) from (20), in the cases of: 1-dose/day (purple) and 3-doses/day (green), with $\phi = 60$, as compared to standard administration (red). In this case, the optimization is obtained on patient-specific PK and PD models. **(A)** Imatinib scheduled dosage in $mg$ (y axis), displayed on $14$ days (x axis). **(B)** Imatinib concentration in blood in $[mg/L]$ (y axis). **(C)** Temporal variation of the AUC in $[mg \cdot h/L]$ (y axis). **(D)** Longitudinal data points on tumor burden recorded are presented in purple. The best fit is shown with red lines. The slope of the right-most line is used to determine the cancer stem cell death rate and, in turn, the patient-specific PD parameters. The blue (green) line represents the predicted cancer subpopulation decay in case the 1-dose (3-doses) optimal administration

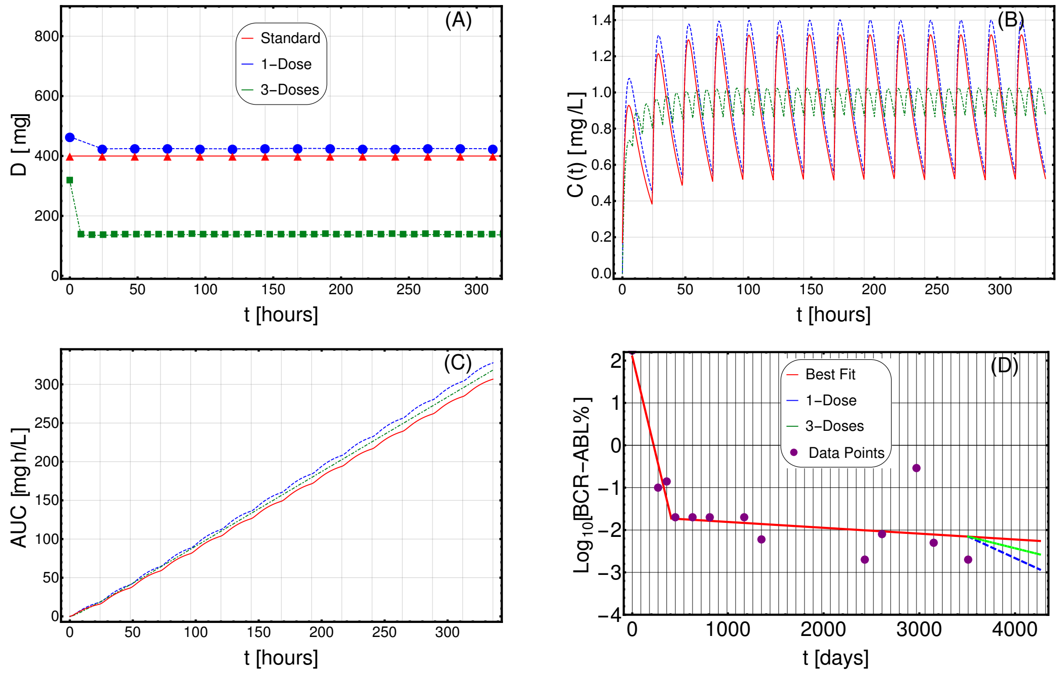


**Fig. S42. Adjusting therapy for tumor burden minimization.** Imatinib administration optimized for tumor burden minimization in patient 0007 00002 DPS (male) from (20), in the cases of: 1-dose/day (purple) and 3-doses/day (green), with $\phi = 60$, as compared to standard administration (red). In this case, the optimization is obtained on patient-specific PK and PD models. **(A)** Imatinib scheduled dosage in $mg$ (y axis), displayed on $14$ days (x axis). **(B)** Imatinib concentration in blood in $[mg/L]$ (y axis). **(C)** Temporal variation of the AUC in $[mg \cdot h/L]$ (y axis). **(D)** Longitudinal data points on tumor burden recorded are presented in purple. The best fit is shown with red lines. The slope of the right-most line is used to determine the cancer stem cell death rate and, in turn, the patient-specific PD parameters. The blue (green) line represents the predicted cancer subpopulation decay in case the 1-dose (3-doses) optimal administration

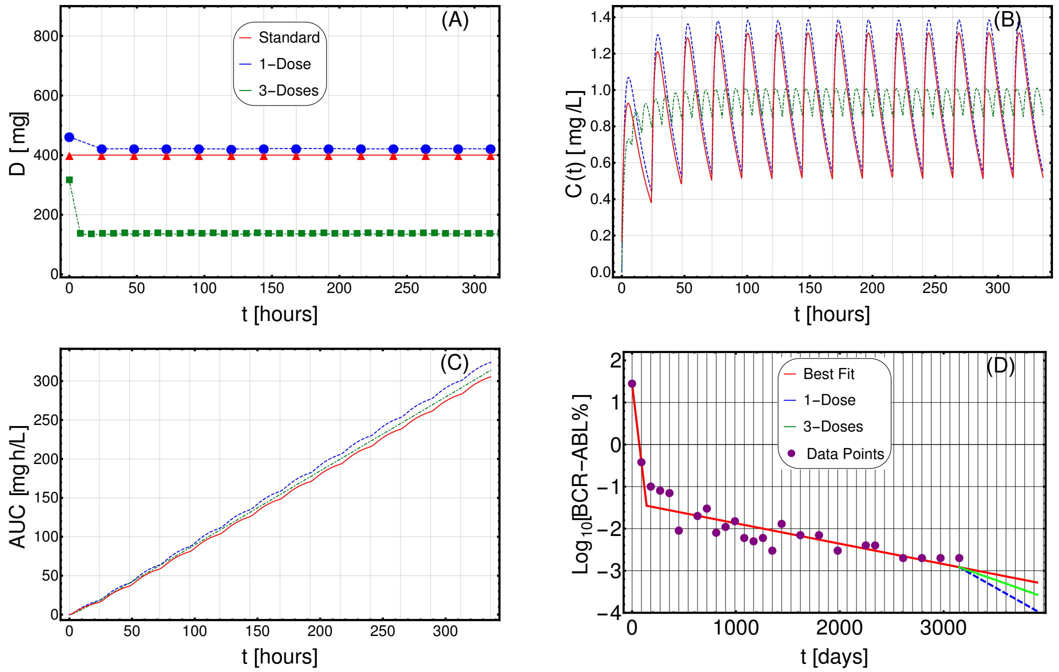


**Fig. S43. Adjusting therapy for tumor burden minimization.** Imatinib administration optimized for tumor burden minimization in patient 0008 00002 PWR (male) from (20), in the cases of: 1-dose/day (purple) and 3-doses/day (green), with $\phi = 60$, as compared to standard administration (red). In this case, the optimization is obtained on patient-specific PK and PD models. **(A)** Imatinib scheduled dosage in $mg$ (y axis), displayed on $14$ days (x axis). **(B)** Imatinib concentration in blood in $[mg/L]$ (y axis). **(C)** Temporal variation of the AUC in $[mg \cdot h/L]$ (y axis). **(D)** Longitudinal data points on tumor burden recorded are presented in purple. The best fit is shown with red lines. The slope of the right-most line is used to determine the cancer stem cell death rate and, in turn, the patient-specific PD parameters. The blue (green) line represents the predicted cancer subpopulation decay in case the 1-dose (3-doses) optimal administration

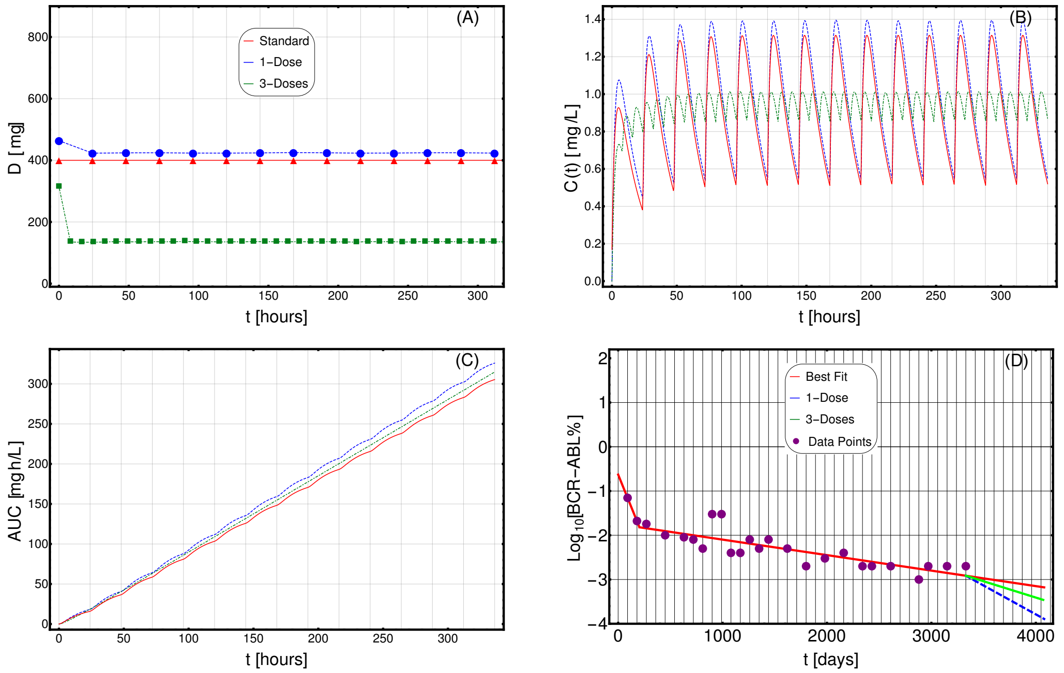


**Fig. S44. Adjusting therapy for tumor burden minimization.** Imatinib administration optimized for tumor burden minimization in patient 0008 00003 LJG (male) from (20), in the cases of: 1-dose/day (purple) and 3-doses/day (green), with $\phi = 60$, as compared to standard administration (red). In this case, the optimization is obtained on patient-specific PK and PD models. **(A)** Imatinib scheduled dosage in $mg$ (y axis), displayed on $14$ days (x axis). **(B)** Imatinib concentration in blood in $[mg/L]$ (y axis). **(C)** Temporal variation of the AUC in $[mg \cdot h/L]$ (y axis). **(D)** Longitudinal data points on tumor burden recorded are presented in purple. The best fit is shown with red lines. The slope of the right-most line is used to determine the cancer stem cell death rate and, in turn, the patient-specific PD parameters. The blue (green) line represents the predicted cancer subpopulation decay in case the 1-dose (3-doses) optimal administration

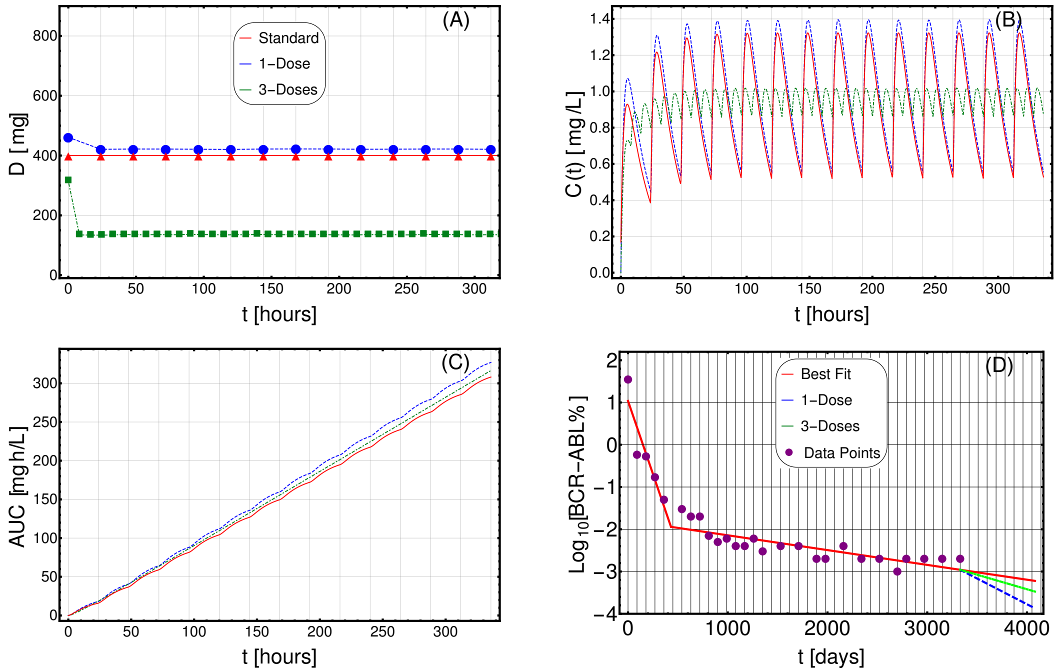


**Fig. S45. Adjusting therapy for tumor burden minimization.** Imatinib administration optimized for tumor burden minimization in patient 0008 00005 BAT (male) from (20), in the cases of: 1-dose/day (purple) and 3-doses/day (green), with $\phi = 60$, as compared to standard administration (red). In this case, the optimization is obtained on patient-specific PK and PD models. **(A)** Imatinib scheduled dosage in $mg$ (y axis), displayed on $14$ days (x axis). **(B)** Imatinib concentration in blood in $[mg/L]$ (y axis). **(C)** Temporal variation of the AUC in $[mg \cdot h/L]$ (y axis). **(D)** Longitudinal data points on tumor burden recorded are presented in purple. The best fit is shown with red lines. The slope of the right-most line is used to determine the cancer stem cell death rate and, in turn, the patient-specific PD parameters. The blue (green) line represents the predicted cancer subpopulation decay in case the 1-dose (3-doses) optimal administration

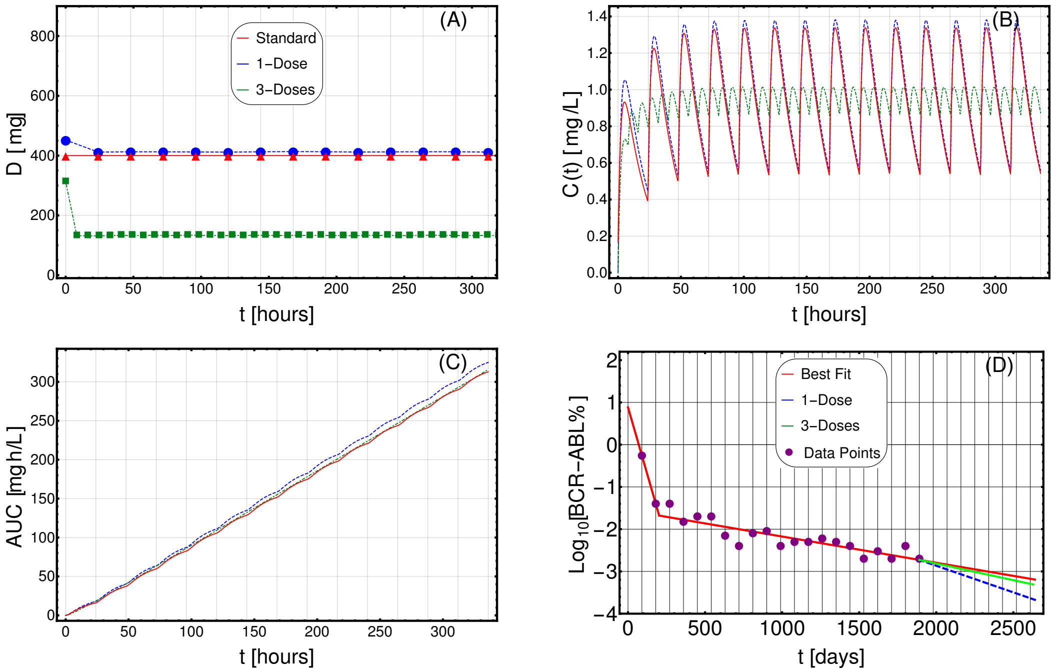


**Fig. S46. Adjusting therapy for tumor burden minimization.** Imatinib administration optimized for tumor burden minimization in patient 0009 00003 LH (male) from (20), in the cases of: 1-dose/day (purple) and 3-doses/day (green), with $\phi = 60$, as compared to standard administration (red). In this case, the optimization is obtained on patient-specific PK and PD models. **(A)** Imatinib scheduled dosage in $mg$ (y axis), displayed on $14$ days (x axis). **(B)** Imatinib concentration in blood in $[mg/L]$ (y axis). **(C)** Temporal variation of the AUC in $[mg \cdot h/L]$ (y axis). **(D)** Longitudinal data points on tumor burden recorded are presented in purple. The best fit is shown with red lines. The slope of the right-most line is used to determine the cancer stem cell death rate and, in turn, the patient-specific PD parameters. The blue (green) line represents the predicted cancer subpopulation decay in case the 1-dose (3-doses) optimal administration

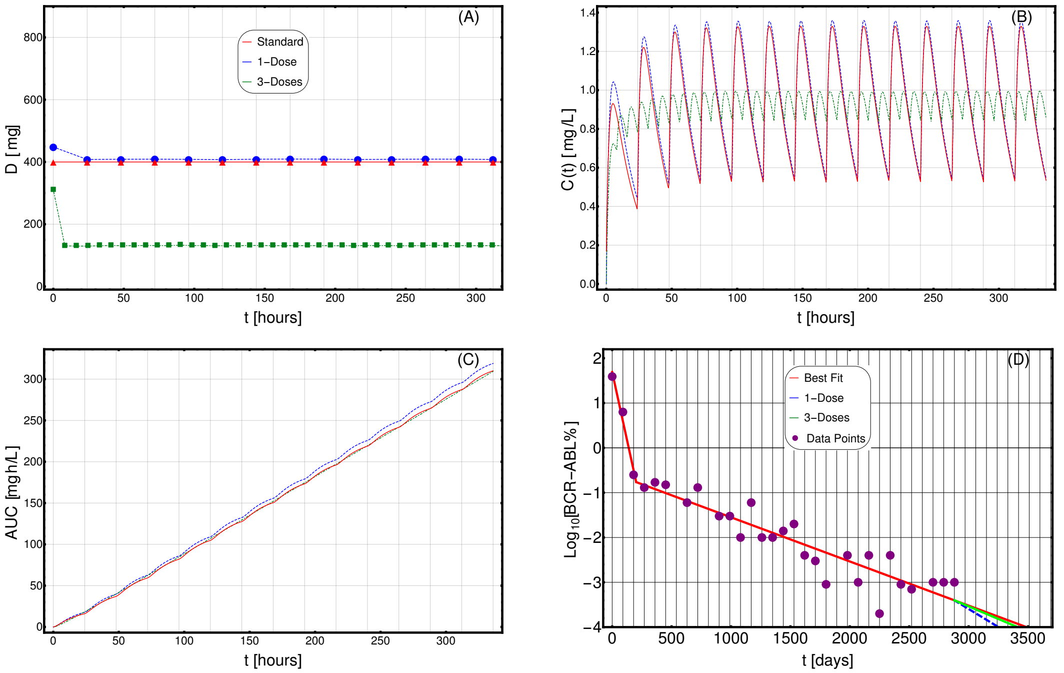


**Fig. S47. Adjusting therapy for tumor burden minimization.** Imatinib administration optimized for tumor burden minimization in patient 0010 00001 HJ (male) from (20), in the cases of: 1-dose/day (purple) and 3-doses/day (green), with $\phi = 60$, as compared to standard administration (red). In this case, the optimization is obtained on patient-specific PK and PD models. **(A)** Imatinib scheduled dosage in $mg$ (y axis), displayed on $14$ days (x axis). **(B)** Imatinib concentration in blood in $[mg/L]$ (y axis). **(C)** Temporal variation of the AUC in $[mg \cdot h/L]$ (y axis). **(D)** Longitudinal data points on tumor burden recorded are presented in purple. The best fit is shown with red lines. The slope of the right-most line is used to determine the cancer stem cell death rate and, in turn, the patient-specific PD parameters. The blue (green) line represents the predicted cancer subpopulation decay in case the 1-dose (3-doses) optimal administration

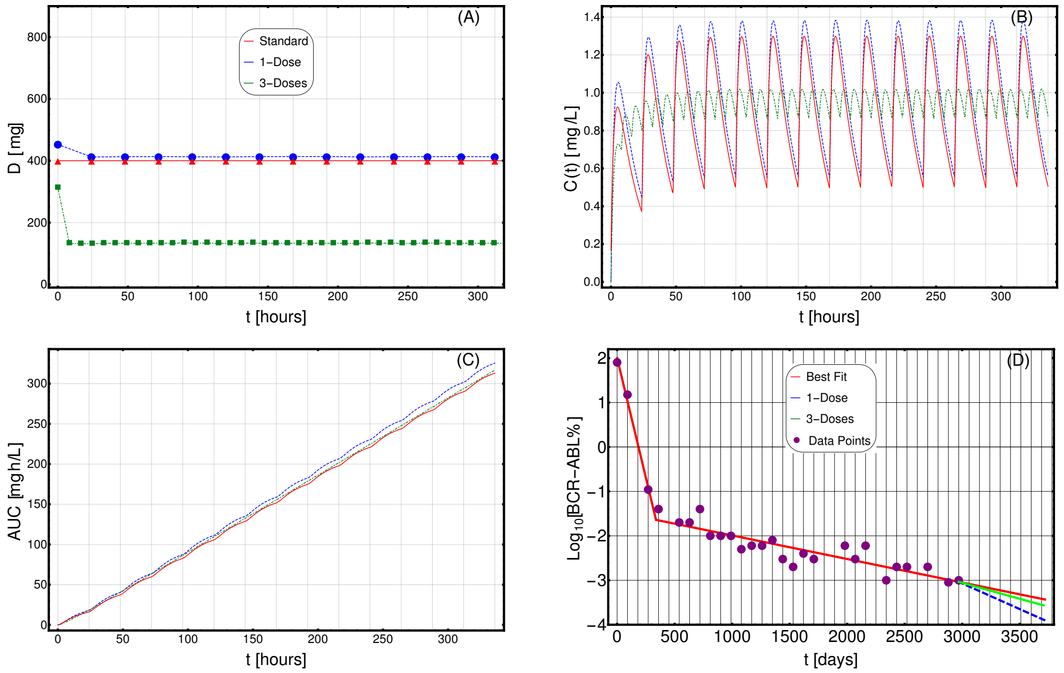


**Fig. S48. Adjusting therapy for tumor burden minimization.** Imatinib administration optimized for tumor burden minimization in patient 0010 00002 GR (male) from (20), in the cases of: 1-dose/day (purple) and 3-doses/day (green), with $\phi = 60$, as compared to standard administration (red). In this case, the optimization is obtained on patient-specific PK and PD models. **(A)** Imatinib scheduled dosage in $mg$ (y axis), displayed on $14$ days (x axis). **(B)** Imatinib concentration in blood in $[mg/L]$ (y axis). **(C)** Temporal variation of the AUC in $[mg \cdot h/L]$ (y axis). **(D)** Longitudinal data points on tumor burden recorded are presented in purple. The best fit is shown with red lines. The slope of the right-most line is used to determine the cancer stem cell death rate and, in turn, the patient-specific PD parameters. The blue (green) line represents the predicted cancer subpopulation decay in case the 1-dose (3-doses) optimal administration

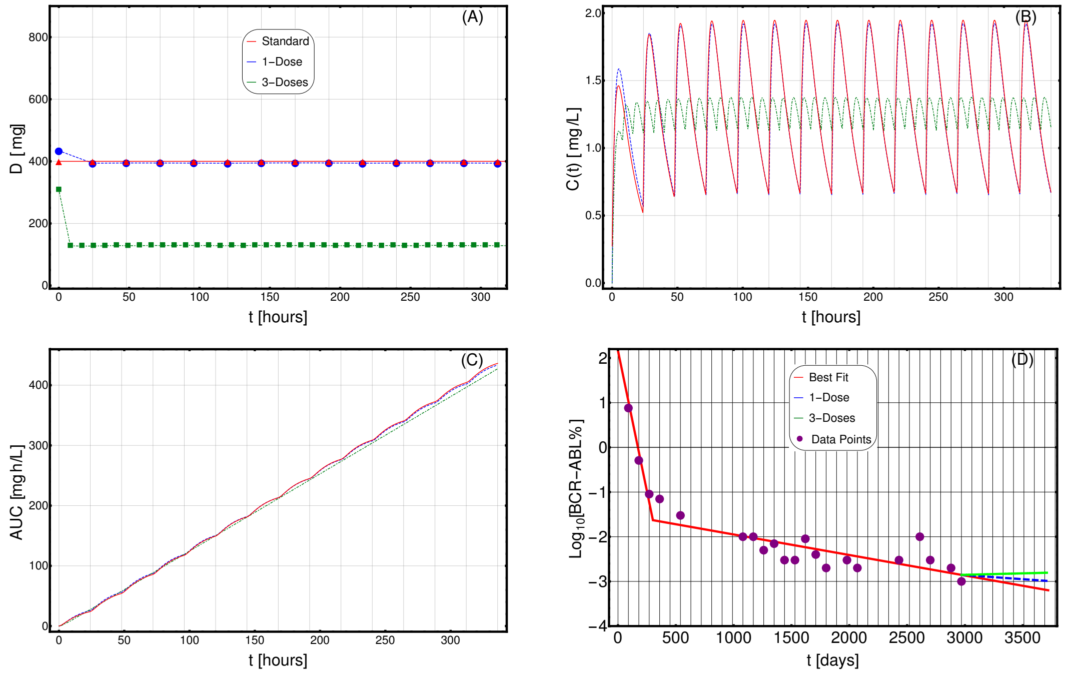


**Fig. S49. Adjusting therapy for tumor burden minimization.** Imatinib administration optimized for tumor burden minimization in patient 0011 00001 CMC (female) from (20), in the cases of: 1-dose/day (purple) and 3-doses/day (green), with $\phi = 75$, as compared to standard administration (red). In this case, the optimization is obtained on patient-specific PK and PD models. **(A)** Imatinib scheduled dosage in $mg$ (y axis), displayed on $14$ days (x axis). **(B)** Imatinib concentration in blood in $[mg/L]$ (y axis). **(C)** Temporal variation of the AUC in $[mg \cdot h/L]$ (y axis). **(D)** Longitudinal data points on tumor burden recorded are presented in purple. The best fit is shown with red lines. The slope of the right-most line is used to determine the cancer stem cell death rate and, in turn, the patient-specific PD parameters. The blue (green) line represents the predicted cancer subpopulation decay in case the 1-dose (3-doses) optimal administration

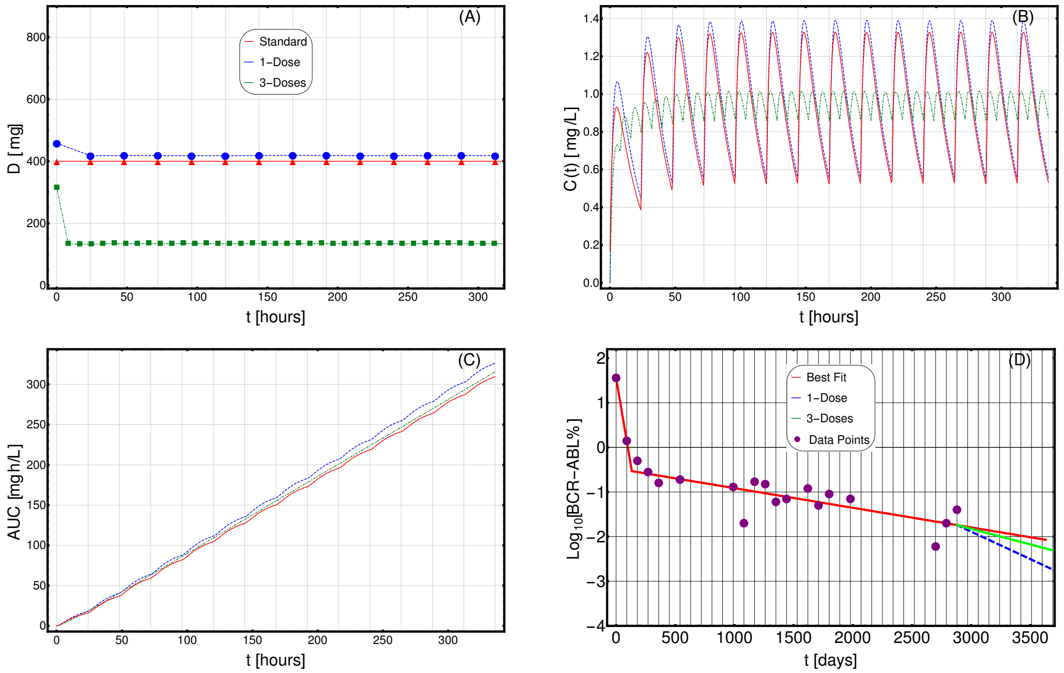


**Fig. S50. Adjusting therapy for tumor burden minimization.** Imatinib administration optimized for tumor burden minimization in patient 0011 00005 SCW (male) from (20), in the cases of: 1-dose/day (purple) and 3-doses/day (green), with $\phi = 60$, as compared to standard administration (red). In this case, the optimization is obtained on patient-specific PK and PD models. **(A)** Imatinib scheduled dosage in $mg$ (y axis), displayed on $14$ days (x axis). **(B)** Imatinib concentration in blood in $[mg/L]$ (y axis). **(C)** Temporal variation of the AUC in $[mg \cdot h/L]$ (y axis). **(D)** Longitudinal data points on tumor burden recorded are presented in purple. The best fit is shown with red lines. The slope of the right-most line is used to determine the cancer stem cell death rate and, in turn, the patient-specific PD parameters. The blue (green) line represents the predicted cancer subpopulation decay in case the 1-dose (3-doses) optimal administration

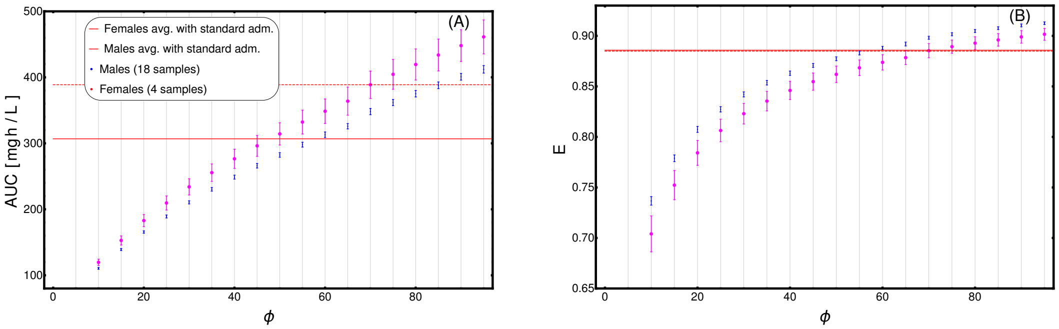


**Fig. S51. Assessment of term weights in cost function definition.** The definition of the cost function for the adjusting treatment scenario requires to set the weights of the different terms. We here considered two terms, in order to: $(i)$ minimize the tumor burden (weight $W_1$), and $(ii)$ minimize the AUC (weight $W_2$) (see Materials and Methods for further details). We scanned the values of $\phi = \frac{W_1}{W_2}$ in the range $[10, 100]$, by repeatedly applying CT4TD to the $22$-patients CML dataset from (20). **(A)**. Distribution of the value of the AUC after 14-days of the optimized therapy retrieved by CT4TD (1-dose case), for distinct values of $\phi$, with respect to the $22$ samples in the datasets, divided in males (blue) and females (pink), and compared to the average AUC values returned by standard administration (400 mg Imatinib/day) in males (red solid line) and females (red dashed line). **(B)** Distribution of efficiency computed via Eq. (9) on the time-average concentration over 14 days of the optimized therapy retrieved by CT4TD (3-dose case), for distinct values of $\phi$, and compared to the average efficiency in the standard administration scenario (solid and dashed red lines overlap).

**Table S1. Average parameters of PK model of Imatinib from (21).**

| Parameters | Value | Standard error | Units of measurement |
|---|---|---|---|
| $k_a$ | $0.61$ | $30\%$ | $h^{-1}$ |
| $CL$ | $14.3$ | $7.1\%$ | $Lh^{-1}$ |
| $v$ | $347$ | $17.9\%$ | $L$ |
| $B\bar{W}$ | $70$ | Na | $Kg$ |
| $A\bar{G}E$ | $50$ | Na | $Years$ |

**Table S2. Summary of the demographic population PK parameters for Imatinib from (21).**

| Parameter | Value |
|---|---|
| $k_a$ | $0.437$ |
| $\theta_a$ | $12.8$ |
| $\theta_b$ | $258$ |
| $\theta_1$ | $12.7$ |
| $\theta_2$ | $0.8$ |
| $\theta_3$ | $-2.1$ |
| $\theta_4$ | $61.0$ |

**Table S3. Summary data analysis.**

| Code | $\alpha \quad [days]^{-1}$ | $\beta \quad [days]^{-1}$ | $T \quad [days]$ | $R^2$ | Index |
|---|---|---|---|---|---|
| 0001 00002 RH | -0.00793675 | -0.000430922 | 453 | 0.992329 | A |
| 0001 00004 AJR | -0.0335431 | -0.000455828 | 112 | 0.99256 | B |
| 0001 00006 GMC | -0.0080923 | -0.000454651 | 421 | 0.979851 | C |
| 0001 00009 MJG | -0.0153962 | -0.000432947 | 218 | 0.985078 | D |
| 0002 00003 SMV | -0.000734153 | -0.000223423 | 1710 | 0.992211 | E |
| 0002 00007 PMG | -0.0124442 | -0.000680913 | 139 | 0.966844 | F |
| 0002 00008 SAT | -0.0204914 | -0.000527755 | 121 | 0.987923 | G |
| 0003 00002 CL | -0.00158763 | -0.000382959 | 1260 | 0.969166 | H |
| 0004 00003 CAR | -0.00249199 | -0.00104476 | 540 | 0.980976 | I |
| 0004 00006 GM | -0.0186312 | -0.00105491 | 105 | 0.987429 | J |
| 0006 00006 ML | -0.0154954 | -0.00127265 | 90 | 0.949279 | K |
| 0006 00007 RJW | -0.01559 | -0.000650637 | 224 | 0.994058 | L |
| 0007 00001 AJV | -0.0138184 | -0.000720104 | 211 | 0.987969 | M |
| 0007 00002 DPS | -0.00937013 | -0.000137684 | 408 | 0.918741 | N |
| 0008 00002 PWR | -0.0207251 | -0.000485452 | 140 | 0.985233 | O |
| 0008 00003 LJG | -0.00583334 | -0.000350359 | 204 | 0.991091 | P |
| 0008 00005 BAT | -0.00691467 | -0.000350271 | 431 | 0.987516 | Q |
| 0009 00003 LH | -0.0126738 | -0.000620824 | 202 | 0.994167 | R |
| 0010 00001 HJ | -0.0121572 | -0.000984917 | 202 | 0.977477 | S |
| 0010 00002 GR | -0.0107683 | -0.000530116 | 338 | 0.992 | T |
| 0011 00001 CMC | -0.0125545 | -0.000459044 | 302 | 0.985232 | U |
| 0011 00005 SCW | -0.0157079 | -0.000440187 | 133 | 0.948588 | V |

**Table S4. PK personal parameters. Weights are taken from (22)**

| Code | Sex | Age [Years] | Weight [Kg] | $K_a$ | $CL$ | $V$ | $EC_{50}$ $[mg/L]$ |
|---|---|---|---|---|---|---|---|
| 0001 00002 RH | M | 65 | 88.6 | 0.437 | 17.3446 | 319 | 0.121311 |
| 0001 00004 AJR | M | 55 | 88.8 | 0.437 | 17.8009 | 319 | 0.118284 |
| 0001 00006 GMC | M | 35 | 86.6 | 0.437 | 18.2417 | 319 | 0.11573 |
| 0001 00009 MJG | M | 53 | 88.8 | 0.437 | 17.8849 | 319 | 0.118019 |
| 0002 00003 SMV | M | 24 | 80.7 | 0.437 | 17.6333 | 319 | 0.121672 |
| 0002 00007 PMG | M | 64 | 88.6 | 0.437 | 17.3866 | 319 | 0.118461 |
| 0002 00008 SAT | M | 39 | 86.6 | 0.437 | 18.0737 | 319 | 0.115963 |
| 0003 00002 CL | F | 27 | 67.9 | 0.437 | 13.585 | 197 | 0.158738 |
| 0004 00003 CAR | F | 23 | 67.9 | 0.437 | 13.753 | 197 | 0.148087 |
| 0004 00006 GM | M | 49 | 89.1 | 0.437 | 18.1073 | 319 | 0.110521 |
| 0006 00006 ML | M | 58 | 88.8 | 0.437 | 17.6749 | 319 | 0.110734 |
| 0006 00007 RJW | F | 22 | 67.9 | 0.437 | 13.795 | 197 | 0.152898 |
| 0007 00001 AJV | M | 50 | 88.8 | 0.437 | 18.0109 | 319 | 0.114397 |
| 0007 00002 DPS | M | 22 | 80.7 | 0.437 | 17.7173 | 319 | 0.122035 |
| 0008 00002 PWR | M | 55 | 88.8 | 0.437 | 17.8009 | 319 | 0.117983 |
| 0008 00003 LJG | M | 55 | 88.8 | 0.437 | 17.8009 | 319 | 0.119356 |
| 0008 00005 BAT | M | 59 | 88.8 | 0.437 | 17.6329 | 319 | 0.120372 |
| 0009 00003 LH | M | 65 | 88.6 | 0.437 | 17.3446 | 319 | 0.119339 |
| 0010 00001 HJ | M | 61 | 88.6 | 0.437 | 17.5126 | 319 | 0.114586 |
| 0010 00002 GR | M | 63 | 88.6 | 0.437 | 17.3446 | 319 | 0.12028 |
| 0011 00001 CMC | F | 52 | 75.9 | 0.437 | 12.535 | 197 | 0.169956 |
| 0011 00005 SCW | M | 26 | 80.7 | 0.437 | 17.5493 | 319 | 0.119957 |

**Table S5. Results of study of robustness.** $a$ **and** $b$ **refer to parameter of the curve** $\Delta\mathcal{L}(\sigma_r)/\mathcal{L}_0 = a + b\sigma_r^2/r$ **with** $r = k_a, CL, v$**. Plot of these curves are presented in fig. S8**

| Parameter | $a$ | $b$ | $R^2$ |
|---|---|---|---|
| $CL$ | -0.0051486 | 1.99948 | 0.997 |
| $v$ | -4.81123 | 212.498 | 0.997 |
| $k_a$ | -0.0179207 | 2.7182 | 0.998 |

**Table S6. Configuration of RedCRAB**

| Number of SI | Type of basis | Number of basis used in the first iteration | Number of basis used in the others iteration | Number of time points |
|---|---|---|---|---|
| 50 | Fourier | 120 | 100 | 1001 |
